# A T2T-CHM13 recombination map and globally diverse haplotype reference panel improves phasing and imputation

**DOI:** 10.1101/2025.02.24.639687

**Authors:** Joseph L. Lalli, Andrew N. Bortvin, Rajiv C. McCoy, Donna M. Werling

## Abstract

The T2T-CHM13 complete human reference genome contains ∼200 Mb of previously unresolved sequence, improving read mapping and variant calling compared to GRCh38. However, the benefits of using complete reference genomes for phasing and imputation are unclear. Here, we present a reference T2T-CHM13 recombination map and phased haplotype panel derived from 3,202 samples from the 1000 Genomes Project (1kGP). Using published long-read based assemblies as a reference-neutral ground truth, we compared our T2T-CHM13 1kGP panel to the previously released GRCh38 1kGP panel. We found that alignment to T2T-CHM13 resulted in 38% fewer assembly-discordant SNP genotypes and 16% fewer phasing switch errors. The largest gains in panel accuracy were observed on chromosome X and in the regions flanking loci prone to disease-causing CNVs. Moreover, downsampled genomes from the Simons Genome Diversity Project attained higher imputation accuracy when using the T2T-CHM13 versus GRCh38 1kGP panel. Our study demonstrates that use of the T2T-CHM13 phased haplotype panel improves statistical phasing and imputation for samples from diverse human populations.

## Introduction

The T2T-CHM13 reference genome corrects thousands of errors in previous reference genome assemblies and resolves approximately 200 megabases (Mb) of human genetic sequence^1^. When compared to the GRCh38 reference genome, use of this telomere-to-telomere (T2T) assembly as a reference has been shown to reduce short-read alignment errors and improve variant discovery and genotyping^2–4^. Nevertheless, several practical barriers hinder the widespread adoption of T2T-CHM13 for downstream applications.

For example, statistical phasing and imputation of haplotypes are critical for genetic applications as diverse as genome-wide association studies (GWAS)^5–7^, Mendelian disease mapping^8^, admixture analyses^9^, and allele-specific studies of gene regulation^10–14^. Phasing assigns heterozygous variants to parental haplotypes, allowing for the investigation of compound heterozygosity^15^ and other haplotype-level phenotypic associations^16^.

Meanwhile, genotype imputation leverages these haplotype panels to statistically infer genotypes at variants not directly assayed, extending genomic analysis beyond genotyped markers to the potentially numerous additional variants in linkage disequilibrium (LD) among which causal mutations may occur^5^. Both methods require the use of reference haplotypes and a genome-wide map of recombination rates that represent the population in question. However, to our knowledge there is no publicly available recombination map or large haplotype reference panel generated directly from T2T-CHM13-aligned data that is suitable for phasing or imputation algorithms^17^. While GRCh38 resources can be “lifted over” to T2T-CHM13 coordinates^18^, these methods are frequently unreliable in highly variable regions of the genome, do not benefit from improved read mapping or variant calling^2,3^, and preclude the investigation of newly resolved regions^19^.

In this study, we present both population-specific and globally averaged recombination maps generated from T2T-CHM13-aligned data, along with a phased reference haplotype panel of 3,202 individuals. We evaluate the accuracy of genotyping, phasing, and imputation of our T2T-CHM13 1kGP panel compared to the previous GRCh38 1kGP panel. Both were generated by a functionally equivalent pipeline, permitting direct comparison of the two callsets^20,21^.

To facilitate these comparisons, we leveraged the near-reference quality full length assemblies of 1kGP samples from the Human Pangenome Reference Consortium (HPRC)^22^ and the Human Genome Structural Variant Consortium (HGSVC)^23^ as sources of reference-neutral ground truth variation. Importantly for our comparisons, these aligned genome assemblies do not need to be lifted over, as genetic variation in shared genomic regions can be accurately described in either GRCh38 or T2T-CHM13 coordinates, allowing us to directly measure the accuracy of short-read variant calls in complex genomic regions of diverse ancestral origin. We additionally developed LiftoverIndel, a liftover tool that accounts for differences in indel representation between reference assemblies, to assess downstream uses of these panels.

We found that the T2T-CHM13 1kGP panel contains 38% fewer assembly-discordant SNP genotypes and 16% fewer phasing switch errors than the GRCh38 panel, with the largest improvements on chromosome X and in regions flanking disease-associated CNV loci. When genotyping-derived flip errors are excluded, true phasing error rates are 50% lower in the T2T-CHM13 panel overall, and ten-fold lower on chromosome X. We also found that variants that were phased and imputed with this T2T-CHM13 panel are more accurate than those generated from the GRCh38 panel. Both the T2T-CHM13 recombination map and 1kGP T2T-CHM13 panel are publicly available.

## Results

### LD-based recombination maps from 1000 Genomes Project data aligned to T2T-CHM13

Our panel builds upon data from Rhie et al.^24^, which produced an unphased T2T-CHM13 1kGP callset using the functionally equivalent GATK pipeline^21^. To reduce false positive variant calls that may interfere with haplotype inference, we applied strict variant quality controls, including filtering out variants with a variant quality score log-odds (VQSLOD^25^) of less than zero (see Methods). Our VQSLOD filter resulted in the exclusion of an additional 14,491,949 variants compared to the filtering methods used by Byrska-Bishop et al. to produce the GRCh38 1kGP panel. We chose to retain singleton variants, as recent advances in computational phasing now allow for singleton phasing^26^. The inclusion of singleton alleles added 40,226,623 unique variants to our dataset. Overall, variant filtering had removed 46.4% of variants in the Byrska-Bishop et al. 1kGP GRCh38 panel, including 34.1% of all variants that were filtered out solely because they were singletons. In contrast, our filtering strategy removed 22% of 1kGP T2T-CHM13 variants, with 16.3% of variants filtered solely due to the VQSLOD filter (Supplemental Figure 1).

We then used this set of high-confidence variant calls to generate LD-based recombination maps for T2T-CHM13 autosomes and the X chromosome using the software package pyrho^27,28^ (Figure 1). Patterns of LD, and thus recombination maps derived from these patterns, can vary among populations due to differences in demographic history, as well as differences in the locations of crossover recombination events. For this reason, we first inferred recombination maps separately for the 26 subpopulations defined by the 1kGP. A combined global map was then created by averaging all population-specific maps, weighting by respective sample sizes (see Methods). We therefore note that this “global” average is a reflection of the population definitions and sample sizes of the original input data. We also report regions of highest variance in genetic distance across populations (highest variance in centimorgans (cM) per Mb), where the use of population-specific maps may be most advantageous (Supplemental Table S2).

**Figure 1.**
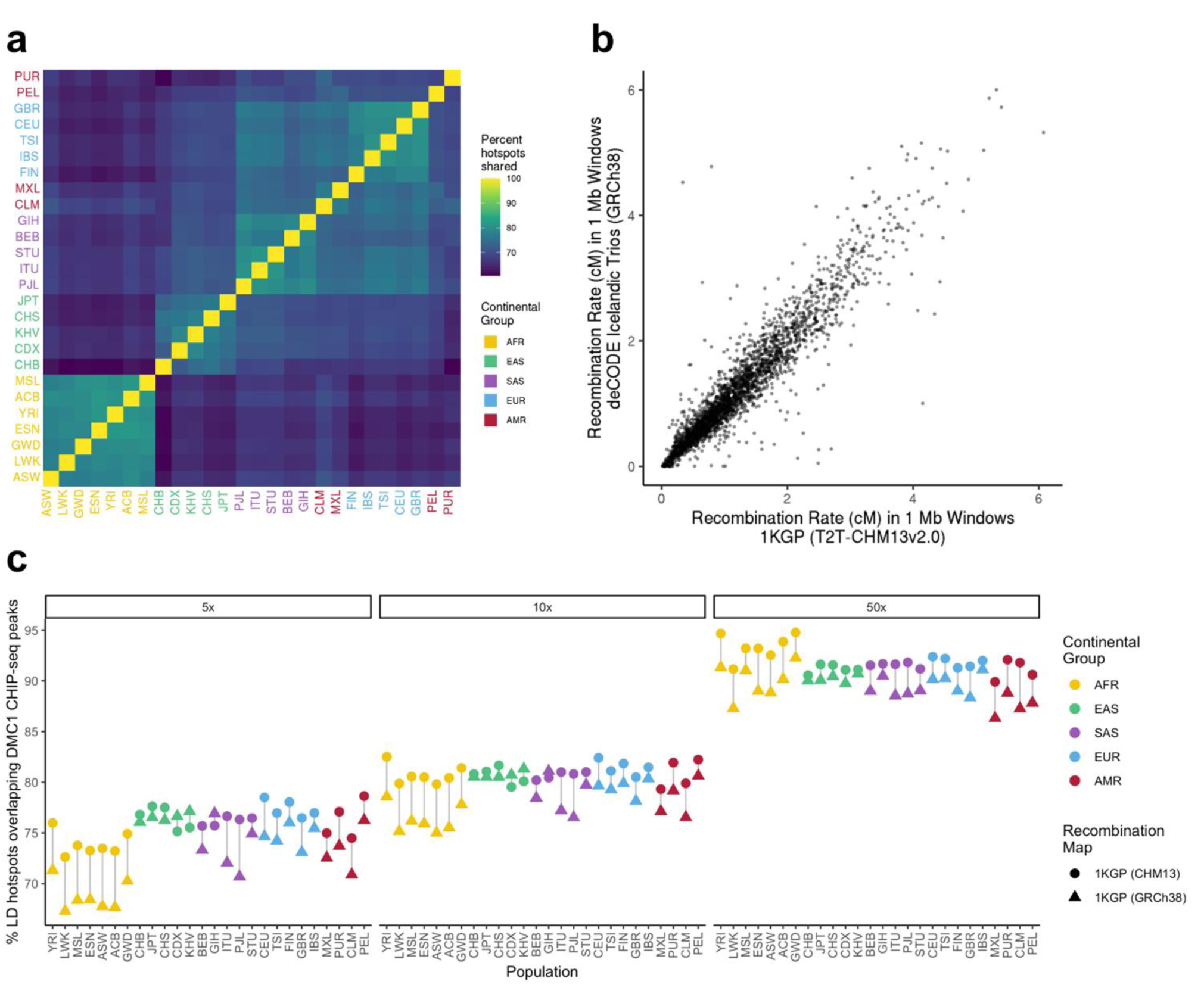
A CHM13v2.0 reference recombination map. a) Percentage of recombination hotspots shared between populations b) Recombination rates in 1MB windows in T2T-CHM13 maps and GRCh38 deCODE maps lifted over to T2T-CHM13. Only shared autosomal regions are represented. Comparison is shown between GBR and deCODE; full set of comparisons are in Supplemental Figure 5. c) Percentage of recombination hotspots overlapping DMC1 ChIP-SSDS peaks from Pratto et al 2014. Data is presented at three separate thresholds for calling recombination hotspots.

For the population-averaged map, we calculated an unadjusted cumulative map length of 2,187 cM, corresponding to a genome-wide sex-averaged recombination rate of 0.72 cM/Mb. This is notably lower than either the pedigree-based 3,615 cM estimate produced by deCODE or the 2,959 cM estimate obtained using cytological observations of meiotic recombination^29,30^. Meanwhile, our estimates are consistent with previous LD-based maps generated from 1kGP data aligned to GRCh38 and analyzed with pyrho^27^ (Supplemental Figure 2). Notably, pyrho applies a smoothing ℓ1 regularization penalty to mitigate variation in recombination rate estimates, damping the strength of some hotspots and resulting in a lower cumulative genetic distance depending on chosen hyperparameters. To address this issue, we scaled our recombination maps to the per-chromosome lengths of the pedigree-based recombination maps from deCODE. Because the default settings of most phasing and imputation tools are tuned using the GRCh38 HapMap2 recombination map^26,30,31^, we also provide our maps scaled to HapMap2 chromosome lengths.

### T2T-CHM13 recombination maps are broadly consistent with corresponding maps generated with GRCh38

To evaluate the validity of the T2T-CHM13-based recombination maps, we performed several additional analyses of their general features, as well as comparison to orthogonal resources. As an internal comparison, we first measured the Spearman correlation between recombination maps for all populations at multiple resolutions (Supplemental Figure 3). Hotspot locations were similar across all populations, but this similarity was highest between populations from the same continental group, replicating past results from corresponding maps generated with GRCh38^27^ (Figure 1a). These trends held across multiple thresholds for defining a recombination hotspot (Supplemental Figure 3). We also compared these recombination maps to previously published LD-based maps generated by applying pyrho to 1kGP variant calls generated via alignment to GRCh38^27^. Spearman correlations between the 1kGP recombination maps generated with the different reference genomes were high across all populations (ρ = 0.914-0.976) (Supplemental Figure 4).

Pedigree-based recombination maps offer an orthogonal source of validation, as they are based on direct detection of recombination events in the sequenced offspring rather than estimation of historical recombination rates from patterns of LD. We thus compared our results to recombination maps generated by deCODE based on detection of crossovers in Icelandic trios (i.e., parents and offspring)^32^ (Figure 1b) and found that Spearman correlations were generally very high (ρ = 0.759-0.979 at 1 Mb resolution). Given the strong correlation in recombination maps between populations^28^, we expect the deCODE pedigree-based and 1kGP LD-based recombination maps to be similar, especially for closely related populations. Consistent with this hypothesis, we observed that for all populations, 74-92% of hotspots detected in our 1kGP LD-based maps were also present in the deCODE map. When we restricted our analysis to hotspots that are shared between all 26 subpopulations, we found that 19,338 of 20,341 (95%) were also identified by deCODE.

### DMC1 ChIP-SSDS peaks validate recombination hotspots in the T2T-CHM13 map

We also compared our recombination maps to the locations of DNA double-strand breaks (DSBs) reported by Pratto et al^33^. While we do not expect recombination hotspots and DSB locations to be identical, we expect a strong correlation, as recombination is initiated by DSBs. Pratto et al. inferred DSB locations using chromatin immunoprecipitation (ChIP) followed by single-stranded DNA sequencing (SSDS) to identify the binding sites of the DSB marker DMC1. To compare the recombination landscapes identified in these two datasets, we realigned ChIP-SSDS data from Pratto et al. to the T2T-CHM13 reference genome and measured the fraction of recombination hotspots (defined according to three different thresholds: 5×, 10×, and 50× the genome-wide average) that overlap DMC1 ChIP-SSDS peaks.

We found that a high proportion of recombination hotspots in both T2T-CHM13 and GRCh38 maps overlap inferred DMC1 binding sites (Figure 1c), increasing as the threshold for calling a hotspot becomes more stringent. Across autosomal regions shared by both assemblies, the proportion of hotspots overlapping DMC1 binding sites is consistently higher in recombination maps aligned to T2T-CHM13 versus GRCh38 (Figure 1c). While only the T2T-CHM13-based recombination maps (and not the GRCh38-based maps) include the X chromosome, we found that a high proportion of recombination hotspots on the X chromosome overlap with DMC1 ChIP-SSDS peaks (Supplemental Figure 5), consistent with genome-wide trends. For both autosomes and sex chromosomes, the proportion of peaks overlapping inferred DMC1 binding sites was significantly higher than expected based on random permutation of recombination hotspot locations (see Methods; P<0.01). Because regions newly resolved by T2T-CHM13 generally remain difficult to access with short-read methods, callable variant density is low in these regions, limiting our ability to assess hotspot–DMC1 overlap.

DSBs are initiated by PRDM9 binding^34,35^, whose variable zinc finger domain determines DNA motif binding specificity. Variation in this zinc-finger domain can cause dramatically different recombination hotspot patterns^36^ and human *PRDM9* genotypes can be broadly clustered into *PRDM9*-A/B and *PRDM9*-C types which recognize distinct DNA motifs^37^. Pratto et al.^33^ measured DMC1 binding in individuals with distinct *PRDM9* genotypes. To investigate this aspect of the data, we divided DMC1 ChIP-SSDS peaks into those occurring in individuals carrying *PRDM9*-A/B versus *PRDM9*-C alleles. We observed that a greater proportion of recombination hotspots from 1kGP samples from the African continental group overlapped the DMC1 ChIP-SSDS peaks from individuals with *PRDM9*-C alleles. Conversely, a greater proportion of recombination hotspots from 1kGP samples from non-African continental groups overlapped the DMC1 ChIP-SSDS peaks from individuals with *PRDM9*-A/B alleles (Supplemental Figures 6-8). These results are consistent with the observations of Pratto et al.^33^ and are expected given the higher frequency of the *PRDM9*-C allele in African populations.

### Pangenomic reference assemblies provide ground truth for haplotype panel evaluation

We then sought to create a haplotype-resolved panel of 1kGP SNP and indel variation generated from T2T-CHM13 alignments that could be used as a suitable reference for statistical imputation and phasing of variation in T2T-CHM13 coordinates. To this end, we used the statistical phasing package SHAPEIT5^26^, which allowed us to pre-phase variants using a pedigree-aware phasing algorithm before applying the classical Li and Stephens HMM^38^. The rare variant phasing algorithm from SHAPEIT5 is also able to phase singleton variants with a switch error rate (SER) of ∼30%^26^. The resulting panel contains 6,404 haplotypes and 101,154,695 unique variants, including 306,634 high-confidence phased variants in previously unresolved regions of the T2T-CHM13 genome.

The standard measure of phasing accuracy is the switch error rate (SER), defined as the percentage of heterozygous variants incorrectly phased with respect to the nearest 5’ heterozygous variant (Supplemental Figure 9)^39–41^. Erroneously phased variants are identified using parental samples as ground truth. However, this approach would require withholding 1,195 duo/trio parents when phasing the dataset (Supplemental Figure 10). Instead, we utilized reference-agnostic, high-quality phased T2T assemblies from the HPRC and HGSVC as ground truth^22,23^. This method had the additional benefit of allowing for the evaluation of phasing accuracy in both trios and statistically phased non-trio samples.

### 1kGP short-read variant calls are more accurate when aligning to the T2T-CHM13 reference genome

Errors in variant calling are known to affect panel phasing accuracy^40^. Therefore, we began by assessing the concordance of our panel’s genotype calls with the HPRC and HGSVC assemblies. An important caveat is that T2T-CHM13 and GRCh38 contain insertions relative to each other, and variants within these insertions cannot be represented in the other reference. T2T-CHM13 contains ∼182 Mb of unique sequence that is not present in GRCh38 (mostly previously unresolved or hard-masked sequence). In contrast, ∼1.2 Mb of GRCh38 represents unique sequence not present in T2T-CHM13^42^. After stratifying variants by shared/unique region status, we compared our T2T-CHM13 reference haplotype panel to the corresponding published GRCh38 panel^43^.

Overall, T2T-CHM13 1kGP variants were 99.08% concordant with assembly-derived variation, while GRCh38 1kGP variants were 98.89% concordant (Table 1). Genotype discordance was higher among indels than SNPs in both panels, particularly for common indels (Table 1, Supplemental Figure 11). Genotyping accuracy did not vary substantially by minor allele frequency (0.02%-50%), consistent with the VQSR method’s design to maintain a uniform false positive rate regardless of allele frequency^20,25,44^. SNPs with only two minor alleles present in the panel (doubletons) were called almost as accurately as SNPs in the overall panel (GRCh38: 0.51% discordant, T2T-CHM13: 0.45% discordant), and singleton SNPs in the T2T-CHM13 panel were only 1.41% discordant with assembly genotypes. When examining variants in shared regions, SNPs from the T2T-CHM13 panel were less discordant with assembled genomes (0.26%) than those from the GRCh38 panel (0.42%). Indels in shared regions followed the same pattern (T2T-CHM13: 4.00%, GRCh38: 4.22%, Table 1). Both panels were more concordant with HPRC assemblies than HGSVC assemblies (Supplemental Figure 12), largely driven by samples from the African continental group, which comprise a greater proportion of the HGSVC dataset and show higher discordance with short-read variant calls. This pattern is consistent with the reduced accuracy of most variant calling tools when analyzing highly diverse genomes^45^.

**Table 1.** Concordance of panel genotypes and phasing with assembly-derived ground truth. Panel variants from 39 HPRC samples were compared to assembly-derived genotypes at overlapping variant sites (sites with ≥1 non-reference allele in both the assembly ground truth and the 1kGP panel). All non-missing genotypes at overlapping sites were evaluated. See Methods for detailed definitions. ^†^ SER, switch error rate: n_switch errors_/n_heterozygous sites_ ^‡^ Discordance rate: When comparing non-missing variant calls to variant representations of sample-reference pangenome assembly differences, the discordance rate is the proportion of variants that do not match. Phasing is not taken into account. Genotypes were evaluated if they were non-missing in both the panel and the vcf representation of the pangenome. (i.e., n_discordant_ / n_genotype calls_, equivalent to 1 − genotype concordance). ^a^Shared genomic regions: sequences with primary alignment between T2T-CHM13 and GRCh38. ^b^ Unshared genomic regions: sequences present in only one reference. For T2T-CHM13, this is ∼182 Mb of previously unresolved sequence with no primary alignment to GRCh38^42^. For GRCh38, this is ∼1.2 Mb of mostly assembly errors that are unable to be aligned to T2T-CHM13.

| Genomic Region | Reference Genome | Variant type | SER<sup>†</sup> (%) | Discordance rate<sup>‡</sup> (%) | # of haplotype switches | # of heterozygous sites | # of discordant variant calls | # of evaluated genotypes |
| --- | --- | --- | --- | --- | --- | --- | --- | --- |
| All regions | CHM13v2.0 | SNPs | 0.149 | 0.257 | 125,429 | 83,970,616 | 1,581,451 | 615,986,668 |
|  |  | Indels | 1.312 | 4.010 | 236,515 | 18,027,890 | 5,285,347 | 131,789,145 |
|  |  | SNPs + Indels | 0.355 | 0.918 | 361,944 | 101,998,506 | 6,866,798 | 747,775,813 |
|  | GRCh38 | SNPs | 0.211 | 0.425 | 203,830 | 96,829,961 | 3,025,689 | 711,355,997 |
|  |  | Indels | 1.385 | 4.440 | 270,802 | 19,555,065 | 6,480,948 | 145,982,877 |
|  |  | SNPs + Indels | 0.408 | 1.109 | 474,632 | 116,385,026 | 9,506,637 | 857,338,874 |
| Unshared genomic regions<sup>a</sup> | CHM13v2.0 | SNPs | 0.497 | 4.597 | 133 | 26,737 | 15,469 | 336,505 |
|  |  | Indels | 2.730 | 9.586 | 410 | 15,019 | 16,586 | 173,023 |
|  |  | SNPs + Indels | 1.300 | 6.291 | 543 | 41,756 | 32,055 | 509,528 |
|  | GRCh38 | SNPs | 0.366 | 2.371 | 450 | 122,930 | 25,015 | 1,054,999 |
|  |  | Indels | 1.123 | 6.925 | 174 | 15,490 | 10,527 | 152,017 |
|  |  | SNPs + Indels | 0.451 | 2.945 | 624 | 138,420 | 35,542 | 1,207,016 |
| Shared genomic regions<sup>a</sup> | CHM13v2.0 | SNPs | 0.149 | 0.254 | 125,296 | 83,943,879 | 1,565,982 | 615,650,163 |
|  |  | Indels | 1.311 | 4.003 | 236,105 | 18,012,871 | 5,268,761 | 131,616,122 |
|  |  | SNPs + Indels | 0.354 | 0.915 | 361,401 | 101,956,750 | 6,834,743 | 747,266,285 |
|  | GRCh38 | SNPs | 0.210 | 0.422 | 203,380 | 96,707,031 | 3,000,674 | 710,300,998 |
|  |  | Indels | 1.385 | 4.437 | 270,628 | 19,539,575 | 6,470,421 | 145,830,860 |
|  |  | SNPs + Indels | 0.408 | 1.106 | 474,008 | 116,246,606 | 9,471,095 | 856,131,858 |

One challenge of using de novo assemblies as ground truth is discrepancy in variant representation; long-read assemblies and short-read variant calls often represent indels differently in repetitive regions^46,47^, making it unclear whether discordant variants in complex, indel-rich regions reflect true genotyping errors or representation differences. Manual inspection of pangenome-discordant indels showed that they were frequently found within multiallelic short tandem repeats (STRs) ‒ a trend that we confirmed with genome-wide analysis (GRCh38: 5.56%, T2T-CHM13: 5.11% indel discordance within STRs; Supplemental Figure 13). In contrast, isolated indels that did not overlap with other variant classes in the panel showed substantially lower discordance rates (GRCh38: 0.53%, T2T-CHM13: 0.49%, Supplemental Figure 14).

Prior analyses of 1kGP variants called using an earlier version of the T2T-CHM13 assembly suggested that ∼25% of non-reference SNV calls in T2T-CHM13-unique regions are false positives^2^. That estimate, however, was based on all GATK-passing variants compared against long-read HiFi sequences as ground truth. Our stricter VQSLOD filter removes 35.3% of GATK-passing variants in T2T-CHM13-unique regions. When all filters are considered, only 5.8% of GATK-passing variants in these regions are retained (Supplemental Figure 1), and those variants have assembly discordance rates 4.60% for SNPs and 9.56% for indels (Table 1).

### The 1kGP T2T-CHM13 haplotype panel is more accurately phased than the 1kGP GRCh38 panel in both previously unresolved and shared regions of the genome

We next sought to quantify the phasing accuracy of each panel. To maximize phasing accuracy, both panels considered Mendelian inheritance rules when phasing trio samples. The GRCh38 1kGP panel used post-phasing error correction on trio proband samples to ensure consistency between parental and child haplotypes. Similarly, our T2T-CHM13 1kGP panel was pre-phased according to Mendelian inheritance logic where possible, with the resulting haplotypes used as a scaffold for statistical phasing^26^. Unlike the GRCh38 panel, the phasing of both probands and parents was performed with trio-based information, and we confirmed that all proband genotypes were phased in a Mendelian-concordant manner. In both panels, non-trio samples were only phased using the Li and Stephens statistical phasing algorithm^38^. As a result, we expect probands (19% of 1kGP samples) and trio parents (37% of 1kGP) to be more accurately phased than samples that are not part of a trio (44%).

Unlike the 1kGP dataset, all HPRC assemblies come from trio probands. Given the fact that the HPRC assemblies were slightly more concordant with our short-read variant calls than HGSVC assemblies (Supplemental Figure 12), we used HPRC assemblies as a ground truth to measure the accuracy of the phasing of the probands in our dataset. Conversely, HGSVC non-trio samples are used as ground truth of statistically phased variant calls throughout this paper.

Phasing accuracy broadly mirrored genotyping accuracy. Across all benchmarked sample groups and variant classes, variants in genomic regions shared between assemblies were phased more accurately in the T2T-CHM13 panel than in the GRCh38 panel (Figures 2 and 3). For both Mendelian pre-phased trio probands (Figure 2a) and statistically phased non-trio samples (Figure 2b), SNP SERs were higher in rare variants. The most common SNPs were phased roughly two orders of magnitude more accurately than the rarest SNPs.

**Figure 2.**
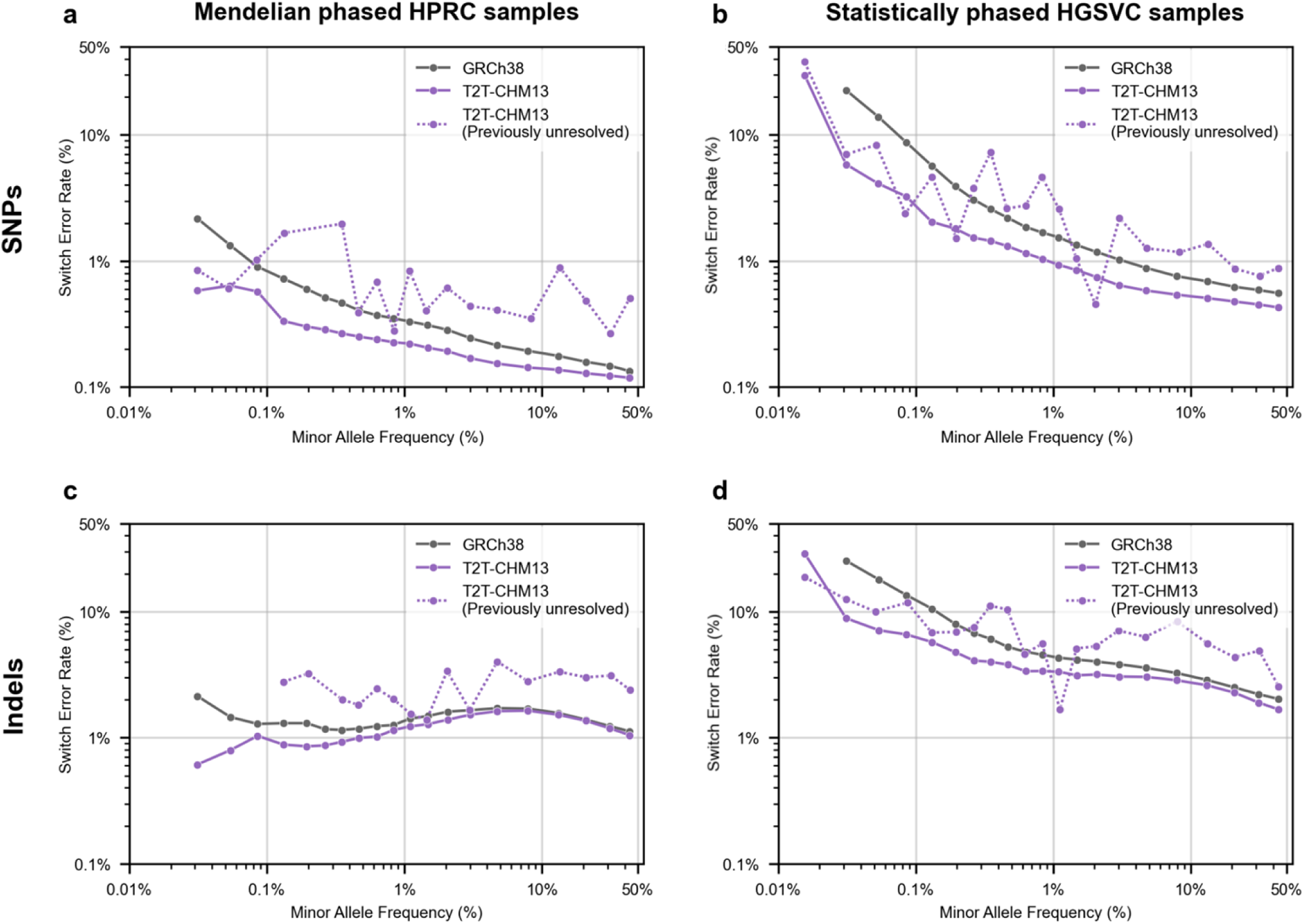
The T2T-CHM13 1kGP haplotype panel is more accurately phased than the GRCh38 1kGP haplotype panel. The T2T-CHM13 1kGP haplotype panel is more accurately phased than the GRCh38 1kGP haplotype panel. Of the 3,202 participants in the 1kGP, 39 had their genomes assembled by the HPRC and 61 by the HGSVC. a) The switch error rate of haplotype panel SNPs from 39 individuals, using their HPRC-assembled genomes as ground truth. Variants located in regions of T2T-CHM13 genome that are not present in GRCh38 are included in overall variant bin error rates but are also plotted separately as T2T-CHM13 (Previously unresolved) variants. b) SER of haplotype panel SNPs from 19 individuals, using their HGSVC-assembled genomes as ground truth. Unlike the 39 HPRC samples, these 19 individuals are not part of a 1kGP trio and therefore were phased without any Mendelian-based error correction. c) Indel SER from the Mendelian pre-phased HPRC samples. d) Indel SER from the 19 individuals phased without Mendelian error correction. Switch error rates and average minor allele frequency per bin are displayed on a log scale.

**Figure 3.**
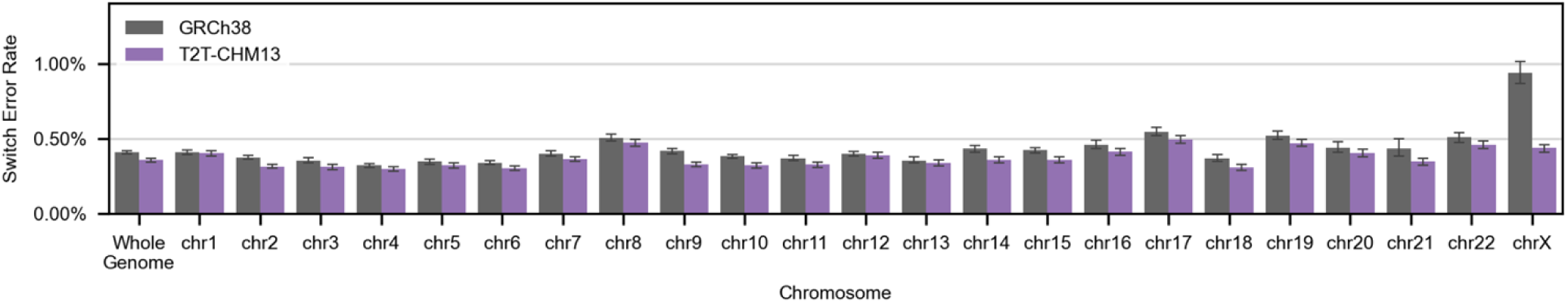
Whole genome and per-chromosome switch error rates from all 100 samples with HPRC or HGSVC assemblies. Ground truth phasing is defined by HPRC or HGSVC assembly. Error bars show +/- 95% confidence interval.

SERs were higher for variants in T2T-CHM13-unique regions than for variants in assembly-shared regions (Figure 2a,b). Indel SERs were higher than SNP SERs in both panels and were relatively stable across the allele-frequency spectrum, particularly among statistically phased samples (Figure 2c, d). Singleton variants were statistically phased with accuracies consistent with previous reports (29.77% SER for SNPs; 28.85% for indels) (Figure 2b,d)^26^.

We noticed that variants from trio probands that underwent Mendelian error correction had notably lower SERs than variants from samples that were not part of a trio. We therefore calculated the estimated SER for each group (using HPRC probands, and HGSVC parents/non-trio participants as a ground truth) to determine an estimated panel-wide SER. As expected, trio probands that underwent Mendelian pre-phasing had lower SERs than statistically phased samples (Figure 2a-d; Supplemental Table S1). Trio parents in the GRCh38 panel were phased only modestly more accurately than statistically phased samples, whereas parents in the T2T-CHM13 panel had SERs closer to their probands (Supplemental Figure 15, Supplemental Table S1). This difference likely reflects the difference in phasing workflows used to create the two panels: in the GRCh38 panel, only trio probands underwent Mendelian error correction after statistical phasing, while conversely for the T2T-CHM13 panel, all trio members underwent Mendelian phasing prior to statistical phasing. Weighting the observed proband/parent/non-trio SER by panel membership (608 probands, 1,195 parents, and 1,399 non-trio individuals) allowed us to estimate a 3,202-member panel-wide SER of 0.90% for the GRCh38 panel and 0.74% for the T2T-CHM13 panel.

When stratified by chromosome, the T2T-CHM13 panel showed lower SERs than the GRCh38 panel across nearly all chromosomes, with the largest relative improvement observed on chromosome X (Figure 3). In statistically phased HGSVC non-trio samples, chromosome X SER decreased from 0.91% in the GRCh38 panel to 0.42% in the T2T-CHM13 panel. Most of this improvement arose in PAR2 and in non-PAR sequence on chromosome X (Supplemental Figure 16). Improved phasing in the pseudoautosomal regions likely reflects the more complete T2T-CHM13 reference sequence, whereas the marked gain in non-PAR chromosome X is consistent with the use of newer SHAPEIT functionality that models the mixed haploid/diploid structure of this chromosome. Together, these changes bring the chromosome X SER in the T2T-CHM13 panel closer to the autosomal range.

### Most switch errors reflect likely genotyping errors instead of phasing errors

Switch errors can be the result of inaccurate phasing or inaccurate genotyping. Inaccurate phasing tends to result in scattered haplotype switches, whereas inaccurate genotyping frequently results in two pseudo-switch errors immediately surrounding the erroneous genotype. This latter form of paired haplotype switches is called a ‘flip’ (Figure 4a, Supplemental Figure 9)^31^. In addition to calculating the SER, we therefore also calculated the flip error rate (FER), defined as the number of flips per heterozygous site, and the “true switch error rate (tSER)”, defined as the SER minus 2 × FER (i.e., the rate of switch errors that are not part of a flip pair).

**Figure 4.**
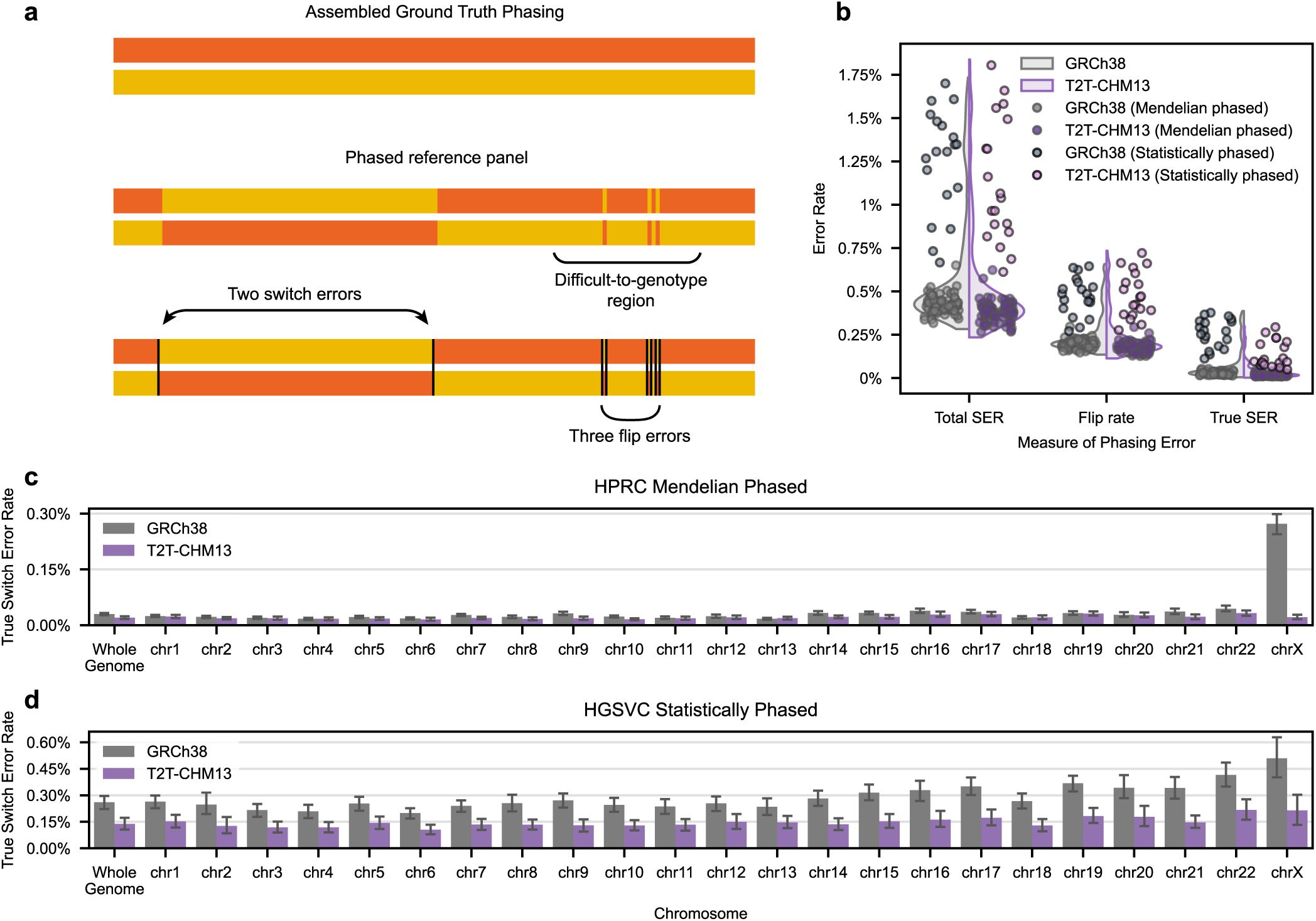
True switch error rates are lower in T2T-CHM13 phased 1kGP haplotypes compared to GRCh38 phased 1kGP haplotypes. a) Throughout this study, we have compared our panel’s phasing with ground truth reference assemblies to identify switch errors, defined as sites at which the maternal and paternal haplotypes switch strands. However, regions which are difficult to sequence can often have genotyping errors that result in flip errors, defined as two consecutive switch errors. Unlike true switch errors, flip errors do not impact the phasing accuracy of nearby variants. Eight switch errors are in the illustrated haplotypes, but only two of these are true switch errors - the other six switch errors are part of three flip errors. b) The SER, flip error rate, and true SER rate of all 100 samples with assembly-based ground truths. Data from the 19 HGSVC individuals that were phased without Mendelian error correction are highlighted. Data points are plotted on top of a violin plot showing the overall distribution of error rates. c) Whole genome and per-chromosome true SERs from 39 HPRC-assembled samples. d) Whole genome and per-chromosome true SERs from 19 HGSVC-assembled samples phased without Mendelian error correction. Error bars show +/- 95% confidence interval.

In our dataset, the majority of switch errors were associated with flips. FERs were correlated with genotyping error rates, while tSERs were uncorrelated with genotyping error rates (SER r^2^ = 0.31, FER r^2^ = 0.35, tSER = 0.006, Pearson correlation, Mendelian pre-phased samples), supporting the interpretation that flips stem from genotyping errors (Supplemental Figure 17). The measured flip error rate for HPRC probands from the 1kGP GRCh38 panel was 0.186%, while the tSER was 0.029%. HPRC probands from the T2T-CHM13 panel had a flip error rate of 0.166% and a tSER of 0.018%. (Figure 4b). As expected, SERs were higher among statistically phased samples compared to Mendelian pre-phased samples.

### True switch error rates in statistically phased samples are 50% lower in the T2T-CHM13 panel compared to the GRCh38 panel

When we excluded flip variants from our analysis, we saw that the T2T-CHM13 tSERs of statistically phased haplotypes often approached the tSER of haplotypes phased via trio concordance (Figure 4b). Mendelian pre-phased samples in the GRCh38 panel had a tSER of 0.029%, while the same samples in the T2T-CHM13 panel had a tSER of 0.018% – a 38% drop in tSER (Figure 4c). For statistically phased samples, tSERs were 50% lower in the T2T-CHM13 panel compared to the GRCh38 panel (tSERs of 0.26% and 0.13%, respectively). (Figure 4d). These differences were especially large on chromosome X, where we observed a 14-fold reduction in the tSERs of Mendelian pre-phased samples (GRCh38: 0.27% tSER; T2T-CHM13: 0.019% tSER). tSERs on acrocentric chromosomes and Chromosome 9 particularly benefited from the phasing workflow used to generate the T2T-CHM13 panel (Figure 4c, d). A large proportion of these chromosomes’ sequences were previously unresolved in the T2T-CHM13 reference genome, which we hypothesized contributed to the phasing improvements. Supporting that hypothesis, we found that the per-chromosome improvement in tSER was associated with the percentage of chromosome length that was previously unresolved (Supplemental Figure 18, p=1.6e-8 by two-sided t-test).

### Impact of use of a T2T-CHM13 1kGP haplotype panel on downstream applications

#### Use of a T2T-CHM13 1kGP haplotype panel improves phasing of out-of-panel samples

To evaluate panel performance for the real-world use case of phasing a new data set, we next assessed the accuracy of variant phasing when phasing out-of-panel samples using our T2T-CHM13 haplotype panel as a reference. For these experiments, we considered the draft human pangenome as ground truth^22^. It is best practice to use reference panels that contain unrelated samples when phasing or imputing genomic variation, so we restricted our analysis to the 2,504 unrelated samples from the 1kGP GRCh38 and 1kGP T2T-CHM13 reference haplotype panels (i.e., unrelated trio parents and non-trio individuals). To avoid leakage between the test and evaluation sets, we additionally removed the parents of pangenomic samples, leaving 2,426 unrelated samples in each reference panel.

We initially evaluated each haplotype panel as a reference for variant phasing. To this end, we used SHAPEIT5 to phase the previously described filtered short-read variant calls from the 39 HPRC samples present in the 1kGP dataset. We applied SHAPEIT5’s common variation algorithm, as rare variant phasing is not available for small datasets or reference panels. When phasing the HPRC short-read variant genotypes, we observed lower SERs when using 1kGP T2T-CHM13 phased haplotypes as a reference haplotype panel (Figure 5). The improvement over use of the 1kGP GRCh38 panel was small and consistent across minor allele frequencies. SNPs and indels in reference-shared genomic regions were phased more accurately than variants in CHM13-unique genomic regions (T2T-CHM13 SNPs: 1.52% CHM13-unique SER vs 1.01% shared SER; T2T-CHM13 Indels: 4.00% CHM13-unique SER vs. 2.10% shared SER).

**Figure 5.**
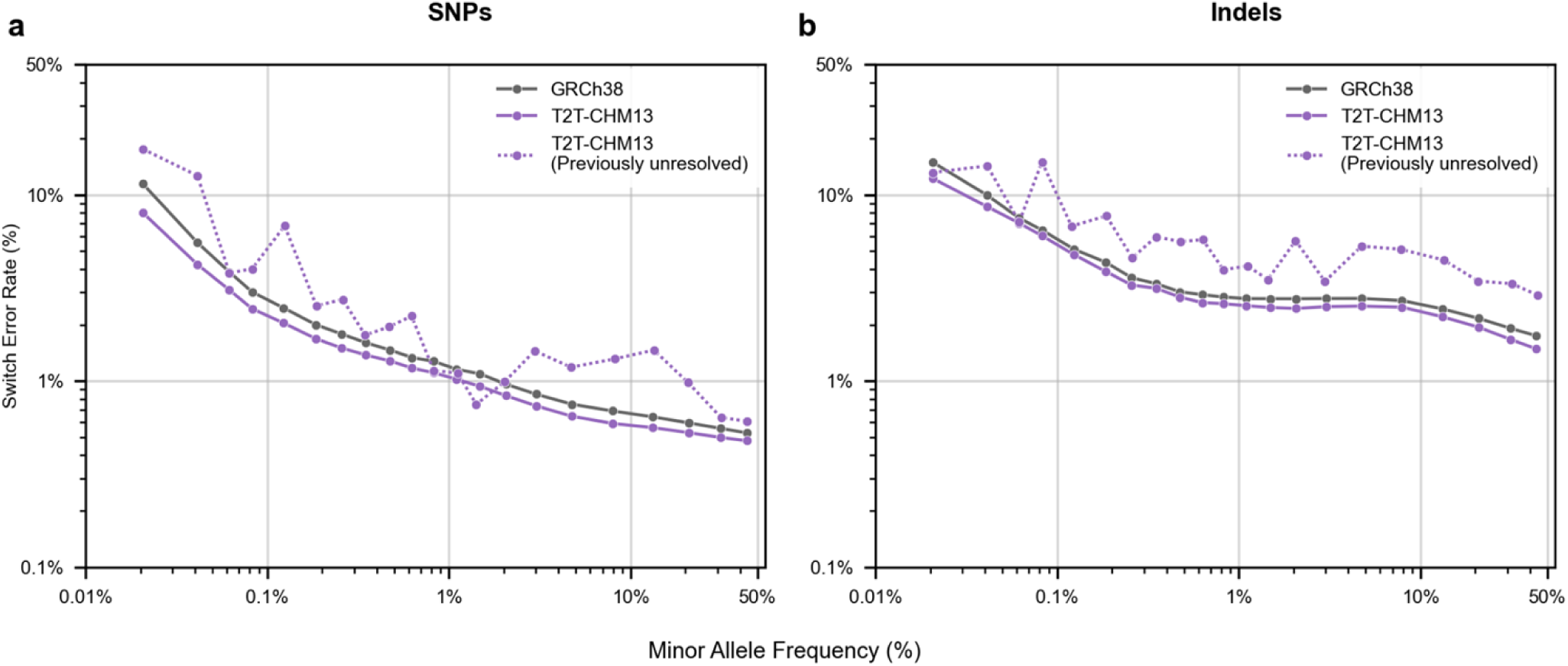
Accuracy of out-of-panel variant phasing when using a GRCh38 or T2T-CHM13 1kGP haplotype panel as reference. All HPRC-assembled samples and their relatives were removed from the GRCh38 1kGP and the T2T-CHM13 1kGP panels of 2,504 unrelated samples, yielding ‘non-HPRC’ panels containing data from 2,426 individuals. Filtered short-read derived SNPs from 39 HPRC-assembled individuals were phased, using either the GRCh38 or T2T-CHM13 2,426 member panel as a source of reference haplotypes for SHAPEIT5. a) SNP and b) Indel SERs were measured by comparison with ground truth pangenomic assemblies, and variants were binned by reference panel minor allele frequency (MAF). SERs and average MAF per bin are displayed on a log scale.

### Genomic regions prone to disease-causing CNVs disproportionately benefit from use of a T2T-CHM13 reference panel

Improvements in out-of-sample phasing performance were not uniform throughout the genome. To gain insight into which genomic regions should be expected to change the most when using the T2T-CHM13 panel to phase, we calculated the SERs of the rephased pangenome samples, stratified by cytogenetic band (Supplemental Table S5). We noticed five of the top 20 regions which benefited the most from the T2T-CHM13 reference panel (Table 2) are co-located with DECIPHER^48^-defined common copy number variant (CNV) disorders, including 22q11.2 (DiGeorge Syndrome) and 15q11.2 (Angelman Syndrome and Prader-Willi Syndrome). Statistical analysis showed that DECIPHER-overlapping cytobands represented 62 of 786 eligible cytobands (7.9%) and 8.9% of cytoband base-pair span, but accounted for 12.6% of positive T2T-CHM13 associated switch error reduction, a 1.64-fold enrichment over random DECIPHER-label permutations (one-sided permutation p = 0.0024). Additionally, DECIPHER-overlapping T2T-CHM13 cytobands had larger reductions in SER CHM13-associated SER than would be expected by chance (0.147% vs. 0.109%; p = 0.0036, one-sided weighted permutation test).

**Table 2.** Top 15 chromosomal regions with an improved switch error rate when using a T2T-CHM13 panel to phase short-read variant calls. 1kGP short-read variant calls from 39 samples with HPRC assemblies were phased using either a panel of GRCh38 reference haplotypes from unrelated individuals, or a T2T-CHM13 reference haplotypes from unrelated individuals. The parents of the 39 HPRC samples were removed from both panels prior to analysis, resulting in reference panels with haplotypes from 2,426 individuals. Switch errors were calculated with phased pangenomic variation as a ground truth. Regions were sorted by the relative difference in SER when using the GRCh38 panel or the T2T-CHM13 panel. Regions associated with recurring copy number variant syndromes listed in the DECIPHER^48^ database are highlighted in grey.

| Genetic Region | GRCh38 panel |  |  |  | T2T-CHM13 panel |  |  |  | Relative reduction in SER | Relevant Medical Conditions |
| --- | --- | --- | --- | --- | --- | --- | --- | --- | --- | --- |
|  | Num. of alt genotypes | Num. of het sites | GT error rate (%) | Switch error rate (%) | Num. of alt genotypes | Num. of het sites | GT error rate (%) | Switch error rate (%) |  |  |
| 10q11.22 | 580390 | 70591 | 5.63% | 2.23% | 322394 | 44215 | 4.09% | 0.89% | 59.93% | Developmental delay (case reports)<sup>88</sup> |
| 6p21.32 | 914834 | 203990 | 1.94% | 0.40% | 767662 | 171939 | 0.74% | 0.20% | 50.70% | HLA Locus. Many autoimmune disorders<sup>89,90</sup>, SYNGAP1 related disorders<sup>91</sup> |
| 9p13.1 | 312420 | 43454 | 2.38% | 1.71% | 241082 | 35374 | 0.93% | 0.88% | 48.25% | Case reports of developmental delay<sup>92</sup> |
| **16p12.2** | 535588 | 71002 | 1.91% | 1.74% | 440490 | 63059 | 1.21% | 0.92% | 47.25% | 16p12.2 deletion syndrome, variable penetrance<sup>93</sup> |
| **15q11.2** | 1003012 | 151040 | 3.56% | 1.07% | 721733 | 111488 | 0.80% | 0.57% | 46.85% | Start of Prader-Willi / Angelman imprinting region<sup>94</sup> |
| **8p23.1** | 2058897 | 315945 | 1.45% | 1.12% | 1447837 | 226297 | 0.64% | 0.61% | 45.46% | 8p23.1 deletion<sup>95</sup>/duplication<sup>96</sup> syndromes |
| 2q12.2 | 410177 | 65312 | 0.83% | 0.82% | 356355 | 58469 | 0.66% | 0.45% | 44.25% | Ectodermal dysplasias<sup>97</sup> |
| 14q11.2 | 1500779 | 235393 | 4.55% | 1.78% | 1183890 | 195995 | 1.09% | 1.03% | 41.93% | Developmental delay, CHD8-associated ASD<sup>98</sup> |
| **15q13.1** | 584572 | 87031 | 1.39% | 1.28% | 462218 | 72298 | 0.82% | 0.75% | 41.16% | End of Prader-Willi / Angelman imprinting region<sup>49,94</sup> |
| Xq13.2 | 245349 | 18645 | 2.19% | 1.42% | 212160 | 16538 | 2.07% | 0.86% | 38.93% | Deletions/duplications associated with ID<sup>99</sup> |
| Xq22.3 | 703365 | 63798 | 1.11% | 1.06% | 624078 | 57598 | 0.84% | 0.65% | 38.87% | Alport syndrome<sup>100</sup> |
| 11q22.2 | 197399 | 30579 | 0.69% | 0.59% | 166861 | 25921 | 0.61% | 0.37% | 38.08% | Neuroblastoma<sup>101</sup> |
| 14q31.2 | 338419 | 48894 | 0.74% | 0.61% | 317172 | 45660 | 0.51% | 0.38% | 37.54% | Case reports of microcephaly/ID<sup>102</sup> |
| 2q21.1 | 705785 | 101019 | 3.49% | 1.08% | 495197 | 73730 | 2.59% | 0.68% | 36.84% | ADHD w/ ID<sup>103</sup> |
| **22q11.21** | 995554 | 145817 | 1.67% | 1.20% | 1150482 | 172333 | 1.44% | 0.76% | 36.79% | DiGeorge Syndrome<sup>104</sup> |

To investigate the nature of this improvement, we calculated the panel genotype error rate in non-overlapping 10 kb intervals across the genome. We also determined the SER of short-read variation from pangenomic samples phased using either panel as a reference. As a general trend, we observed that both panel genotype error rates and out-of-panel phased SERs tended to peak on either side of regions of the genome associated with CNV-related disorders (Figure 6). Improvements in panel genotyping accuracy and out-of-panel phasing accuracy were observed within CNV disorder regions, but were much more apparent in their flanking regions. This pattern was true for both panels, but was especially pronounced when genetic variation was phased with the GRCh38 panel. Often, these spikes were associated with a drop in variant density, suggesting that these are regions which are difficult to genotype. In some instances, such as the loci flanking the 22q11 region, the T2T-CHM13 panel genotypes were not more accurate than the GRCh38 panel, but phasing error was still substantially improved. This suggests that improvements in phasing with the use of a T2T-CHM13 panel are not solely due to improved panel genotyping.

**Figure 6.**
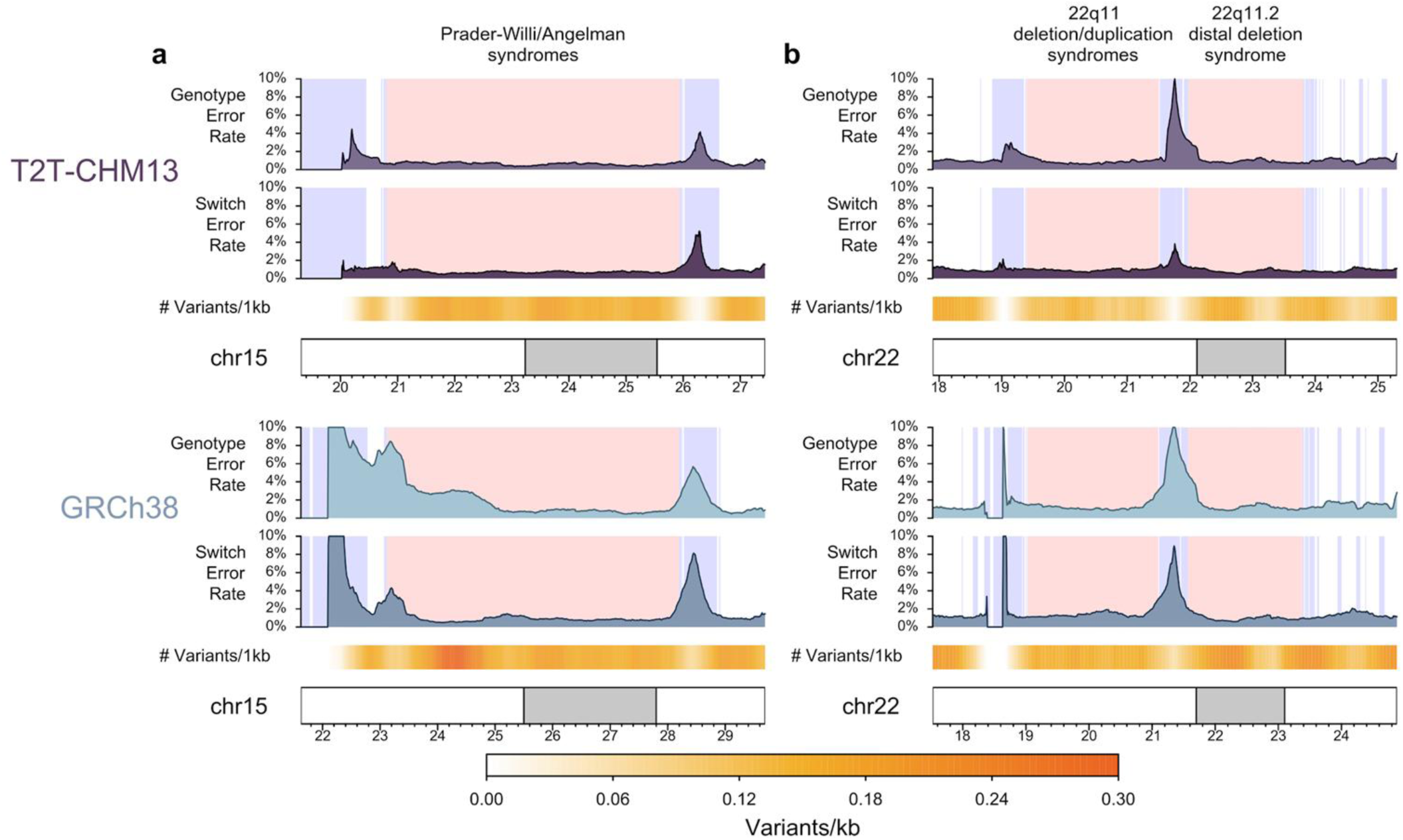
Genotyping and switch error rates spike in areas flanking CNV disorder associated regions when phasing out-of-panel HPRC samples. Unphased GRCh38 or T2T-CHM13 1kGP short read variant calls from 39 HPRC members were phased using a reference-matched panel of 2,426 unrelated 1kGP haplotypes. Genotyping error rates, switch error rates, and variant density were calculated in a 500kb rolling window of 10kb increments across each chromosome. These statistics were then plotted in regions (+/-1.5mb) associated with the copy number variant disorders a) Prader-Willi/Angelman syndromes and b) 22q11 disorders. DECIPHER-defined CNV disorder regions are highlighted in red (T2T-CHM13 coordinates are lifted from GRCh38 coordinates), and segmental duplication regions are highlighted in blue.

The Prader-Willi/Angelman region on chromosome 15 is a notable example of the effects of using a complete reference on variant calling and phasing. This region harbors parent-of-origin-imprinted genes whose disruption underlies Prader-Willi and Angelman syndromes^49^. Accurate genotyping and phasing is therefore essential to understand disease mechanisms and potential modifying variants within this region^50^. Disease-causing CNV disorders in this region, like most CNVs, commonly arise from non-allelic homologous recombination between flanking segmental duplications (SDs)^51^. Similarities between SDs lead to mismapping reads and false heterozygous genotype calls. We observe this pattern in our dataset (Supplemental Figure 19). The 15q11-q13 locus is rich in SDs (Figure 6a), including the Prader-Willi breakpoint 2 region (chr15:23,097,000–23,447,333). In the GRCh38 panel, we see a peak in genotype discordance and switch error rates at this site. This peak is entirely absent in the T2T-CHM13 panel (Figure 6a, Supplemental Figure 20). The lower T2T-CHM13 panel error rates are unlikely to be due to reference sequence differences - the two references are 99.9% identical at this locus. However, a complete representation of segmental duplications is likely to lead to fewer mismapping reads, and indeed the T2T-CHM13 reference panel is more accurate in these regions (Supplemental Figure 21). This example demonstrates that the T2T-CHM13 reference can improve variant calling even in regions that are well resolved in GRCh38 by reducing read mismapping.

### Use of a T2T-CHM13 panel improves imputation of rare SNPs

We next sought to assess the effect of using the T2T-CHM13 1kGP reference haplotype panel when imputing missing genetic variation. Variant imputation is most commonly used to increase the resolution of microarray or low-coverage sequencing-based genomic studies, as whole-genome sequencing (WGS) can be cost prohibitive. To mimic this use case, we downsampled GRCh38 and T2T-CHM13 WGS data from 256 individuals from the Simons Genome Diversity Project (SGDP)^52^, isolating the set of variants that are assayed by Illumina Omni 2.5 genotyping array. We chose this dataset because it is highly diverse (containing 60 populations compared to 26 populations in 1kGP) and it is one of the few well-studied, publicly available genotype datasets that has been mapped to both reference genomes using the same pipeline^24^.

To ensure that imputation accuracy was based on the same set of variants in both builds, we compared variants imputed using the GRCh38 panel with variants imputed using the T2T-CHM13 panel lifted to GRCh38 coordinates (Figure 7a) and vice versa (Figure 7b). In both cases, we only examined variants present in both the lifted over and originally produced panel. We initially tested Picard LiftoverVCF^53^ and Bcftools +liftover^19^ to compare panels, but encountered significant limitations in their handling of indels in repetitive regions. We therefore developed LiftoverIndel, which accounts for differences in indel representation between reference assemblies (Supplemental Note 1).

**Figure 7.**
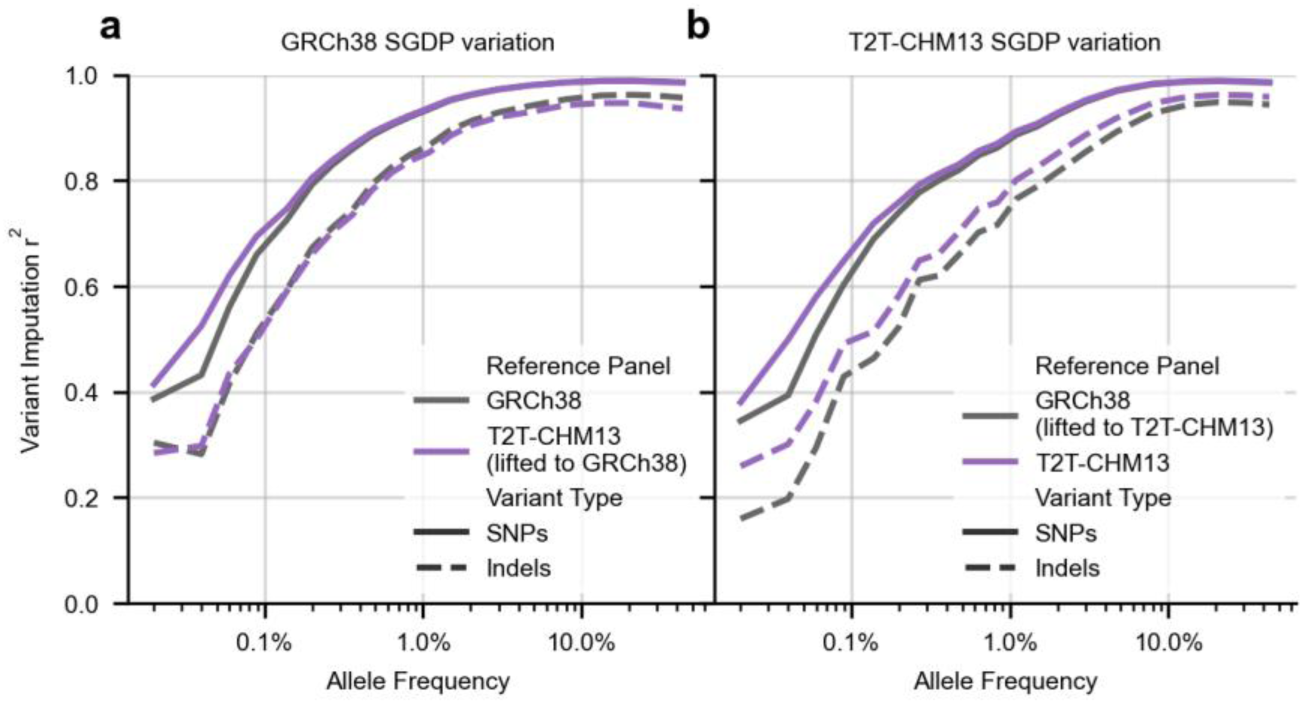
Imputation of genomic variation in 256 non-1kGP Human Genome Diversity Project (HGDP) samples, using 1kGP haplotype panels as references. a) Genotyping array data was simulated by downsampling variation derived from short reads aligned to GRCh38 to those variants present in the Infinium Omni2.5 genotyping array. Variation was imputed with the 1kGP GRCh38 reference panel or the 1kGP T2T-CHM13 reference panel lifted to GRCh38. b) Downsampled CHM13v2.0 HGDP variation was phased and imputed with the 1kGP GRCh38 reference haplotype panel or the 1kGP T2T-CHM13 reference panel lifted to GRCh38 coordinates. Variants that were not present in both panels were removed from analysis. All imputed variants were binned by the minor allele frequency of the variant in the reference panel used during imputation. r^2^ values were calculated per bin. r^2^ statistics are stratified by SNPs and Indels. Average minor allele frequency per bin is displayed on a log scale.

When imputing SGDP variation from pseudo-array variant calls, use of the T2T-CHM13 1kGP reference panel resulted in comparable or slightly improved imputation accuracy compared to the use of the GRCh38 reference panel. For SNPs, imputation was consistently enhanced by the use of the T2T-CHM13 panel, even when working with a lifted over panel in GRCh38 coordinates (Figure 7a). For indels, imputation was less accurate than for SNPs, regardless of the reference panel, as previously observed^26^. Indel imputation was best when working with a panel that had not been lifted between builds. As previously reported, the 1kGP GRCh38 panel produced lower imputation accuracy for individuals from African and Oceanic populations versus individuals from American, European, and Asian populations, likely due to the absence or limited representation of these populations in 1kGP^43^. We also observed population-specific differences in imputation accuracy using both panels, although no single population was disproportionately affected by the use of either panel (Supplemental Figure 20, 21).

The T2T-CHM13 1kGP haplotype panel we have generated includes singleton variants, which are not typically included in haplotype panels used for imputation. Therefore, to assess the impact of including singleton variants on imputation accuracy, we compared SGDP samples imputed with reference panels that included or excluded 1kGP singletons (Supplemental Figures 22, 23). We observed a consistent but miniscule reduction in per-sample imputation accuracy when singletons were included (median Δr² on the order of 10⁻⁴–10⁻³).

However, we also saw an increase in rare variant imputation accuracy, especially among rare indels (Δr² ∼=+0.02). Accordingly, we include singletons in the released panel for completeness; however, users who wish to maximize overall imputation accuracy may elect to remove singletons before imputation.

## Discussion

In this study, we used previously published T2T-CHM13-aligned 1kGP short-read DNA variant calls to produce a recombination map and phased imputation panel for the T2T-CHM13 reference genome. Previous adoption of updated reference genomes by the genetics community has been slow due to concerns about inadequate secondary reference resources and uncertain benefits^54^. Our goal was to contribute to the ecosystem of resources that enable the adoption of the T2T-CHM13 reference and to demonstrate the benefits of using this resource in common genomics workflows.

Thorough benchmarking of the panel showed that, compared to the commonly used GRCh38 1kGP panel^43^, the T2T-CHM13 panel is more accurately phased. In addition, outside samples are phased more accurately and single nucleotide variants in SGDP samples are more accurately imputed from the subset of sites on a standard genotyping array (Illumina Omni) when using the T2T-CHM13 panel.

In addition to the higher accuracy and completeness of the T2T-CHM13 reference^1,2^, there are several other differences that could explain the improvements in phasing and imputation accuracy observed when using these reference panels. While the variant calling pipelines used to create each panel were largely identical, the phasing pipelines were not. The GRCh38 1kGP panel was phased using an older software package (SHAPEIT-duohmm v2), which only performs Mendelian phasing of trios as a post-hoc correction. Due to technical limitations, this correction step was not performed for chromosome X. In contrast, the updated SHAPEIT5 package performs trio-based phasing first for both autosomes and chromosome X, using the trio-phased samples as a source of information when phasing non-trio samples.

Additionally, different variant filters were used in the production of each panel. The T2T-CHM13 panel includes singleton variants, while the GRCh38 panel does not, enhancing rare variant imputation. The T2T-CHM13 panel was also generated using a stricter VQSR threshold than the GRCh38 panel to address deficiencies in variant calibration within novel genomic regions. The use of this stricter threshold removed an additional 17% of genetic variants from the T2T-CHM13 panel.

While devising a benchmarking strategy, we discovered that the draft human pangenome^22^ could be a useful source of ground truth for genotyping and phasing. While the assemblies in the pangenome are not reference quality, they are close enough to provide a meaningful measure of genotyping and phasing accuracy. As an example, the HPRCv1.1 HG002 assembly differs from the reference-quality HG002 Q100 assembly at only 0.00016% of sites and exhibits one switch error every 45 Mb^46^. All pangenome assemblies were generated in the same manner as the HG002 assembly, supporting the utility of the draft human pangenome as a ground truth data set.

Prior efforts to measure the effect of reference bias on genotyping accuracy have been limited to a collection of well-characterized samples^2,55,56^. The pangenome assemblies allowed us, for the first time, to directly benchmark the genotyping accuracy of short-read variant calls from an ancestrally diverse collection of 100 individuals generated from either GRCh38 or T2T-CHM13 aligned reads. We observed that, in comparison to the GRCh38 reference panel, the SNP phasing and genotyping in the T2T-CHM13 reference panel was substantially more consistent with the HPRC and HGSVC assemblies.

Throughout this study, we observed much higher discordance rates among indels versus SNPs. Most indels occurred in repetitive or complex genomic regions (Supplemental Figure 13). Excluding variants that overlap other variants from our datasets revealed that for both SNPs and indels, variants outside of these complex regions are better phased and genotyped in the T2T-CHM13 panel. This is consistent with previous reports of differences in variant decomposition leading to isolated indels being represented as multiple variants^46^.

Computationally intensive haplotype-based benchmarking methods perform local realignment to solve this problem^57,58^, and recent advances in haplotype alignment algorithms^46,47^ have raised the possibility of benchmarking the accuracy of entire assemblies against a ground truth reference. However, whole genome multiple sequence alignment still poses a computational challenge, making consistent variant representation difficult in cohort studies.

As the error rates of genotyping and phasing algorithms continue to decline, variant normalization challenges will represent a relatively larger source of genotype discrepancies. While graphical representations of genomes present a promising solution to this problem, they raise the question of how to define a genetic variant. The concept of a variant requires a reference sequence from which to define both the position and the nature of the variation; even in graphical genome representations, a variant as currently defined requires a reference node from which the alternate path emerges. In the future, we hope consensus can be reached on how to conceptualize genetic differences in the absence of a shared reference. Until then, linear reference genomes are a necessary tool to understand genetic differences.

Importantly, this panel was produced using short-read-based variant calls. Short-read genotyping often fails to resolve large-scale variation, and as a result can produce hotspots of genotype errors in regions prone to structural variants^59^. As an example, one potential cause of genotyping and phasing errors around common disease-causing regions is a prevalence of inversions, which would lead to phasing and genotyping errors around the breakpoints. This is a common limitation in the field and applies to most available reference haplotype panels. However, this limitation is particularly challenging in the case of T2T-CHM13, as most of the previously unresolved sequence in this reference is inaccessible to short-read variant calling^2^. Large structural variation, centromeric satellites, telomeres, and surrounding regions experience unique dynamics of mutation, recombination, and selection^60–62^. Future development of a panel that accurately represents these regions will be crucial for clarifying these dynamics.

Using T2T-CHM13 as a reference genome poses further challenges when using the functionally equivalent GATK pipeline. This pipeline requires reference datasets to calibrate variant quality thresholds, and those datasets have not been generated for T2T-CHM13. (The Rhie et al. T2T-CHM13 1kGP dataset was calibrated using GRCh38 variation lifted to T2T-CHM13 coordinates.) The absence of training data in T2T-CHM13 unique regions means that VQSLOD calibration is likely to be poor in these regions, and the elevated discordance rates we observe in these regions may partly reflect this miscalibration. Strict quality filtering can yield credible variant calls, but further improvements in variant calling and quality calibration will be possible once new T2T-CHM13 resources are generated.

In our hands, variant calling and phasing accuracy within these newly accessible regions remains poor; however, including these regions in the haplotype panel has broader impacts across the genome, improving variant calling and phasing in short-read-accessible regions of the genome. We anticipate additional population-scale long-read assemblies from the HPRC and other consortia, which should enable accurate variant calling, phasing, and imputation within newly accessible regions, unlocking these complex loci for association studies, evolutionary analyses, and other applications^22,63–65^. Until those datasets are available, our short-read-based resources can be paired with previously released genome accessibility masks that define regions of T2T-CHM13 where variant calls are most reliable^24^.

We hope that our study will improve the practical value of the complete human reference genome for common genetic applications. Our work demonstrates that the benefits of using this reference are especially pronounced around disease-associated CNV loci, with potential implications for both fundamental biological research and clinical genetics. To facilitate these goals, the recombination maps and reference panel are freely available at https://github.com/marbl/CHM13.

## Resource Availability

### Lead Contact

Further information and requests for resources and reagents should be directed to and will be fulfilled by the lead contact, Donna Werling.

### Materials availability

This study did not generate new unique reagents.

### Data and code availability

- T2T-CHM13 population-specific and global averaged recombination maps are available at https://zenodo.org/records/19601957 and https://github.com/JosephLalli/phasing_T2T/tree/main/resources/recombination_maps/t2t_native_scaled_maps
- SHAPEIT5v1.1 can be found at https://github.com/odelaneau/shapeit5 [commit ID 3ba8b5c]
- SHAPEIT5v1.1 with ability to measure flip events can be found at https://github.com/JosephLalli/shapeit5/tree/main [commit ID e6dd007]
- 1kGP T2T-CHM13 phased variant calls are available at https://s3-us-west-2.amazonaws.com/human-pangenomics/index.html?prefix=T2T/CHM13/assemblies/variants/1000_Genomes_Project/chm13v2.0/Phased_SHAPEIT5_v1.1
- 1kGP T2T-CHM13 unphased variant calls are available at https://s3-us-west-2.amazonaws.com/human-pangenomics/index.html?prefix=T2T/CHM13/assemblies/variants/1000_Genomes_Project/chm13v2.0/all_samples_3202/
- 1kGP GRCh38 phased variant calls are available at https://ftp.1000genomes.ebi.ac.uk/vol1/ftp/data_collections/1000G_2504_high_coverage/working/20220422_3202_phased_SNV_INDEL_SV/
- 1kGP unphased variant calls are available at https://ftp.1000genomes.ebi.ac.uk/vol1/ftp/data_collections/1000G_2504_high_coverage/working/20201028_3202_raw_GT_with_annot/
- Combined HGSVC3 and HPRC pangenome available in T2T-CHM13 coordinates at https://ftp.1000genomes.ebi.ac.uk/vol1/ftp/data_collections/HGSVC3/release/Graph_Genomes/1.0/2024_02_23_minigraph_cactus_hgsvc3_hprc/hgsvc3-hprc-2024-02-23-mc-chm13-vcfbub.a100k.wave.norm.vcf.gz
- Combined HGSVC3 and HPRC pangenome available in GRCh38 coordinates at https://ftp.1000genomes.ebi.ac.uk/vol1/ftp/data_collections/HGSVC3/release/Graph_Genomes/1.0/2024_02_23_minigraph_cactus_hgsvc3_hprc/hgsvc3-hprc-2024-02-23-mc-chm13.GRCh38-vcfbub.a100k.wave.norm.vcf.gz
- GATK Resource bundle in T2T-CHM13 coordinates can be found at https://s3-us-west-2.amazonaws.com/human-pangenomics/index.html?prefix=T2T/CHM13/assemblies/variants/GATK_CHM13v2.0_Resource_Bundle/
- T2T-CHM13 short-read accessibility mask is at https://s3-us-west-2.amazonaws.com/human-pangenomics/T2T/CHM13/assemblies/annotation/accessibility/combined_mask.bed.gz
- Simons Genome Diversity Project variant calls can be found in T2T-CHM13 coordinates at https://s3-us-west-2.amazonaws.com/human-pangenomics/index.html?prefix=T2T/CHM13/assemblies/variants/SGDP/chm13v2.0/ and GRCh38 coordinates in the T2T_ChrY Anvil workspace at https://anvil.terra.bio/#workspaces/anvil-datastorage/AnVIL_T2T_CHRY under SGDP_GRCh38_chromosome
- The pedigree of 1kGP samples can be found at http://ftp.1000genomes.ebi.ac.uk/vol1/ftp/data_collections/1000G_2504_high_coverage/working/1kGP.3202_samples.pedigree_info.txt
- Bed file of T2T-CHM13 unique regions is at https://s3-us-west-2.amazonaws.com/human-pangenomics/T2T/CHM13/assemblies/chain/v1_nflo/chm13v2-unique_to_hg38.bed
- CHM13v2.0 cytoband coordinates can be found at https://s3-us-west-2.amazonaws.com/human-pangenomics/T2T/CHM13/assemblies/annotation/chm13v2.0_cytobands_allchrs.bed
- Chain files to lift from T2T-CHM13 to GRCh38 and vice-versa are at https://s3-us-west-2.amazonaws.com/human-pangenomics/T2T/CHM13/assemblies/chain/v1_nflo/grch38-chm13v2.chain
- Code used to lift variation between T2T-CHM13 and GRCh38 is at https://github.com/JosephLalli/LiftoverIndel [Commit ID 370e322]
- Code used to generate recombination maps is at https://github.com/andrew-bortvin/1kgp_chm13_maps
- Panel phasing scripts are available at https://github.com/JosephLalli/phasing_T2T [commit ID 020230a]
- Code to create figures is at https://github.com/mccoy-lab/1kgp_chm13_maps [commit ID a4247a7] and https://github.com/JosephLalli/phasing_T2T. [commit ID 020230a]
- Original code used to generate the figures in this publication has been deposited at Zenodo at https://zenodo.org/record/19905132 and is publicly available as of the date of publication.
- Any additional information required to reanalyze the data reported in this paper is available from the lead contact upon request.

## Acknowledgements

The authors would like to thank Dr. Oliver Delaneau and the T2T Consortium for their generosity and advice during the development of our benchmarking methods and Dr. Samantha Shapiro for her constructive feedback during manuscript drafting. This paper could not have been attempted, let alone completed, without Nils Irland and the generous computing support of the University of Wisconsin-Madison Laboratory of Genetics and the UW-Madison Center for High Throughput Computing. Thank you to the Johns Hopkins Department of Biology and Center for Computational Biology, as well as members of the McCoy laboratory for constructive feedback. We also thank the staff of Advanced Research Computing at Hopkins (ARCH) for computing support. Research reported in this publication was supported by the National Institute of General Medical Sciences of the National Institutes of Health under Award R35GM133747 to RCM, the Brain and Behavior Research Foundation 29815 to DMW, US Department of Agriculture National Institute of Food and Agriculture Hatch Act Formula Fund WIS04078 to DMW, Simons Foundation Autism Research Initiative 606289 to DMW, and the University of Wisconsin-Madison Medical Scientist Training Program T32GM140935 to JLL. We would like to acknowledge the Human Pangenome Reference Consortium (BioProject ID: PRJNA730823) and its funder, the National Human Genome Research Institute (NHGRI). The content is solely the responsibility of the authors and does not necessarily represent the official views of the National Institutes of Health.

## Author Contributions

Conceptualization: J.L.L., A.N.B., R.C.M., D.M.W.; Methodology, investigation: J.L.L. and A.N.B. Writing—original draft, J.L.L. and A.N.B.; writing—review & editing, J.L.L., A.N.B., R.C.M., D.M.W.; funding acquisition, R.C.M., D.M.W.; resources, R.C.M., D.M.W.; supervision, R.C.M., D.M.W.

## Declaration of Interests

The authors declare no competing interests.

## Supplemental Information

**Document S1.** Supplementary Note 1, Supplementary Methods, Figures S1–S26

**Supplementary Data S1.** Table of variance in recombination rates in 1MB windows.

**Supplementary Data S2.** Per-sample variant counts, error counts, switch error rates, and genotyping error rates stratified by panel, method of phasing, and ground truth data source.

**Supplementary Data S3.** Per-MAF bin variant counts, error counts, switch error rates, and genotyping error rates stratified by panel, method of phasing, unique/nonunique status, and ground truth data source.

**Supplementary Data S4.** Imputation summary statistics by subject, panel, variant category, and unique/nonunique status.

**Supplementary Data S5.** Cytoband and disease-CNV region summary statistics.

## Methods

### Dataset generation

#### Input callsets and pedigree information

Prior work from our group used the functionally equivalent GATK pipeline to align and call Illumina paired-end 30× WGS datasets from the 3,202 1000 Genomes Project samples to the T2T-CHM13v2.0 reference genome, producing a joint-called, VQSR-calibrated unphased variant callset. Pedigree relationships were obtained from the 1kGP pedigree file and were used to identify parent–offspring trios and duos for Mendelian pre-phasing and downstream validation (data sources and accessions are listed in Data availability and Supplementary Methods).

#### Variant normalization and filtering

Prior to phasing, multiallelic sites were decomposed into biallelic records and variants were left-shifted using Bcftools (v1.21). Chromosome X variants were forced to diploid representation using the bcftools fixploidy plugin. Variants in assembly-shared regions were annotated, the mendelian2 bcftools plugin was used to count Mendelian violations, and per-superpopulation HWE p-values were calculated using bcftools fill-tags plugin.

After exporting all annotations to a tsv file, we excluded all variants that failed quality filters. Quality filters consisted of removing variants with a VQSR < 0, and did not include filtering singleton variants. Otherwise, variant filtering was identical to those used to produce the GRCh38 1kGP haplotype panel (see Supplemental Methods). Other than removing structural variants >50bp in size, star alleles, and variants with an MAC of 0, no additional filtration was performed on the GRCh38 panel.

### Generating CHM13 full-length recombination maps

#### Map Generation

We generated population-specific recombination maps from the unphased, filtered T2T-CHM13 genotypes using pyrho^28^ utilizing previously published historical population size estimates^66^. For each of the 26 populations, we selected optimal hyperparameters to generate the maps. To create a single, unified map, we scaled individual population maps to deCODE^67^ cumulative map lengths and then computed a sample-size-weighted average recombination rate across all populations. The X chromosome was handled separately, dividing it into PAR and non-PAR regions to account for differences in ploidy and demographic history (see Supplemental Methods).

#### Comparisons of Recombination Maps and Hotspots

We compared our T2T-CHM13 recombination map to existing maps by binning them into non-overlapping 1 Mb, 100 kb, and 10 kb intervals and measuring the Spearman correlation of genetic distance within each bin. We defined recombination hotspots as regions with a rate at least 10-fold greater than the genome-wide average (with alternative 5× and 50× thresholds) and used bedtools^68^ intersect to quantify the overlap between hotspot landscapes of different maps.

To compare recombination hotspots with sites of meiosis-initiating DNA double-stranded breaks (DSBs), we reanalyzed DMC1 ChIP-SSDS data, aligning it to T2T-CHM13v2.0. We identified DMC1 ChIP-SSDS peaks and assessed their intersection with our LD-based hotspots, separating peaks by PRDM9 subtype. A null distribution for the expected overlap was generated by randomly shuffling hotspot locations.

### Phasing T2T-CHM13 1000 Genomes Project variant calls

#### Phasing strategy

Filtered T2T-CHM13 variant calls from the 3,202 member 1kGP variant panel were statistically phased using SHAPEIT5v1.1.1’s two step protocol. First, common variants (MAF ≥ 0.1%) were phased using SHAPEIT5_phase_common with non-default parameters optimized for accuracy, including an effective population size (Ne) of 135,000, a PBWT depth of 8, and an HMM window of 5 cM. Chunking was not employed at this step. Rare variants (MAF < 0.1%) were subsequently phased onto this scaffold using SHAPEIT5_phase_rare in 40 Mb chunks with 1 Mb overlap. Outputs were joined with bcftools concat.

Pedigree information was provided to enable Mendelian pre-phasing. Chromosome X was split into PAR and non-PAR regions and separately phased, with non-PAR regions phased in a haploid-aware manner using an Ne of 101,250 (75% of autosomal Ne). Complete parameter specifications are provided in Supplementary Methods.

### Statistical Analysis of Phased Variant Panels

#### Genomic region stratification

##### Assembly shared/assembly-unique regions

”Assembly-shared” regions are defined as T2T-CHM13 sequences with primary alignment to GRCh38. “CHM13-unique” regions (∼182 Mb) are sequences unique to T2T-CHM13 with no primary alignment to GRCh38, encompassing centromeric satellites, segmental duplications, acrocentric short arms, and other complex sequences that were unresolved or hard-masked in GRCh38. “GRCh38-unique” regions (∼1.2 Mb) are GRCh38 sequences lacking alignment to T2T-CHM13, typically representing assembly errors or false duplications that were corrected in T2T-CHM13. Annotations defining these regions were obtained from Vollger et al 2022^42^.

##### Short tandem repeat (STR) regions

Variants were annotated using two complementary annotation sets. “GIAB STR regions” were defined using the GIAB v3.6 AllTandemRepeats stratification bed files for each reference assembly (Dwarshuis et al. 2024^69^). “Platinum STR regions” were defined using tandem repeat annotations from the Platinum Genomes Project^70^, which cover a larger proportion of the genome.

##### Segmental duplication regions

Variants were stratified by overlap with segmental duplication (SD) annotations from Vollger et al. 2022.

##### Overlapping and isolated variants

We classified variants based on whether their allelic representations overlapped another variant in the panel. A variant was classified as “overlapping” if multiple variants shared the same genomic position, or if the variant’s reference allele span overlapped a position containing multiple variants. Exact file sources and annotation methods are available in the Supplementary Methods.

#### Phasing accuracy

To evaluate phasing accuracy, we calculated error rates against ground truth haplotypes obtained from VCF representations of the joint HPRC v1.1/HGSVC3 pangenome assemblies, comprising 39 samples with HPRC assemblies (all trio probands) and 61 samples with HGSVC assemblies (including 29 trio probands, 13 trio parents, and 19 non-trio samples). This stratification allowed us to separately assess error rates for Mendelian pre-phased samples versus statistically phased samples. To ensure consistent variant representation between the HaplotypeCaller-produced variation and the pangenome vcf, we used bcftools norm to atomize variants. Variants were then left-shifted and split into biallelic variation.

Switch errors were defined as heterozygous variants who change from haplotype 1 (h_1_) to haplotype 2 (h_2_) relative to the previous heterozygous variant, with ground truth haplotypes defined as those represented in the HPRCv1.1-HGSVC pangenome. Flip errors were defined as pairs of consecutive switch errors, i.e., when a heterozygous site is a switch error and the immediately preceding heterozygous site was also a switch error. True Switch Errors were defined as switch errors that are flanked by heterozygous sites that are not switch errors. See Supplemental Figure 9 for an illustration of these concepts. Switch error rate (SER), flip error rate (FER), and true switch error rate (tSER) are all defined as the percentage of heterozygous sites that are erroneous. Specifically, SER = (number of switch errors) / (number of adjacent heterozygous pairs evaluated), reported as a percentage. FER = (number of flip errors) / (number of heterozygous sites evaluated) and tSER = (number of true switch events) / (number of heterozygous sites evaluated), each reported as a percentage. SHAPEIT5_switch was modified to count flips and true switch errors (https://github.com/JosephLalli/shapeit5). Metrics were computed genome-wide, and stratified by allele frequency and/or genomic region annotation.

#### Genotype concordance with pangenome assemblies

For the 100 1kGP samples with haplotype-resolved assemblies, we assessed genotype concordance between the 1kGP short-read callsets and normalized and filtered HPRCv1.1-HGSVC pangenome callsets. Briefly, GRCh38 and T2T-CHM13 phased haplotype panels were subset to the same list of 100 samples, and monomorphic variant calls were removed. Genotype concordance was evaluated using SHAPEIT5_switch, which compares genotypes only at sites present in both datasets (intersection-based evaluation). Concordance rate is defined as the fraction of compared genotypes that are identical (not considering phase) between query and truth datasets. Sites containing missing genotypes in either dataset were excluded from analysis.

Genotype discordance is equal to 1 - genotype concordance. Concordance summaries were stratified by allele frequency and genomic context, and computed separately for SNPs and indels.

#### 3,202-member panel-wide error rate estimation

To estimate panel-wide error rates, we stratified the 100 samples with assembly-based ground truth by their trio status: trio probands (n=68), trio parents (n=13), and non-trio samples (n=19). Trio probands and parents underwent Mendelian pre-phasing, while non-trio samples were statistically phased. Because these categories have different expected error rates and different prevalence in the 1kGP panel (608 probands, 1195 parents, and 1399 unrelated non-trio samples), we calculated weighted average error rates by weighting each group’s error rate by its prevalence in the panel.

### Measuring performance in out-of-panel samples

To determine the switch error rate of out-of-panel variants, we removed the 78 parents of HPRCv1.1 samples from the 1kGP 2,504-member unrelated panel to produce a 2,426-member panel. Additionally, we created unphased variant files that only contained the 39 1kGP participants present in the HPRCv1.1 panel. We then applied the same method outlined above to phase these 39 HPRC members’ variation. We were unable to use the SHAPEIT5_rare program on the resulting phased vcf file, as the lowest possible MAF was 1/78 (1.3%), and SHAPEIT5’s rare variant algorithm is not designed to operate on common variants.

### Regional error rate analyses

To identify genomic regions with elevated error rates, we calculated error statistics at multiple resolutions. Cytoband-level switch error and genotype discordance rates were computed by aggregating per-variant counts within each cytoband. CNV disorder region coordinates were obtained from the DECIPHER database and used to subset per-variant error statistics for regions associated with recurrent copy number variation syndromes.

For visualization of error rate variation across chromosomes, we computed rolling average error rates using a 500 kb window. Each reference genome was divided into non-overlapping 10 kb bins, and for each bin we calculated the number of switch errors and heterozygous sites evaluated. A rolling window spanning ±24 bins (490 kb total, plus the center bin) was then applied, with window-level error rates calculated as the sum of errors divided by the sum of heterozygous sites across all bins in the window. Results were plotted using karyoploteR^71^.

### SGDP Imputation and evaluation

To quantify the relative accuracy of variants imputed with either the GRCh38 or T2T-CHM13 panels, we performed genotype imputation into 256 unrelated Simons Genome Diversity Project (SGDP)^24,52^ participants using IMPUTE5 v1.2.0^72^. Target genotypes were first downsampled to Illumina Omni 2.5 array sites, then pre-phased against the 2,504-unrelated-sample 1kGP reference panel using SHAPEIT5_phase_common with default settings before imputation (full parameters in Supplementary Methods). Imputation accuracy was measured using r², defined as the squared Pearson correlation between imputed dosages and true genotypes within allele-frequency bins, calculated using GLIMPSE2_concordance^7^ with the --gt-val flag. Imputation accuracy was additionally stratified by the ancestry group assignments provided by the SGDP to assess panel performance across globally diverse populations.

### Coordinate harmonization and liftover

For analyses requiring coordinate conversion between GRCh38 and CHM13v2.0 (e.g., comparing imputation accuracy across reference builds), we used LiftoverIndel (https://github.com/JosephLalli/LiftoverIndel), which extends standard chain file-based liftover to handle indels in regions where reference sequences differ. The algorithm adjusts variant representations when lifted coordinates overlap sequence differences between assemblies, with genotype flipping when the alternate allele becomes the reference in the target assembly (detailed procedure in Supplementary Methods). Sites that could not be unambiguously lifted were excluded from cross-assembly comparisons. Chain files were obtained from the T2T Consortium (see Resources).

### Statistical analysis and reproducibility

Per-variant and per-sample counts (switch errors, flip errors, heterozygous sites checked, genotype calls) were obtained using a modified version of SHAPEIT5_switch that distinguishes flip errors from true switch errors. These counts were aggregated across chromosomes, MAF bins, and stratification categories into analysis dataframes using polars^73^ (v1.30.0). Summary statistics (switch error rate, flip error rate, true switch error rate, genotype discordance) were then calculated from these aggregated counts as described in Supplementary Methods. Complete workflows, analysis scripts, and a Docker container (jlalli/phasing_T2T) with all dependencies are available in the GitHub repository (https://github.com/JosephLalli/phasing_T2T).

## Notes

https://zenodo.org/records/17178669

https://github.com/JosephLalli/phasing_T2T

https://s3-us-west-2.amazonaws.com/human-pangenomics/index.html?prefix=T2T/CHM13/assemblies/variants/1000_Genomes_Project/chm13v2.0/Phased_SHAPEIT5_v1.1/

https://zenodo.org/records/19601957

https://github.com/mccoy-lab/1kgp_chm13_maps

### Competing Interest Statement

The authors have declared no competing interest.

### Summary of Updates

In addition to numerous small changes to the text to improve clarity, key revisions include: Added orthogonal validation of our recombination maps using DMC1 ChIP single-stranded DNA sequencing (SSDS) data (Pratto et al. 2014), demonstrating strong concordance between LD-based hotspots and double-strand DNA break locations across autosomes, the X chromosome, and newly resolved regions of T2T-CHM13 (Fig. 1c). Stratified genotyping and phasing error rates by genomic context (STR regions, segmental duplications, isolated variants), clarifying that elevated indel error rates are driven by complex repetitive regions rather than systematic panel quality issues (Supplemental Figures 13, 19). Evaluated the impact of including singleton variants in the reference panel on downstream imputation accuracy, demonstrating minimal effect on overall r2 while providing modest improvements for rare variant imputation (Supplemental Figures 23, 24). Revised manuscript language to report singleton phasing accuracy as an explicit SER of ∼30%. Substantially refactored all analysis code to improve reproducibility, portability, and readability. Repositories now include explicit version/commit IDs, MIT licenses, and test datasets enabling users to replicate key analyses. Added analysis comparing short-read variant call concordance with HPRC versus HGSVC assemblies, identifying sample-specific factors that contribute to differences in concordance rates between assembly sources (Supplemental Figure 12).

