## Supplemental Text for "A T2T-CHM13 recombination map and globally diverse haplotype reference panel improves phasing and imputation"

### Supplemental Note 1

Comparing variant calls across genome assemblies requires translating genomic coordinates while preserving allele representations that remain valid in the target reference. Standard liftover tools such as Picard LiftoverVcf[53] use chain files[74] to map coordinates between assemblies, but they do not explicitly model sequence differences between the source and target references. When a lifted variant falls in a region where the two assemblies differ, naive coordinate translation can produce a VCF record whose REF allele no longer matches the target reference. This problem is particularly acute for indels, because indel representation depends on the surrounding reference context. We developed LiftoverIndel to address these limitations.

##### Inputs

LiftoverIndel takes as input a VCF to be lifted, a chain file, the target reference FASTA, and a VCF cataloguing sequence differences between the two assemblies. While the other inputs are standard in most liftover tools, the VCF catalogue is not. It can be generated from both references and a chain file using the ‘to_vcf.py’ file in chaintools[75]. Before lifting over variation, LiftoverIndel catalogues and indexes the VCF into a dictionary of intervaltrees[76] (one tree per chromosome), allowing for rapid identification of between-reference variants that overlap a given genomic range.

##### Coordinate conversion and strand handling

After indexing, variants to be lifted are reviewed iteratively. Variants that share the same position are grouped and lifted together. For each variant at a position, variant start and end coordinates are converted to the target assembly using pyliftover[77]. If a variant maps to the reverse strand, its REF and ALT alleles are reverse-complemented, and the new position of the anchor base is recalculated. Next, lifted coordinates are queried against the between-reference difference interval trees to identify variants landing in regions where the two assemblies differ. Currently, variants overlapping multiple reference differences are flagged as unliftable due to ambiguous interpretation. We hope to add this feature in the future.

##### Direct-overlap adjustment for between-reference differences

When a lifted variant directly overlaps a single between-reference difference, LiftoverIndel rewrites the REF and ALT alleles to account for the known sequence change between assemblies. For SNPs overlapping a reference SNP, the REF base is replaced with the target-assembly base. For indels overlapping a reference indel, both REF and ALT are updated to reflect the combined effect of the sample variant and the assembly difference in target coordinates, after which common suffixes are trimmed to restore a minimal VCF representation. In some cases, incorporating the reference difference makes REF and ALT identical, indicating that the source variant corresponds to an allele already present in the target reference. When this occurs, the variant is flipped so that the target-reference allele is restored as REF. Because records at the same source position must remain internally consistent, all co-located variants are flipped together.

overlapping a reference SNP, the REF base is replaced with the target-assembly base. For indels overlapping a reference indel, both REF and ALT are updated to reflect the combined effect of the sample variant and the assembly difference in target coordinates, after which common suffixes are trimmed to restore a minimal VCF representation. In some cases, incorporating the reference difference makes REF and ALT identical, indicating that the source variant corresponds to an allele already present in the target reference. When this occurs, the variant is flipped so that the target-reference allele is restored as REF. Because records at the same source position must remain internally consistent, all co-located variants are flipped together.

##### Haplotype realignment

The direct-overlap adjustment described above is necessarily local: it only modifies alleles when a reference difference falls within the lifted variant interval itself. However, a nearby between-reference indel can change the local sequence context enough that a valid source-assembly indel no longer has the same minimal representation in the target assembly, even if the two do not directly overlap. Coordinate translation alone may therefore yield a formally valid VCF record that is not the representation a variant caller would produce if the same haplotype were called directly against the target reference. This is distinct from a left-alignment problem: the allele itself can change class, for example from an insertion in the source assembly to a SNP in the target assembly.

To resolve such cases, LiftoverIndel searches for nearby between-reference indels within a configurable distance of each lifted indel (50 bp by default). The algorithm selects the nearest such indel and attempts to define a local window that contains both the lifted variant and the nearby between-reference indel, but no additional between-reference differences. Within that window, realignment proceeds in four steps:

1. **Reconstruct the local source-reference haplotype.** Starting from the target reference sequence, LiftoverIndel applies the selected between-reference indel to recover the homologous local source-reference sequence.

2. **Apply the sample variant.** The source ALT allele is substituted into this reconstructed source-reference haplotype, producing the full alternate haplotype carried in the source assembly.

3. **Align to the target reference.** This alternate haplotype is globally aligned back to the local target-reference sequence using Needleman-Wunsch alignment. Both left-favoring and right-favoring tie-breaking are evaluated, and the most parsimonious valid target-reference representation is retained.

4. **Extract the minimal variant.** The variant implied by the alignment is extracted, normalized by left-alignment and common-affix trimming, and used as the lifted representation.

This procedure compares complete local haplotypes rather than isolated variant records. Direct-overlap adjustment asks how a lifted REF/ALT pair should be edited when a reference difference falls inside the variant interval. Haplotype realignment asks a more fundamental question: what is the minimal target-reference representation of the same alternate haplotype? In repetitive or indel-rich sequences, these are not equivalent. By reconstructing the source haplotype and re-expressing it against the target reference, LiftoverIndel produces variant representations that more closely match those generated by local-reassembly callers such as GATK HaplotypeCaller.

As an example, consider the CHM13 source variant chr22:48543900:A:AG, an insertion of G into a poly-A tract. After coordinate conversion, this variant maps to GRCh38 position chr22:48051306. The between-reference difference VCF records a nearby one-base difference at chr22:48051295 (CA→C), indicating that GRCh38 contains one additional A in this region relative to CHM13. This difference is visible in the local reference sequences, where the GRCh38 window is one base longer than the homologous CHM13 window:

```
GRCh38  AAATACAAAAAAAAAAAAAAAAT
CHM13   AAATAC-AAAAAAAAAAAAAAAT
```

Critically, this assembly difference at position 48051295 lies 11 bases upstream of the lifted variant at position 48051306: close enough to alter the local sequence context, but outside the lifted variant interval itself. Because the direct-overlap adjustment only rewrites alleles when a reference difference falls within the variant interval, it does not apply in this case, and the source insertion representation is carried through unchanged as chr22:48051306:A:AG.

Haplotype realignment reaches a different conclusion. Applying the source ALT allele to the CHM13 reference produces the alternate haplotype shown below, alongside the two reference sequences:

```
GRCh38 ref         AAATACAAAAAAAAAAAAAAAAT
CHM13 normed ref   AAATAC-AAAAAAAAAAAAAAAT
CHM13 ref          AAATACAAAAAAAAAA-AAAAAT
CHM13 alt          AAATACAAAAAAAAAAGAAAAAT
```

After accounting for the upstream assembly indel, the CHM13 alternate haplotype and the GRCh38 reference are the same length. Aligning the alternate haplotype against the GRCh38 reference reveals a single-base mismatch rather than an insertion:

```
GRCh38 ref          AAATACAAAAAAAAAAAAAAAAT
CHM13 alt           AAATACAAAAAAAAAAGAAAAAT
                                    *
```

The correct target-assembly representation is therefore chr22:48051306:A:G, a SNP rather than an insertion. In other words, once the nearby assembly difference is taken into account, the source alternate haplotype differs from the GRCh38 reference at only a single base. The upstream assembly indel absorbs the apparent length change, converting what was an insertion in CHM13 coordinates into a substitution in GRCh38 coordinates. Imputation performance confirms this distinction: the realigned SNP representation achieves lifted dosage $r^2 = 0.9997$, whereas the un-realigned insertion representation is essentially uninformative ($r^2 < 0.002$).

##### Output and post-processing

LiftoverIndel outputs successfully lifted variants together with separate files for variants that could not be lifted because of missing chain-file coordinates, multiple overlapping reference differences, or allele mismatches. Source coordinates and alleles are preserved as INFO tags (SRC_CHROM, SRC_POS, SRC_REF_ALT) to enable provenance tracking. Lifted variants are subsequently normalized with bcftools norm, sorted, and re-annotated with updated allele frequencies using bcftools +fill-tags.

##### Comparison with bcftools +liftover

During manuscript revision, we benchmarked LiftoverIndel against bcftools +liftover[19], which was released during the preparation of this manuscript and includes an indel-aware local-alignment rescue step. The two tools differ in their realignment strategy: bcftools +liftover maps the 5' and 3' anchors of a maximally extended variant representation and uses local alignment when an anchor falls at the edge of a chain gap and cannot be mapped directly, whereas LiftoverIndel reconstructs the full source-assembly alternate haplotype and re-derives the variant from a global alignment against the target reference. In our evaluation, indel imputation $r^2$ from panels lifted with LiftoverIndel was consistently equal to or higher than from panels lifted with bcftools +liftover (Supplementary Figures 23, 24). This pattern is consistent with LiftoverIndel's haplotype-level realignment producing variant representations closer to those emitted by GATK HaplotypeCaller[44], the caller used to generate the original 1kGP call set.

##### Implementation

LiftoverIndel is implemented in Python and requires cyvcf2[78], pyliftover[77], intervaltree[76], biopython[79], numpy[80], and tqdm[81]. The tool is available at https://github.com/JosephLalli/LiftoverIndel.

### Supplementary Methods

#### S1. Software, versions, and reference files

All analyses were performed using T2T-CHM13v2.0 as the primary reference assembly (FASTA: https://s3-us-west-2.amazonaws.com/human-pangenomics/T2T/CHM13/assemblies/analysis_set/chm13v2.0.fa.gz), with GRCh38 resources used only for cross-assembly comparisons and stratifications. VCF/BCF manipulation, normalization, filtering, and annotation were performed using bcftools (v1.21) and htslib (v1.21). Genomic interval operations used bedtools (v2.31.0). Statistical phasing was performed using SHAPEIT5 (v1.1.1, static binaries obtained from github repository, commit 990ed0d). Phasing accuracy was computed using a modified build of SHAPEIT5_switch (https://github.com/JosephLalli/shapeit5) that distinguishes flip errors from true switch errors, as described in Section S7. Imputation was performed using IMPUTE5 (v1.2.0, static binary), and imputation accuracy summaries were computed using GLIMPSE2_concordance. Recombination maps were generated using pyrho with demographic models from smc++. DMC1 ChIP-SSDS ChIP-seq analysis used bwa-mem2[82] for alignment and MACS2 (v2.2.9.1) for peak calling via the nf-core/ssds pipeline. Data analysis and visualization were performed using Python (polars, seaborn, matplotlib) and R (karyoploteR). All software dependencies for phasing and imputation analysis are available in a Docker container (jlalli/phasing_T2T) on Docker Hub.

Reference interval tracks used for stratification include: assembly correspondence annotations from Vollger et al. 2022 (chm13v2-unique_compared_to_hg38.bed from https://github.com/marbl/CHM13), GRCh38-unique regions (hg38.GCA_009914755.4.synNet.summary.bed.gz from https://hgdownload.soe.ucsc.edu/goldenPath/hs1/vsHg38/), segmental duplication annotations from Vollger et al. 2022, GIAB v3.6 AllTandemRepeats stratification bed files, and Platinum Genomes tandem repeat annotations. Cytoband coordinates were obtained from UCSC[74], and CNV disorder intervals were obtained from the DECIPHER database. Chain files for coordinate conversion (chm13v2-hg38.over.chain and hg38-chm13v2.over.chain) were obtained from the T2T Consortium (https://s3-us-west-2.amazonaws.com/human-pangenomics/T2T/CHM13/assemblies/chain/).

#### S2. Input datasets and sample sets

##### S2.1 1kGP T2T-CHM13 cohort callset

We analyzed the 1kGP high-coverage cohort callset aligned to T2T-CHM13v2.0, comprising 3,202 individuals with autosomes and chromosome X (available at https://s3-us-west-2.amazonaws.com/human-pangenomics/index.html?prefix=T2T/CHM13/assemblies/variants/1000_Genomes_Project/chm13v2.0/all_samples_3202/). This callset was generated using the functionally equivalent GATK HaplotypeCaller pipeline applied to Illumina paired-end 30X whole-genome sequencing data aligned with bwa-mem. Sample identifiers, population labels (26 populations across 5 superpopulations), and sex annotations were obtained from 1kGP release metadata. Pedigree relationships used for Mendelian QC and pedigree-aware phasing were obtained from the 1kGP pedigree file (http://ftp.1000genomes.ebi.ac.uk/vol1/ftp/data_collections/1000G_2504_high_coverage/1000G_698_related_high_coverage.ped).

For comparison, we used the consortium-released 1kGP GRCh38 phased panel (https://ftp.1000genomes.ebi.ac.uk/vol1/ftp/data_collections/1000G_2504_high_coverage/working/20220422_3202_phased_SNV_INDEL_SV/). Unphased, unfiltered 1kGP GRCh38 variant calls were obtained from the consortium ftp server (https://ftp.1000genomes.ebi.ac.uk/vol1/ftp/data_collections/1000G_2504_high_coverage/working/20190425_NYGC_GATK/).

##### S2.2 Sample subsets used in analyses

Several sample subsets were used throughout the analyses. The full 3,202-sample panel includes 608 trio probands, 1,195 trio parents (who also underwent Mendelian pre-phasing), and 1,399 unrelated non-trio samples (statistically phased only). The 2,504-sample unrelated panel excludes all trio parents and was used for imputation reference panels. For out-of-panel phasing evaluation, we created a 2,426-sample panel by additionally removing the 78 parents of HPRC-assembled samples from the unrelated panel. Sample lists defining these subsets are available in the GitHub repository under resources/sample_subsets/.

#### S2.3 Ground truth resources for phasing benchmarking

Phasing evaluation used independent truth resources derived from haplotype-resolved assemblies. The primary truth resource was the combined HGSVC3 and HPRC pangenome VCF, released in February 2024 (available in CHM13v2.0 coordinates at https://ftp.1000genomes.ebi.ac.uk/vol1/ftp/data_collections/HGSVC3/release/Graph_Genomes/1.0/2024_02_23_minigraph_cactus_hgsvc3_hprc/hgsvc3-hprc-2024-02-23-mc-chm13-vcfbub.a100k.wave.norm.vcf.gz and in GRCh38 coordinates at the corresponding GRCh38 URL). This unified callset contains 100 samples overlapping with 1kGP: 39 HPRC-assembled samples (all trio probands) and 61 HGSVC-assembled samples (including 29 trio probands, 13 trio parents, and 19 non-trio samples).

#### S2.4 Out-of-panel cohorts for imputation benchmarking

Imputation benchmarking used 256 unrelated individuals from the Simons Genome Diversity Project (SGDP)[52,2]. SGDP variant calls were obtained in T2T-CHM13 coordinates (https://s3-us-west-2.amazonaws.com/human-pangenomics/index.html?prefix=T2T/CHM13/assemblies/variants/SGDP/chm13v2.0/). GRCh38 calls generated by the same group were obtained from the T2T_ChrY Anvil workspace.

### S3. Variant preprocessing, tag derivation, and QC filtering

#### S3.1 Normalization and representation standardization

1kGP VCF files were left-aligned and normalized relative to the reference FASTA. Multiallelic records were split into biallelic records using bcftools norm -m -any.

A representative normalization command sequence:

```
bcftools norm -Oz -f reference.fa -m -any -W -o $normalized_calls $input_vcf
```

#### S3.2 Cohort-level annotation to create INFO tags used for filtering

Several INFO tags used for filtering and stratification were computed from the normalized cohort callset. Site-level missingness (F_MISSING) was computed as the fraction of samples with missing genotypes at each site using bcftools +fill-tags -- -t F_MISSING. Cohort minor allele frequency (MAF) and minor allele count (MAC) were computed using bcftools +fill-tags -- -t AN,AC,MAF,MAC:1=MAC.

Hardy-Weinberg equilibrium metrics were computed within each of the five superpopulations (EUR, AFR, EAS, AMR, SAS) to avoid confounding due to ancestry mixture. Per-superpopulation HWE p-values were computed using bcftools +fill-tags -Ou -S population_ids.txt - -- -t HWE, producing tags HWE_EUR, HWE_AFR, HWE_EAS, HWE_AMR, and HWE_SAS. bcftools +fill-tags -Ou - -- -t HWE was additionally applied to calculate overall HWE scores. Sites were flagged if either the overall HWE score or the maximum superpopulation score violated the HWE threshold ($p < 1\times10^{-10}$).

Mendelian inconsistency counts were computed for trio samples using bcftools +mendelian2 --ped pedigree.ped -m a -m d, which counts Mendelian errors under both autosomal and diploid X chromosome inheritance models. The MERR tag records the number of Mendelian errors at each site. Variants with Mendelian errors occurring at more than 5% of called genotypes were excluded from analysis.

An example annotation command is below:

```
    bcftools view -Ou -r $region $normalized_calls \
    | bcftools norm -Ou -f $ref_fasta -m -any - \
    | bcftools annotate -Ou -a $syntenic_bedfile --mark-sites +SYNTENIC \
                    -c "$chrom_specific_syntenic_annotation_line_part1" \
                -H "$chrom_specific_syntenic_annotation_line_part2" \
                    -x INFO/MAC,INFO/AN,INFO/AC,INFO/MAF \
                --set-id '%CHROM\_%POS\_%REF\_%FIRST_ALT' - \
    | bcftools +mendelian2 -Ou - --ped $pedigree -m a -m d \
    | bcftools +fill-tags -Ou - -- -t AN,AC,MAF,F_MISSING,HWE,MAC:1=MAC \
    | bcftools +fill-tags -Oz - -- -S $population_ids -t HWE \
    > $annotated_variant_calls
```

#### S3.3 Final QC filter expression and retained site set

The QC-passing site set used for phasing was defined by the following filter logic, applied using bcftools view:

```
bcftools view -e "(TYPE!='snp' && (ABS(ILEN) >= 50)) || \
                ALT=='*' || \
                INFO/VQSLOD < 0 || \
                F_MISSING > 0.05 || \
                INFO/MERR > (INFO/AN * 0.05) || \
                INFO/MAC == 0 || \
                  (INFO/HWE_EUR < 1e-10 && INFO/HWE_AFR < 1e-10 && \
                   INFO/HWE_EAS < 1e-10 && INFO/HWE_AMR < 1e-10 && \
                   INFO/HWE_SAS < 1e-10) || \
                FILTER != 'PASS'" -Oz $annotated_variant_calls \
> $variants_to_phase
```

In summary, variants were excluded if they: (1) were indels with reference-alternate length difference ≥50 bp (beyond reliable short-read calling); (2) had a spanning deletion alternate allele ('*'); (3) had VQSLOD < 0 (GATK quality threshold); (4) had >5% missing genotypes; (5) had Mendelian errors exceeding 5% of non-missing alleles; (6) had MAC = 0 (monomorphic after filtering); (7) failed HWE ($p < 1\times10^{-10}$) in all five superpopulations; or (8) did not pass GATK variant filters. Unlike the GRCh38 1kGP panel, we did not exclude singleton variants, as SHAPEIT5 can phase these.

For the GRCh38 panel comparison, only the subset of filters that were used in generating the original consortium release were applied: removal of symbolic alleles, spanning deletions, indels ≥50 bp, and variants with MAC = 0.

### S4. Ground truth VCF processing

#### S4.1 Pangenome VCF normalization

The combined HGSVC3/HPRC pangenome VCF required preprocessing to ensure consistent variant representation with the 1kGP callsets. Processing was performed separately for autosomes and chromosome X.

For autosomes, the processing pipeline consisted of:

```
bcftools view -r $region -s ^$reference_samples $pangenome_vcf \
| bcftools annotate -x INFO/AT \
| bcftools norm --atomize --atom-overlaps . -m +snps \
```

```
| bcftools norm -f $ref_fasta -m -any \
| bcftools view -i "F_MISSING < 0.05" \
| bcftools +fill-tags -- -t AN,AC,MAF,MAC:1=MAC,MISSING:1=F_MISSING \
| bcftools +setGT -- -t a -n p \
| bcftools view -c 1:minor
```

Reference samples (CHM13 and GRCh38 reference haplotypes) were excluded. The atomization and normalization steps match those applied to the 1kGP callset. Sites with >5% missingness across pangenome samples were excluded. Missing genotypes were converted to phased reference calls using bcftools +setGT -t a -n p because assemblies do not have missing variants. Apparent missing genotypes represent reference alleles. Finally, sites monomorphic after processing were removed.

#### S4.2 Chromosome X ground truth processing

The HGSVC pangenome VCFs represent PAR regions on both chrX and chrY separately, rather than combining them into pseudo-diploid chrX representations as is standard for callsets produced from short-read alignments. Accordingly, chromosome X ground truth analysis was limited to female samples when using HGSVC data.

The HPRC pangenome VCFs used pseudo-diploid PAR representations, allowing us to retain male samples. However, male non-PAR regions were represented as true haploids, requiring conversion to diploid format for comparison with short-read callsets (which typically call male chrX as if diploid).

Both pangenome sources exhibited partial missing genotypes (e.g., '1|.') for samples in PAR regions with large heterozygous deletions. Short-read alignments in these regions would be expected to produce valid diploid calls with reduced coverage (eg, 1|1). Instead of retaining these pangenome variants for comparison to these valid diploid calls, the missing allele was causing these sites to be removed by the missingness filter. To prevent the missingness filter from erroneously filtering out these sites, while still allowing for the filter to remove true missing variants (.|.), we used a sed command to mask diploid-missing genotypes, convert partial missing genotypes to haploid calls (e.g., '0|.' -> '0'), then restore the masked fully missing genotypes. After missingness filtering, bcftools fixploidy was used to convert all haploid variants (both partial PAR variants and male haploid regions in HPRC vcfs) to diploid representation; bcftools setGT was used to phase the newly created homozygous diploid variants (e.g., '0' -> '0|0'). This approach introduced potential phasing errors for non-reference partial missing calls in PAR regions (e.g., '1|.' -> '0|1') but enabled more accurate measures of genotype concordance assessment. The only difference between HGSVC and HPRC chromosome X processing was sample selection: male samples were excluded from HGSVC chromosome X pangenome files, while they were retained for HPRC pangenomes.

This strategy was deployed using the following code block:

```
                      # no -S for HPRC_vcf
bcftools view -r $region -S $female_samples $HGSVC_vcf \
| [same normalization as autosomes] \
| sed 's,\.|\.,qqq,g' | sed 's,\.|,,g' | sed 's,|\.,,g' | sed 's,qqq,\.|\.,g' \
| [same missingness filtering and annotation as autosomes] \
| bcftools +fixploidy -- -f 2 \
| bcftools +setGT -- -t a -n p \
| [same minor allele filtering and sorting as autosomes]
```

### S5. Panel phasing

#### S5.1 Two-stage phasing strategy

Panel phasing was performed using SHAPEIT5's two-stage protocol to optimize accuracy across the allele-frequency spectrum. In the first stage, common variants (MAF ≥ 0.1%) were phased using SHAPEIT5_phase_common. In the second stage, rare variants (MAF < 0.1%) were phased onto the common-variant scaffold using SHAPEIT5_phase_rare in genomic chunks.

#### S5.2 SHAPEIT5_phase_common parameters

As SHAPEIT5_phase_common is a version of SHAPEIT4 that has simply been tuned for large datasets, we followed the SHAPEIT4 website's instructions (https://odelaneau.github.io/shapeit4/) for high-accuracy phasing:

> "By default, the sequence used by SHAPEIT4 is 5b,1p,1b,1p,1b,1p,5m. It provided a good trade-off between speed and accuracy in our experiments. If running time is not an issue, you may consider increasing the number of iterations using --mcmc-iterations 10b,1p,1b,1p,1b,1p,1b,1p,10m for instance.
>
> Reducing the number of conditioning neighbours in the PBWT can be achieved using the --pbwt-depth option. The default value is 4. Decreasing it results in faster runs at the cost of some accuracy. Conversely, increasing it to 8 for instance can lead to better accuracy.
>
> You can also change the size of the genomic window into which phasing is carried out using the option –window... Increasing it results in more conditioning haplotypes being used and therefore increased running time."

We also used an hmm-ne value of 135,000. Early in development we noticed that this parameter could affect panel-wide accuracy (using HPRCv1.1 as a ground truth) by 0.1-0.2%, and after some trial and error we found a value of 135000 optimized accuracy.

| Parameter | Value | Rationale |
|---|---|---|
| --filter-maf | 0.001 | Defines rare variant threshold (0.1%) |
| --mcmc-iterations | 10b,1p,1b,1p,1b,1p,1b,1p,10m | High-accuracy iteration scheme from SHAPEIT4 |
| --pbwt-depth | 8 | Per SHAPEIT4 paper, increasing from default 4 improves haplotype matching |
| --pbwt-mac | 5 | Default value |
| --pbwt-mdr | 0.1 | Default value |
| --pbwt-modulo | 0.1 | Default value |
| --hmm-window | 5 | 5 cM HMM window (increased from default 4, again following SHAPEIT4 'high accuracy' settings) |
| --pbwt-window | 5 | Matches HMM window |
| --hmm-ne | 135,000 | Effective population size optimized for this dataset |

Pedigree information was provided via the --pedigree flag to enable Mendelian pre-phasing of trio samples. This allows SHAPEIT5 to use parental genotypes to constrain phasing in offspring before statistical phasing, improving accuracy for both probands and parents.

A representative command:

```
SHAPEIT5_phase_common \
      --input filtered_variants.bcf \
      --map recombination_map.txt.gz \
      --output common_phased.bcf \
      --thread 16 \
      --filter-maf 0.001 \
      --mcmc-iterations 10b,1p,1b,1p,1b,1p,1b,1p,10m \
      --pbwt-depth 8 \
      --pbwt-mac 5 \
      --pbwt-mdr 0.1 \
      --pbwt-modulo 0.1 \
      --hmm-window 5 \
      --pbwt-window 5 \
      --hmm-ne 135000 \
      --pedigree trios.ped \
      --region chr1
```

#### S5.3 SHAPEIT5_phase_rare parameters

Rare variant phasing was performed using SHAPEIT5_phase_rare with the common-variant output as a scaffold. Due to high memory requirements, rare variant phasing was performed in ~40 Mb chunks with 1 Mb overlap between adjacent chunks. Chunk boundaries were defined using a custom Python script (generate_regions_for_rare_phasing.py) that ensures chunks respect variant boundaries. Default parameter values were used with the exception of an --effective-size 135,000.

Chunk outputs were concatenated using bcftools concat -l (ligate mode) to preserve phase across chunk boundaries.

#### S5.4 Chromosome X Phasing

Chromosome X was processed in three separate regions: PAR1 (chrX:10001-2781479), non-PAR (chrX:2781480-155701382), and PAR2 (chrX:155701383-156030895). PAR regions were phased as diploid using the same parameters as autosomes.

For the non-PAR region, male samples were specified via the --haploids flag with a file listing male sample IDs. The effective population size was scaled to 75% of the autosomal value (Ne = 101,250) to reflect the lower effective population size of X chromosomes:

```
SHAPEIT5_phase_common \
      [previously described parameters] \
      --hmm-ne 101250 \
      --haploids male_samples.txt \
      --region chrX:2781480-155701382
```

#### S5.5 Panel outputs

Final phased panel outputs were produced in BCF format with CSI indices. Two versions were generated: a 3,202-sample panel including all trio members (for maximum phasing accuracy via Mendelian constraints) and a 2,504-sample unrelated panel (for use as an imputation reference where relatedness would be problematic) generated by subsetting the 3,202 member panel.

### S6. Recombination map generation

#### S6.1 pyrho workflow

Population-specific recombination maps were generated from the unphased, QC-filtered T2T-CHM13 genotypes using pyrho. The input VCFs were filtered to remove low-confidence genotype calls prior to recombination mapping, retaining only SNPs. A short-read accessibility mask (https://s3-us-west-2.amazonaws.com/human-pangenomics/T2T/CHM13/assemblies/annotation/accessibility/combined_mask.bed.gz) was applied both before recombination mapping and to the resulting maps.

Historical effective population sizes for each of the 26 1kGP populations were obtained from previously published smc++ demographic inference. VCFs were split into population-specific subsets before running pyrho.

#### S6.2 Hyperparameter optimization

pyrho uses an $\ell 1$ regularization penalty to smooth recombination rate estimates across adjacent windows. The specific penalty parameters (block penalty and window size) were selected for each population by minimizing the L2 norm between simulated and inferred recombination rates. Simulated data were generated based on each population's demographic history inferred with smc++. Population-specific lookup tables were calculated using a Moran population size equal to 150% of the sample size.

#### S6.3 Map scaling and averaging

The cumulative genetic map length estimated by pyrho is sensitive to the chosen hyperparameters and generally underestimates absolute recombination rates compared to pedigree-based methods. To address this, we scaled each population-specific map to match the per-chromosome cumulative lengths of the deCODE trio-based recombination map. Scaling was performed by multiplying all recombination rates by the ratio (deCODE_chromosome_cM / pyrho_chromosome_cM).

To create a unified global map, we computed a sample-size-weighted average recombination rate across all 26 populations at each genomic position. We also provide maps scaled to HapMap2 chromosome lengths for compatibility with tools whose default parameters were tuned using HapMap2.

#### S6.4 Chromosome X recombination mapping

For chromosome X, historical effective population sizes from smc++ were rescaled by 0.75 to reflect the lower effective population size of X chromosomes relative to autosomes. To accommodate pyrho's requirement for diploid input, we created "pseudodiploid" males by randomly pairing haploid male genotypes. Specifically, male samples were randomly shuffled and consecutive pairs were merged to create synthetic diploid samples. pyrho was run on these pseudodiploid VCFs for the non-PAR region. PAR regions were processed as autosomal. As with autosomes, chromosome X maps were scaled to deCODE sex-averaged recombination map lengths prior to averaging.

### S7. Panel evaluation metric definitions

#### S7.1 Variant matching rules

SHAPEIT5_switch was used to evaluate panel accuracy. SHAPEIT5_switch pairs panel and truth variants by chromosome, position, reference allele, and alternate allele. Matching required exact agreement after normalization; variants that could not be matched unambiguously due to representation differences are excluded. Both the reference panel and the pangenome vcfs were left-normalized and monomorphic variants filtered prior to evaluation, effectively restricting to phase-informative heterozygous genotypes present in both panel and truth callsets.

#### S7.2 Switch error rate (SER)

Switch error rate quantifies the frequency of incorrect phase assignments:

$$SER = \frac{number\ of\ switch\ errors}{number\ of\ heterozygous\ sites\ evaluated} \times 100$$

A switch error occurs when the inferred haplotype assignment at a heterozygous site differs from the ground truth assignment, relative to the preceding heterozygous site in that sample. Because switch errors are defined relative to adjacent sites, the denominator equals the number of pairs of consecutive heterozygous sites evaluated.

#### S7.3 Flip error rate (FER)

Flip error rate quantifies the rate of flip error events, defined as blocks of back-to-back switch errors. These are typically caused by genotyping errors rather than phasing failures:

$$FER = \frac{number\ of\ flip\ error\ events}{number\ of\ heterozygous\ sites\ evaluated} \times 100$$

A flip error consists of two consecutive switch errors affecting a single variant. When a heterozygous genotype is incorrect (e.g., reported as 0|1 when true genotype is 1|1), it creates the appearance of two switches flanking the erroneous site.

#### S7.4 True switch error rate (tSER)

True switch error rate is the proportion of heterozygous variants that are switch errors that are not part of a flip event:

$$tSER = \frac{number\ of\ switches - 2 \times number\ of\ flip\ error\ events - total\ number\ of\ consecutive\ flip\ errors}{number\ of\ heterozygous\ sites\ evaluated} \times 100$$

#### S7.5 Genotype discordance rate

Genotype discordance quantifies the fraction of genotype calls that differ from ground truth, ignoring phase:

$$GER = \frac{genotypes\ differing\ from\ truth}{genotypes\ present\ in\ both\ query\ and\ truth} \times 100$$

A genotype was counted as discordant if the unphased genotype (e.g., {0,1}) differed between panel and truth. Partial matches (e.g., panel 0|1 vs truth 1|1) were counted as discordant.

##### S7.6 Stratification and aggregation

Metrics were computed per chromosome. SHAPEIT5_switch reports both per-sample and per-variant statistics. All reports were collected into central per-variant and per-contig, per-sample dataframes using custom python3 scripts. Bcftools query was used to produce tsv files of each biallelic panel's variant properties (MAC, MAF, assembly-unique status, etc) and joined to the per-variant dataframes on the variant ID (%CHROM_%POS_%REF_%ALT). Group-by operations were used to calculate per bin and/or per subset summary statistics. Per-group statistics were always calculated by summing numerator and denominator, then determining the ratio. When reference haplotype panels were used to phase variants, variant allele frequency bins were defined based on panel MAF. Stratifications by genomic context (shared/unshared regions, STR regions, segmental duplications) used the annotations described in Section S10. Regional summaries used 10 kb bins aggregated with rolling windows (Section S9).

#### S8. SHAPEIT5_switch modifications

##### S8.1 Motivation

The standard SHAPEIT5_switch tool counts all haplotype switches without distinguishing flip errors (back-to-back switches typically caused by genotyping errors) from true switch errors that affect regional phase. Because flip errors inflate switch error rates without reflecting true phasing failures, we modified SHAPEIT5_switch to provide enhanced error classification.

##### S8.2 Modified error tracking

The modified version tracks the following error categories:

- n_all_switches: Total number of single-allele switches (equivalent to original switch count)
- n_flips: Number of flip errors, defined as pairs of consecutive switches. Incremented when the current site is a switch AND the immediately prior heterozygous site was also a switch.
- n_consecutive_flips: Three or more consecutive switches, incremented starting at the third consecutive switch
- n_true_switch_errors: Calculated as n_all_switches − (2 × n_flips) − n_consecutive_flips

##### S8.3 Algorithm modification

The original SHAPEIT5_switch examines pairs of adjacent heterozygous sites. Our modified version examines each heterozygous site individually while tracking the error state of the prior and prior-prior sites. This enables detection of consecutive error patterns:

1. For each heterozygous site, determine if it represents a switch relative to ground truth
2. If current site is a switch AND prior site was a switch, increment n_flips
3. If current site is a switch AND both prior AND prior-prior sites were switches, increment n_consecutive_flips
4. Calculate true switches as: n_all_switches − (2 × n_flips) − n_consecutive_flips

The modified SHAPEIT5_switch is available at https://github.com/JosephLalli/shapeit5.

#### S9. Panel-wide error rate estimation

##### S9.1 Sample stratification by trio status

The 100 samples with assembly-based ground truth were stratified by their trio status, which determines whether they underwent Mendelian pre-phasing. Trio probands (n=68) underwent Mendelian pre-phasing using both parents. Trio parents (n=13) additionally underwent Mendelian pre-phasing, though with only their child as a source of information. Non-trio samples (n=19) were only statistically phased. These three populations have different expected error rates due to the additional information provided by Mendelian constraints.

##### S9.2 Weighted average calculation

Because the 3,202-sample panel contains different proportions of each category than the ground truth samples, we calculated weighted average error rates to estimate panel-wide performance. The full panel contains 608 trio probands, 1,195 trio parents, and 1,399 unrelated non-trio samples. Panel-wide error rates were calculated as the average of each sample subset's error rate, weighted by the number of samples of that subset in the whole panel.

#### S10. Genomic region stratification

##### S10.1 Assembly correspondence annotations

Variants were stratified based on alignment status between T2T-CHM13 and GRCh38 reference assemblies:

- Shared genomic regions: T2T-CHM13 sequences with primary alignment to GRCh38, representing regions resolved in both assemblies
- CHM13-unique regions (~182 Mb): Sequences present only in T2T-CHM13 with no primary alignment to GRCh38, encompassing centromeric satellites, segmental duplications, acrocentric short arms, and other complex sequences that were unresolved or hard-masked in GRCh38
- GRCh38-unique regions (~1.2 Mb): GRCh38 sequences lacking alignment to T2T-CHM13, typically representing assembly errors or false duplications corrected in T2T-CHM13

Region annotations were obtained from chm13v2-unique_compared_to_hg38.bed (https://github.com/marbl/CHM13) for CHM13-unique regions and hg38.GCA_009914755.4.synNet.summary.bed.gz (https://hgdownload.soe.ucsc.edu/goldenPath/hs1/vsHg38/) for assembly correspondence. Variants were annotated using bcftools annotate -a regions.bed --mark-sites +SYNTENIC.

##### S10.2 Repetitive element stratification

Variants were additionally stratified by overlap with repetitive genomic elements. GIAB STR regions were defined using GIAB v3.6 AllTandemRepeats stratification bed files for each reference assembly, and Platinum STR regions were defined using tandem repeat annotations from the Platinum Genomes Project, which cover a larger proportion of the genome and are available in both GRCh38 and CHM13v2.0 coordinates. Segmental duplication annotations, also available in both coordinates, were obtained from Vollger et al. 2022[42].

##### S10.3 Variant complexity stratification

Variants were additionally classified based on whether their allelic representations overlapped other variants. Isolated variants are defined as variants whose reference allele span does not overlap any other variant position. Variants sharing genomic positions with other variants, or whose reference span overlaps positions with multiple variants, were termed overlapping variants.

This stratification distinguishes simple variants from those in complex, multiallelic regions where representation differences between callsets may inflate apparent error rates.

#### S11. Rolling window error rate calculation

Regional error rate profiles displayed in Figure 6 were calculated using a rolling window approach. Each reference genome was first divided into non-overlapping 10 kb bins. For each bin, we then counted the number of switch errors, heterozygous sites evaluated, and genotype calls across all samples. A rolling window centered on 1 bin, flanked by ±24 bins from each center bin, was applied (50 bins per window total = 500 kbp), and window-level error rates were calculated as the sum of errors divided by the sum of denominators across all bins in the window.

Results were stored in parquet format and visualized using karyoploteR[71] in R.

#### S12. Out-of-panel phasing evaluation

To evaluate phasing accuracy for samples not included in the reference panel (simulating typical use cases), we created an out-of-panel evaluation framework. The 78 parents of HPRC-assembled samples were removed from the 2,504-unrelated panel, yielding a 2,426-sample "non-HPRC" panel. This panel was then used to phase the 39 HPRC samples' 1kGP genotypes. We then evaluated phasing accuracy against the HPRCv1.1 assembly-derived ground truth vcf.

Note that SHAPEIT5_phase_rare could not be applied to this 39-sample dataset because the minimum possible MAF (1/78 = 1.3%) exceeds the rare variant threshold, and the rare variant algorithm is not designed for common variants.

#### S13. Cytoband and CNV disorder-stratified analyses

Cytoband-level switch error and genotype discordance rates were computed by aggregating per-variant error counts within each cytoband. Cytoband coordinates were obtained from UCSC genome annotation tracks for each reference assembly.

CNV disorder region coordinates were obtained from the DECIPHER database (https://www.deciphergenomics.org/) and used to subset per-variant error statistics for regions associated with recurrent copy number variation syndromes. Coordinates were converted between assemblies as needed using the Broad liftover website.

#### S14. Comparison with double-strand break initiation sites

To compare LD-based recombination hotspots with experimentally measured meiotic double-strand break initiation sites, we reanalyzed DMC1 ChIP-SSDS data from Pratto et al. 2014[83].

Reads were aligned to T2T-CHM13v2.0 using bwa-mem2[82] and processed using a Nextflow pipeline implementing the Pratto et al. single-stranded DNA sequencing (ssds) analysis protocol (nf-core/ssds)[84,85]. Duplicate reads were marked and low-quality alignments were filtered. Peaks were called using MACS2[86,87] (v2.2.9.1) with parameters appropriate for ChIP-SSDS data.

Consistent with the original publication, peaks from across individuals were combined, and overlapping peaks were merged. Peak sets were stratified by donor PRDM9 genotype: peaks from individuals with AA/AB PRDM9 genotypes were combined to define PRDM9-A/B peaks. PRDM9-C peaks were obtained by subtracting the PRDM9-A/B peak set from peaks called in the PRDM9-A/C genotype individual.

Hotspot-peak overlap was quantified by intersecting hotspot intervals with αDMC1 peak intervals using bedtools. Enrichment was assessed by comparing observed overlap to a null distribution generated by shuffling hotspot intervals within chromosomes 10,000 times while preserving interval lengths and excluding assembly gaps.

### S15. Imputation benchmarking

#### S15.1 Pseudo-array construction

To simulate array-based genotyping workflows, SGDP variant calls were downsampled to sites present on the Illumina Omni 2.5 genotyping array. T2T-CHM13 Omni 2.5 array site coordinates were obtained from the GATK resource bundle (https://s3-us-west-2.amazonaws.com/human-pangenomics/index.html?prefix=T2T/CHM13/assemblies/variants/GATK_CHM13v2.0_Resource_Bundle/), which contains array sites lifted from GRCh38 to CHM13 coordinates.

Downsampled genotypes were filtered to remove sites with over 5% of genotypes missing, HWE $\leq 1\times10^{-4}$, and a MAF ≤1%.

After filtering, variants that were placed in phase sets by HaplotypeCaller were set to unphased using bcftools +setGT -t a -n u:

```
bcftools isec -n=2 -w 1 sgdp_variants.bcf omni_sites.vcf.gz \
| bcftools +fill-tags -- -t F_MISSING,MAF,HWE \
| bcftools view -e "F_MISSING>=0.05 || MAF<=0.01 || HWE<=1e-4" \
| bcftools +setGT -- -t a -n u \
> sgdp_downsampled.bcf
```

#### S15.2 Pre-phasing

Downsampled pseudo-array genotypes were pre-phased against the 2,504-unrelated-sample 1kGP reference panel using SHAPEIT5_phase_common in reference mode. Pre-phasing used different parameters than panel phasing to reflect array pre-phasing defaults (pbwt-modulo 0.02) and IMPUTE5 default settings (1000000 Ne)

```
SHAPEIT5_phase_common \
      --input sgdp_downsampled.bcf \
      --reference 1kgp_panel.bcf \
      --map recombination_map.txt.gz \
      --output sgdp_prephased.bcf \
      --pbwt-modulo 0.02 \
      --hmm-ne 1000000 \
      --region chr1
```

#### S15.3 Imputation with IMPUTE5

Imputation was performed using IMPUTE5 (v1.2.0) with the pre-phased target haplotypes and phased reference panel:

```
impute5 \
      --g sgdp_prephased.bcf \
      --h 1kgp_panel.bcf \
      --m recombination_map.txt.gz \
      --r chr1 \
      --buffer-region chr1 \
      --out-ap-field \
      --o sgdp_imputed.bcf
```

The entire chromosome was specified as both the imputation region (--r) and buffer region (--buffer-region) to avoid edge effects. Allelic probabilities were output (--out-ap-field) for downstream accuracy assessment.

##### S15.4 Imputation accuracy metrics

Imputation accuracy was quantified using GLIMPSE2_concordance with the following settings:

```
GLIMPSE2_concordance \
     --gt-val \
     --bins "0 0.00021 0.00042 0.00064 0.001 0.0016 0.0022 0.003 0.004 \
             0.0054 0.0072 0.0094 0.0126 0.0172 0.0244 0.0369 0.0601 \
             0.1018 0.1661 0.2556 0.3724 0.5" \
     --af-tag MAF \
     --out-r2-per-site \
     --input concordance_input.txt \
     --output concordance_results
```

The --gt-val flag specifies ground truth validation mode. MAF bins were chosen to provide approximately log-linear spacing with roughly equal numbers of variants per bin, with a separate singleton bin (MAF 0-0.00021).

Imputation $r^2$ was computed by GLIMPSE2_concordance. Metrics were computed within each MAF bin and stratified by variant type (SNP vs indel) and genomic context. Additional GLIMPSE2_concordance runs were performed per ancestry bin.

##### S15.5 Cross-assembly imputation comparison

To compare imputation accuracy between reference assemblies on equivalent variant sets, we used LiftoverIndel (Section S15.2) to convert the T2T-CHM13 panel to GRCh38 coordinates. This enabled imputation of GRCh38-coordinate SGDP genotypes using either the native GRCh38 1kGP panel or the lifted T2T-CHM13 panel, with accuracy assessed against the same ground truth variants.

#### S16. Removing singleton variants from reference haplotype panels

For downstream analyses in which users prefer to exclude singleton variants from the reference haplotype panel, singleton sites (minor allele count = 1 within the panel) can be removed prior to imputation using bcftools. Specifically, we retained only variants with allele count ≥2 using:

```
bcftools view -c 2 -Ob $panel > no_singletons_panel.bcf && \
bcftools index no_singletons_panel.bcf
```

This code will output an indexed BCF containing only non-singleton sites.

#### S17. Data and code availability

##### S17.1 Generated resources

- T2T-CHM13 recombination maps: https://zenodo.org/records/14891074
- T2T-CHM13 phased 1kGP panel: https://s3-us-west-2.amazonaws.com/human-pangenomics/index.html?prefix=T2T/CHM13/assemblies/variants/1000_Genomes_Project/chm13v2.0/Phased_SHAPEIT5_v1.1

#### S17.2 Analysis code

- Phasing and evaluation scripts: https://github.com/JosephLalli/phasing_T2T
- Recombination mapping scripts: https://github.com/andrew-bortvin/1kgp_chm13_maps
- Modified SHAPEIT5_switch: https://github.com/JosephLalli/shapeit5
- LiftoverIndel: https://github.com/JosephLalli/LiftoverIndel
- Docker container with all dependencies: jlalli/phasing_T2T on Docker Hub

#### S17.3 External resources used

- 1kGP T2T-CHM13 unphased callset: https://s3-us-west-2.amazonaws.com/human-pangenomics/index.html?prefix=T2T/CHM13/assemblies/variants/1000_Genomes_Project/chm13v2.0/all_samples_3202/
- 1kGP GRCh38 phased panel: https://ftp.1000genomes.ebi.ac.uk/vol1/ftp/data_collections/1000G_2504_high_coverage/working/20220422_3202_phased_SNV_INDEL_SV/
- Combined HGSVC3/HPRC pangenome (CHM13 coordinates): https://ftp.1000genomes.ebi.ac.uk/vol1/ftp/data_collections/HGSVC3/release/Graph_Genomes/1.0/2024_02_23_minigraph_cactus_hgsvc3_hprc/
- SGDP variant calls: https://s3-us-west-2.amazonaws.com/human-pangenomics/index.html?prefix=T2T/CHM13/assemblies/variants/SGDP/chm13v2.0/
- T2T-CHM13 GATK resource bundle: https://s3-us-west-2.amazonaws.com/human-pangenomics/index.html?prefix=T2T/CHM13/assemblies/variants/GATK_CHM13v2.0_Resource_Bundle/
- T2T-CHM13 accessibility mask: https://s3-us-west-2.amazonaws.com/human-pangenomics/T2T/CHM13/assemblies/annotation/accessibility/combined_mask.bed.gz
- 1kGP pedigree file: http://ftp.1000genomes.ebi.ac.uk/vol1/ftp/data_collections/1000G_2504_high_coverage/1000G_698_related_high_coverage.ped

Chain files: https://s3-us-west-2.amazonaws.com/human-pangenomics/T2T/CHM13/assemblies/chain/

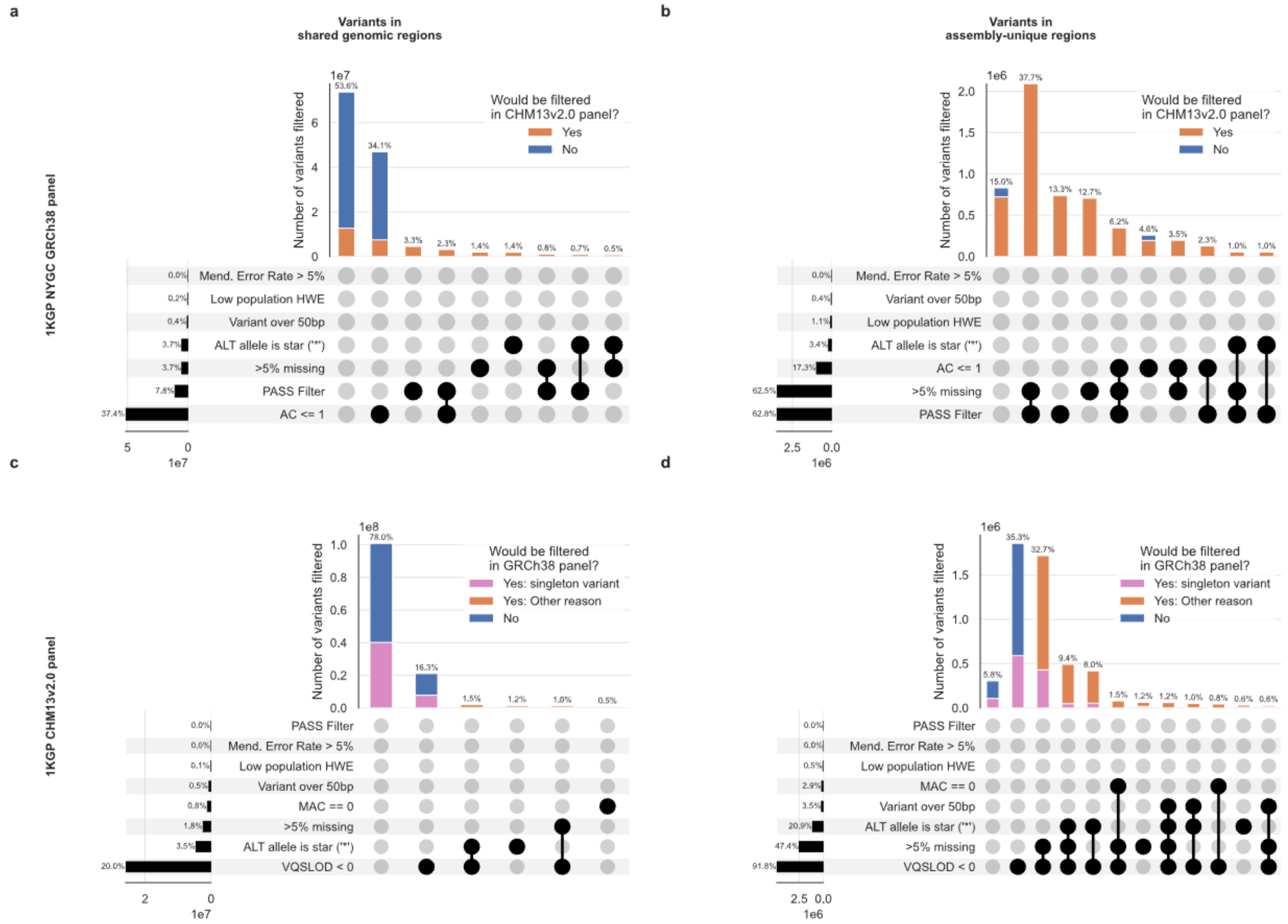


**Supplemental Figure 1. Impact of variant filters on inclusion of syntenic and nonsyntenic variants in the 1KGP GRCh38 reference haplotype panel and the 1KGP T2T-CHM13 reference haplotype panel.**

a) Upset plot showing the number of unfiltered 1KGP GRCh38 panel variants in regions of the genome that are syntenic with T2T-CHM13 that meet different kinds of variant filters. The bar plot portion of the upset plot is colored to illustrate the proportion of GRCh38 variants that would have been filtered by this paper's variant filters. b) Upset plot showing the same information for unfiltered 1KGP T2T-CHM13 variant calls in regions of T2T-CHM13 that are syntenic with GRCh38. The color scheme is used to illustrate the proportion of T2T-CHM13 variant calls that would have been filtered out by the NYGC GRCh38 panel variant filter settings. c) Upset plot displaying the same information for 1KGP GRCh38 variant calls that are in regions of GRCh38 that are nonsyntenic with T2T-CHM13. d) Upset plot displaying the same information for 1KGP T2T-CHM13 variant calls that are in regions of T2T-CHM13 that are nonsyntenic with GRCh38. Variant filter combinations were plotted if they contained at least 0.5% of all variants in the indicated variant subset. Note that only variants that meet VQSR PASS thresholds are available online.

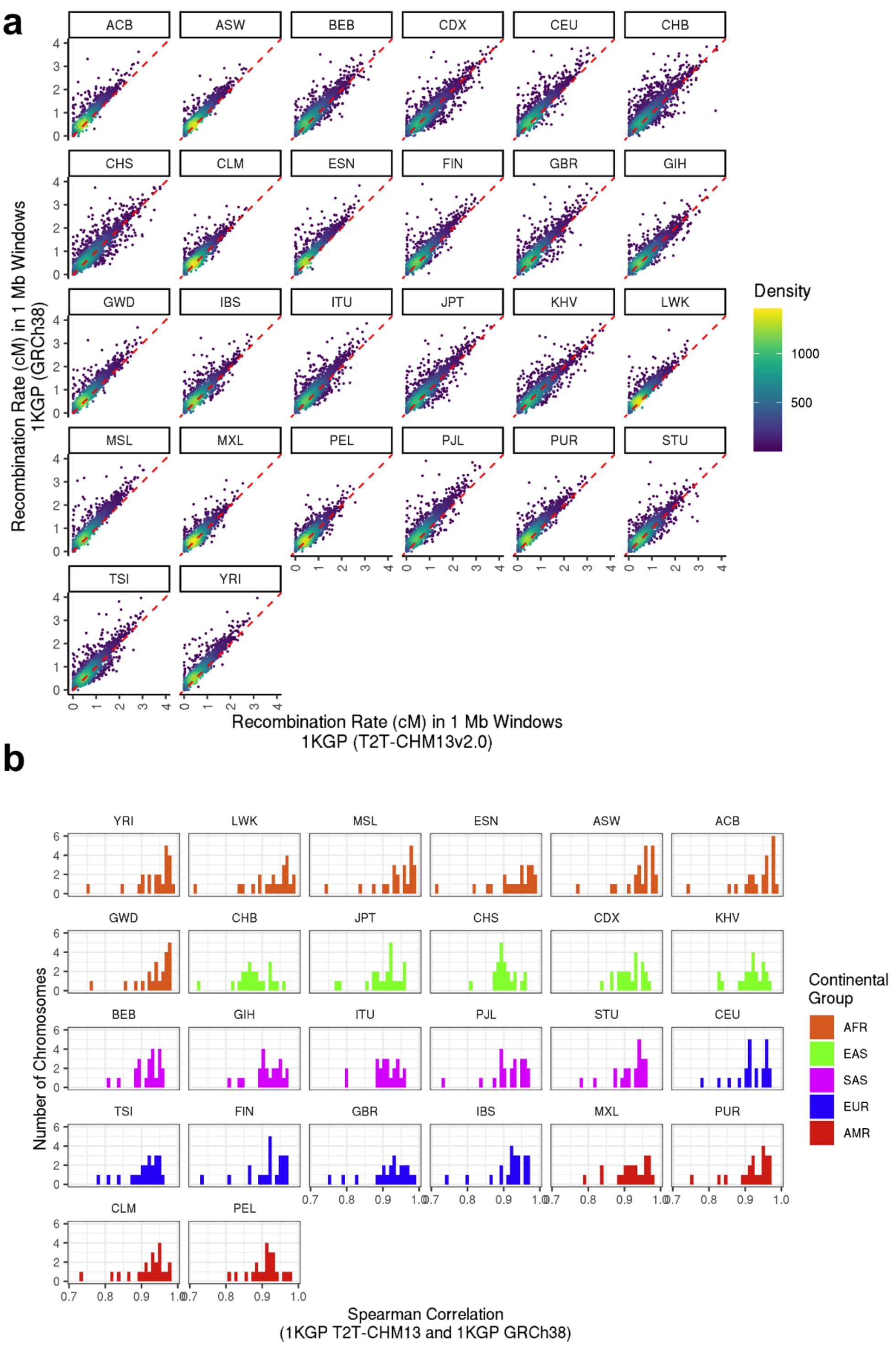


**Supplemental Figure 2. Comparison of T2T-CHM13 and GRCh38 1KGP Recombination Maps**

A). Recombination rate in 1 Mb windows in maps generated from reads aligned to GRCh38 and T2T-CHM13. GRCh38 data is taken from Spence and Song (2019), lifted over to the T2T-CHM13 reference. No scaling has been applied to the maps in this figure. B). Spearman correlation in 1 Mb windows T2T-CHM13-native recombination maps generated from the realigned 1KGP dataset and GRCh38 maps from Spence and Song (2019), lifted over to T2T-CHM13. Correlations are calculated separately per chromosome.

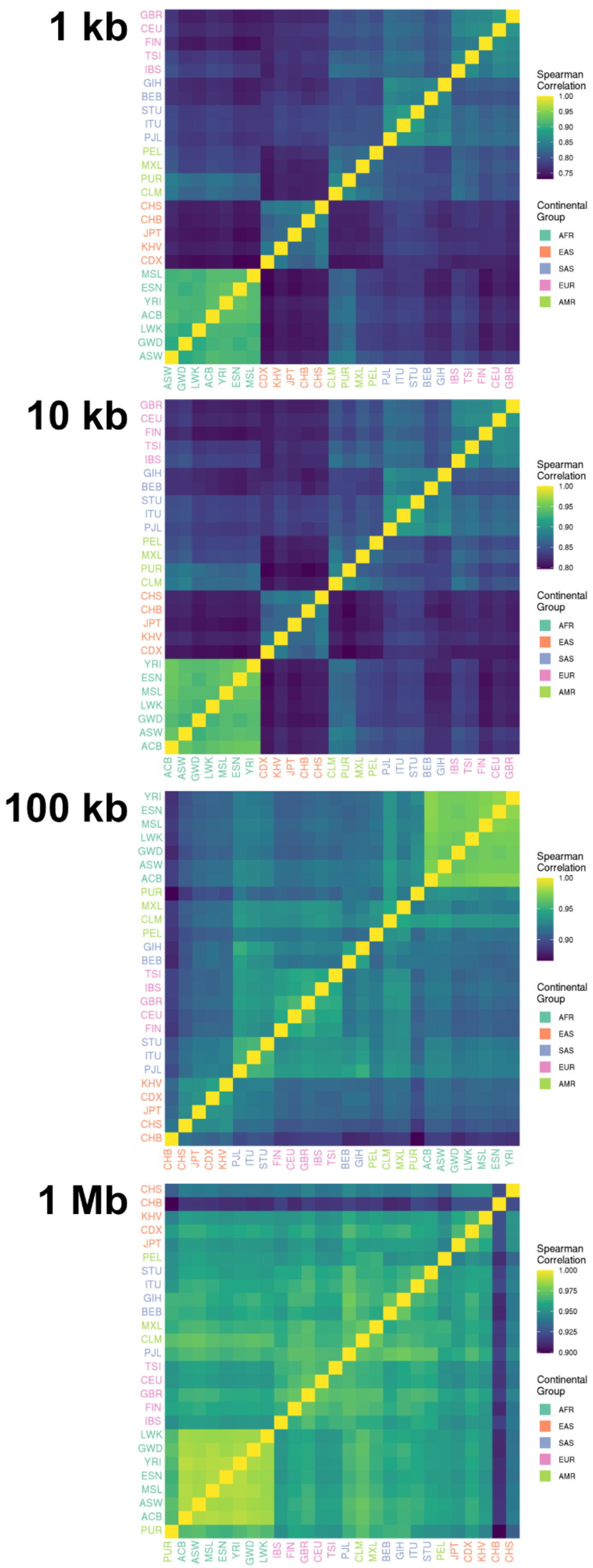


**Supplemental Figure 3. Comparison of T2T-CHM13-native recombination maps between populations.**

Genome-wide Spearman correlations between populations within the T2T-CHM13-native recombination maps generated from 1KGP data. Spearman correlation is calculated at various resolutions between 1 kb and 1 Mb.

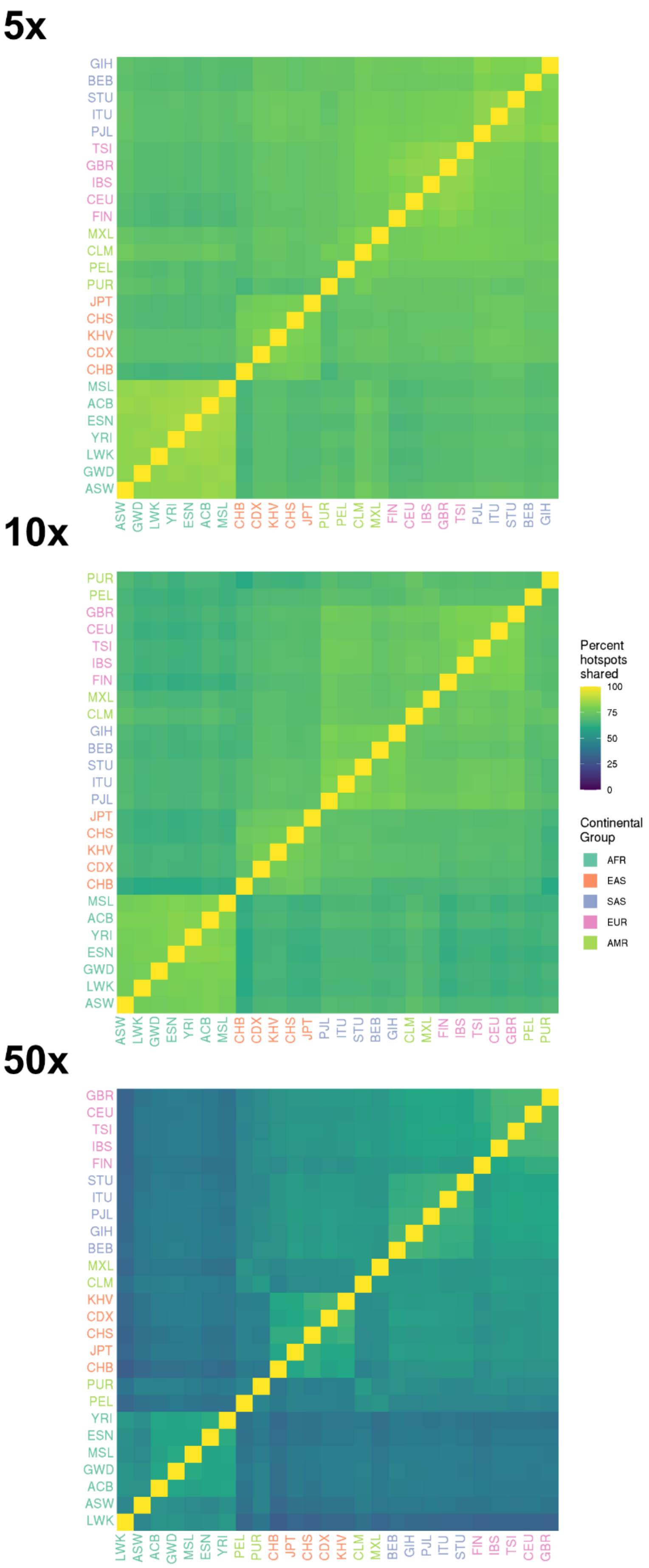


**Supplemental Figure 4. Comparison of recombination hotspots in T2T-CHM13-native recombination maps between populations.**

Percent of hotspots overlapping between populations within the T2T-CHM13-native recombination maps generated from 1KGP data. Hotspots were called at three thresholds - 5x the genome-wide average recombination rate, 10x (as in Halldorsson et al. (2019)[3]), and 50x.

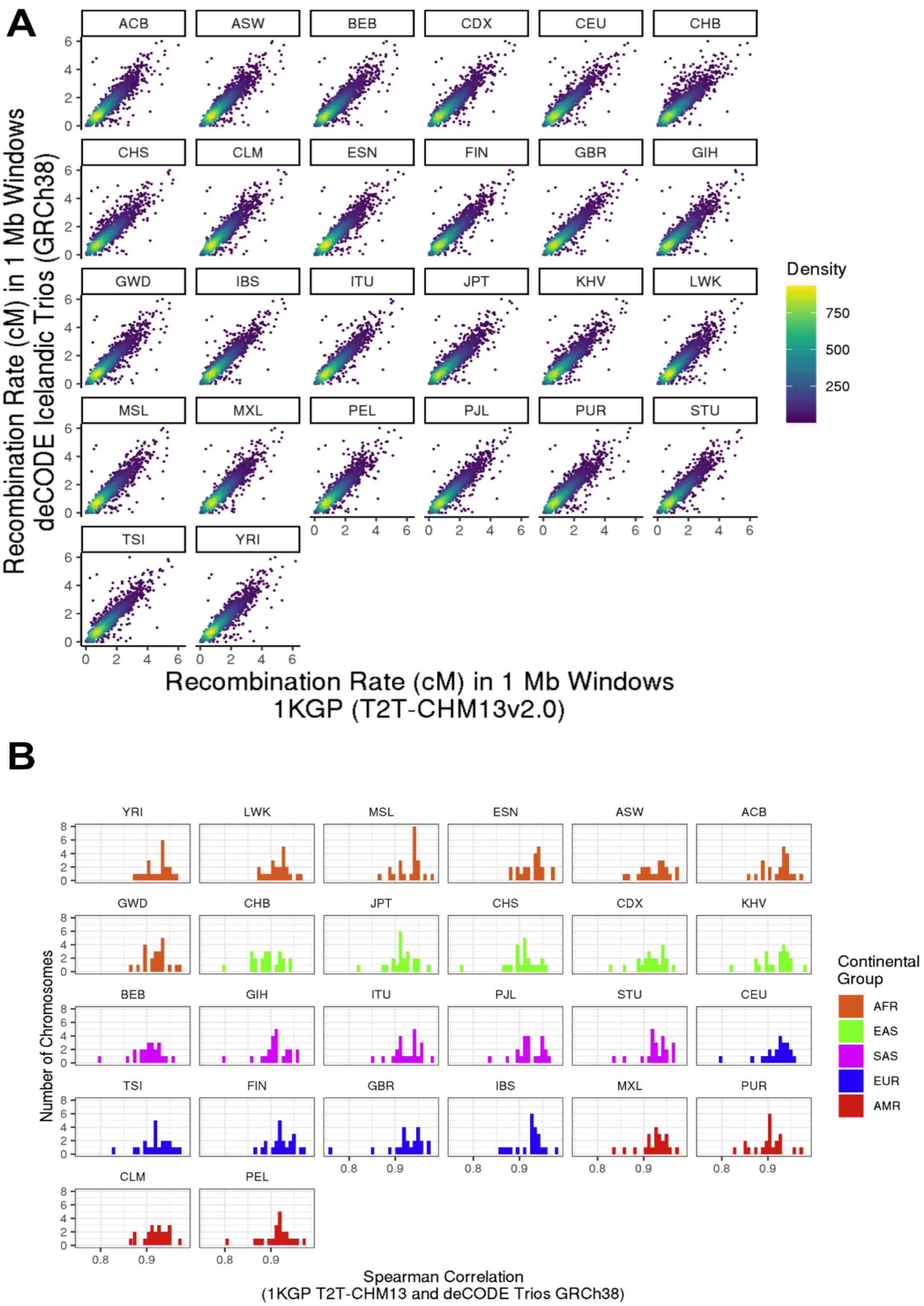


**Supplemental Figure 5 Comparison of T2T-CHM13-native maps to trio-based deCODE recombination maps.**

A). Recombination rate in 1 Mb windows in T2T-CHM13-native LD-based maps generated from 1KGP data and GRCh38 trio-based maps from Icelandic individuals from Halldorsson et al. (2019)[30] lifted over to T2T-CHM13. B). Spearman correlation in 1 Mb windows T2T-CHM13-native recombination maps generated from the realigned 1KGP dataset and trio-based maps from Halldorsson et al. (2019), generated using GRCh38 and lifted over to T2T-CHM13. Correlations are calculated separately per chromosome.

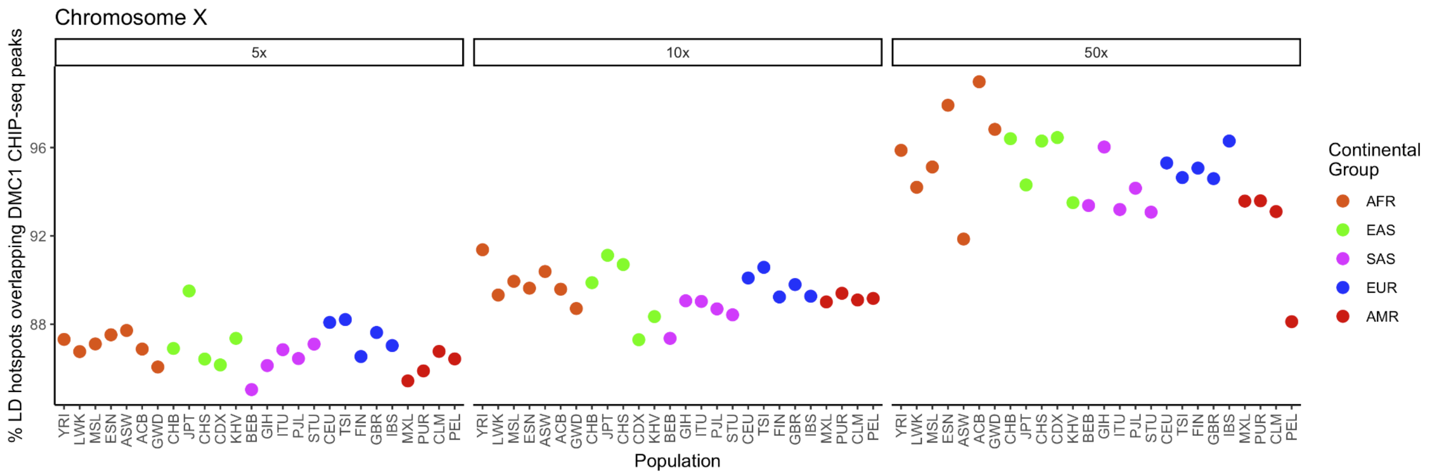


**Supplemental Figure 6 Comparison of recombination hotspot locations in T2T-CHM13-native recombination maps with ChIP-seq data on Chromosome X.**

Percent of recombination hotspots in T2T-CHM13-native LD-based recombination maps on chromosome X that overlap with aDMC1 peaks from the Pratto et al (2014) dataset, realigned to T2T-CHM13. Recombination hotspots are called at three thresholds - 5x the genome-wide recombination rate, 10x (as in Halldorsson et al (2019)), and 50x.

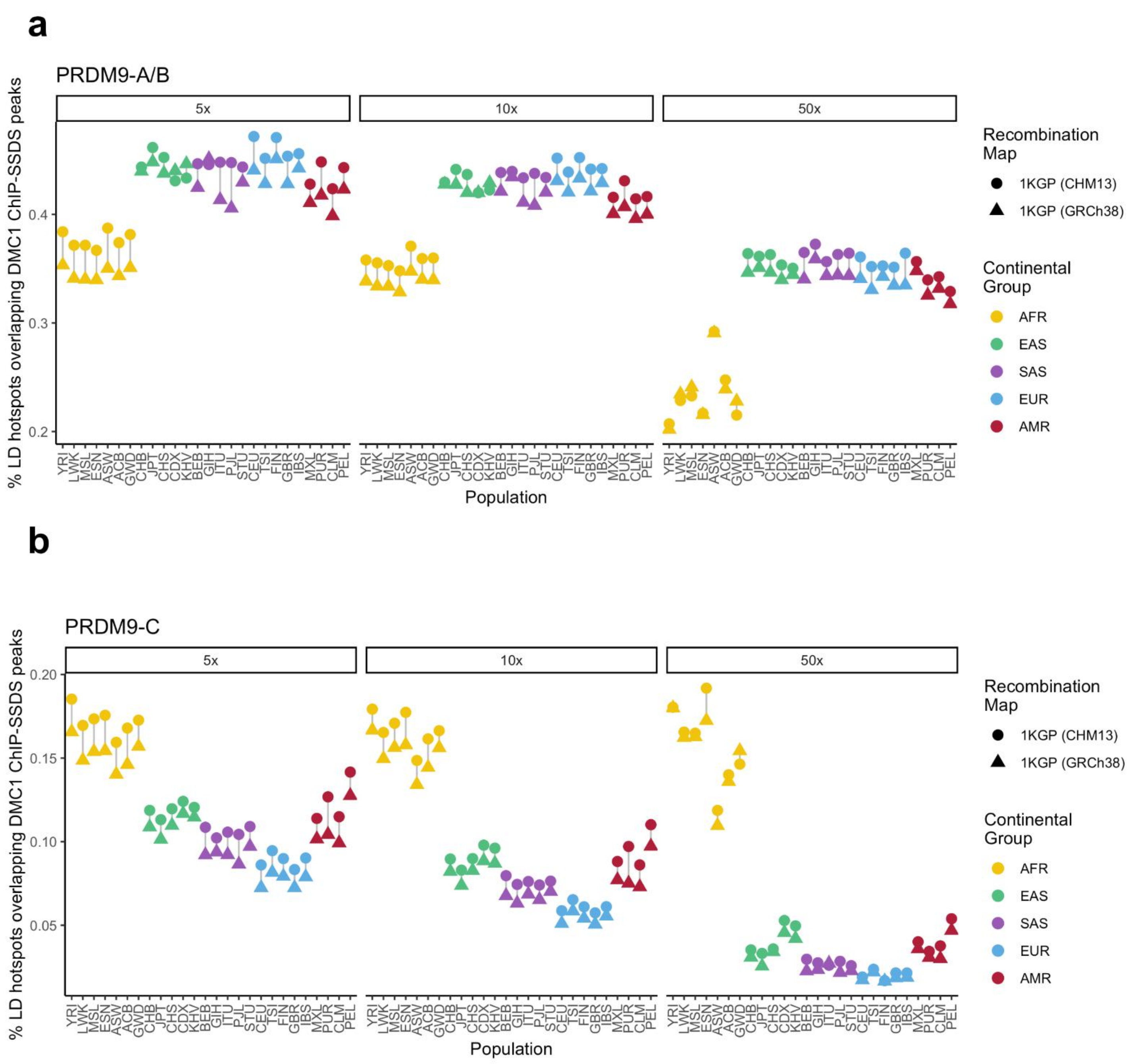


**Supplemental Figure 7 Comparison of recombination hotspot locations in T2T-CHM13-native recombination maps with ChIP-seq data, stratified by PRDM9 genotype.**

Proportion of autosomal LD hotspots that overlap anti-DMC1 ChIP peaks from Pratto et al (2014), subset by the genotype of individuals in which the ChIP-seq peaks are present. A) Is subset to ChIP-seq peaks that do not occur in individuals carrying the PRDM9-C allele B) Is subset to ChIP-seq peaks that occur exclusively in individuals carrying the PRDM9-C allele

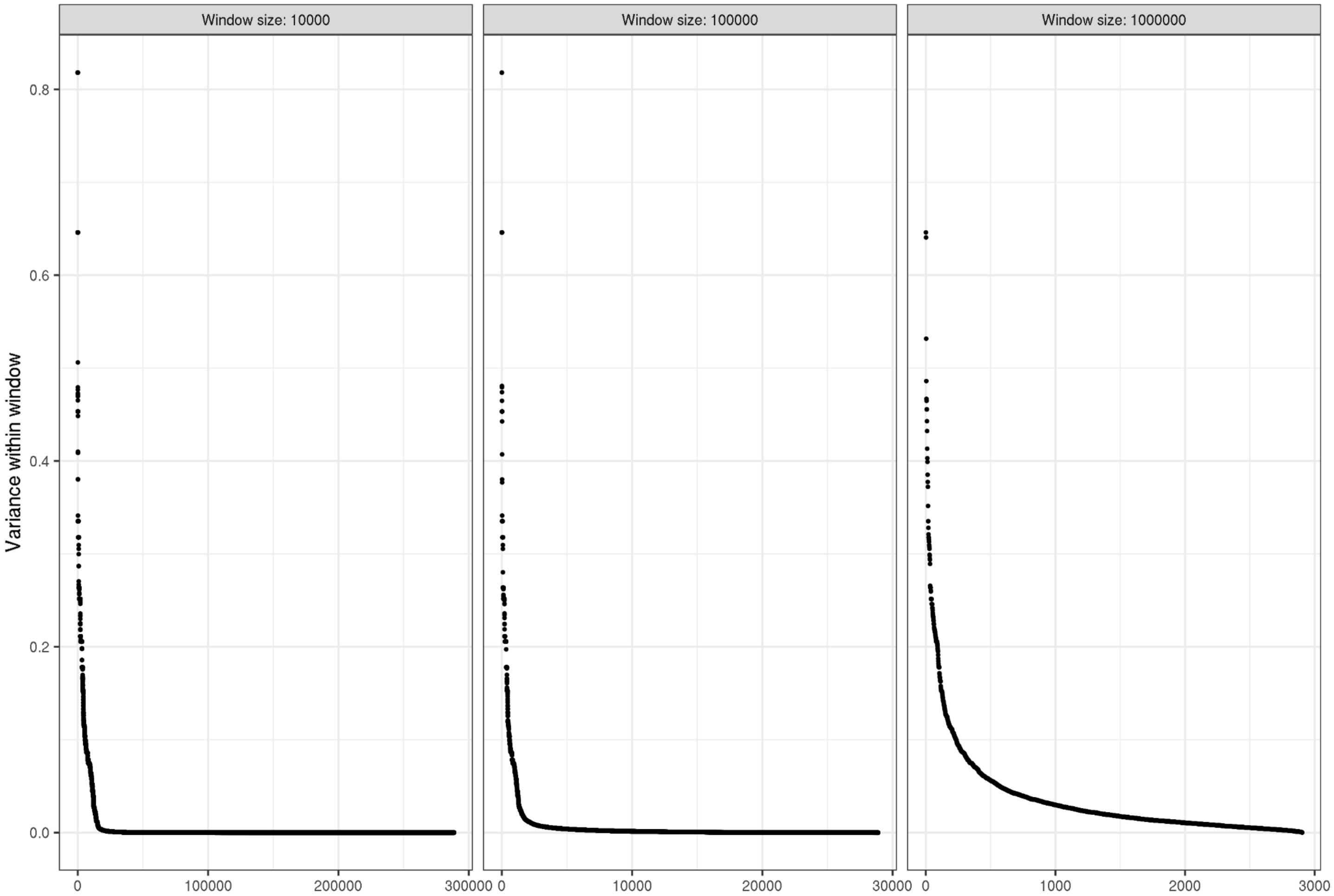


**Supplemental Figure 8. Variance in recombination rate across populations**

Recombination maps from all 26 populations were binned with a bin size of a) 10 kb, b) 100 kb, and c) 1 Mb. The variance of measured recombination rates in each bin was calculated and plotted (one dot per bin), with bins ranked and sorted from highest to lowest measured variance.

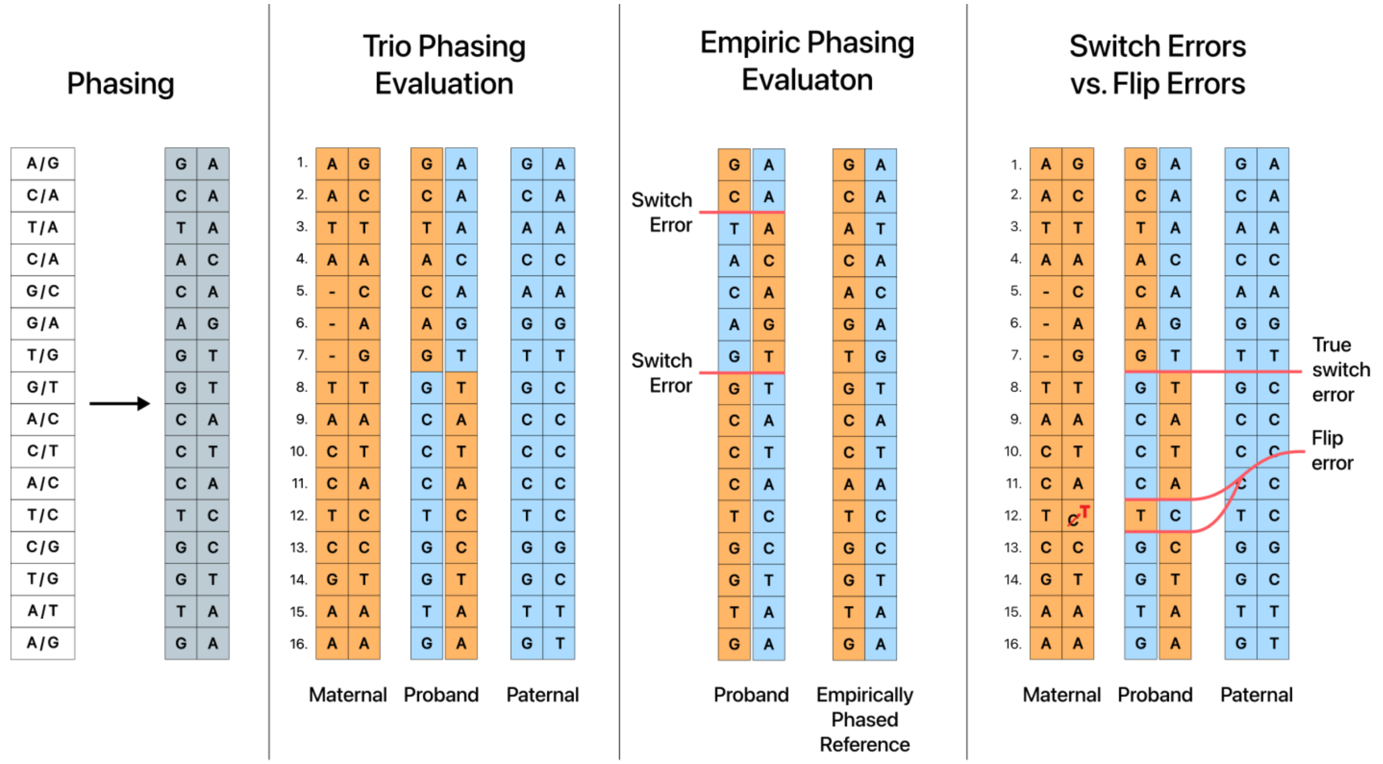


**Supplemental Figure 9. Illustration describing different methods of evaluating phasing quality.**

Phasing is the process of assigning heterozygous variant calls to one of two haplotypes (in the case of diploid organisms.) One of the challenges of assessing the accuracy of statistically phased haplotypes is identifying a ground truth to compare to. In the case of trio-based phasing evaluation, the ground truth is parental genomes. While not every heterozygous site can be assigned to a haplotype of paternal or maternal origin, most can confidently be assigned to one or the other. Alternatively, one can use an empirically phased reference assembly as a source of ground truth.

When a callset's phasing describes a variant that is from a different parent of origin than the preceding variant, that is called a switch error. Genotyping errors, as illustrated in the fourth panel, can force statistical phasing software to assign a variant to a haplotype that is different from nearby heterozygous sites. When two switch errors occur back-to-back, that is described as a flip error. Importantly, flip errors do not affect the phasing of surrounding heterozygous sites. Discerning flip errors from true switch errors is important when evaluating phasing methods, as the majority of switch errors can frequently be found in flip errors. Counting these switches as "true" switch errors inflates switch error rates by including sites that do not affect regional phasing.

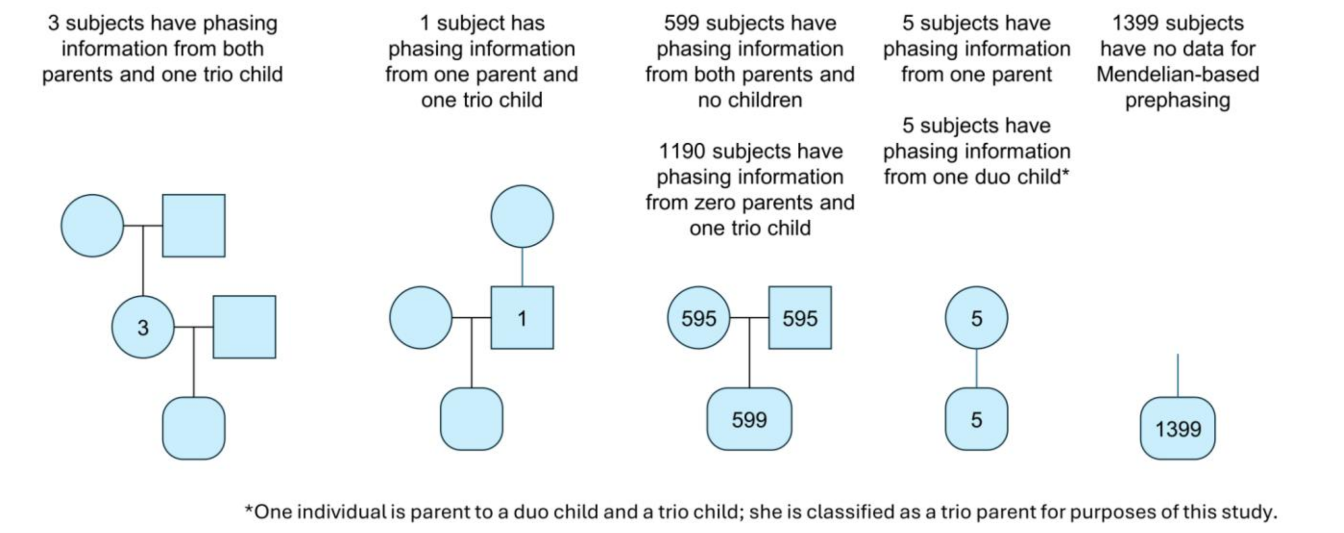


**Supplemental Figure 10. 1000 Genomes Project samples by number of relatives available to provide information for Mendelian pre-phasing.**

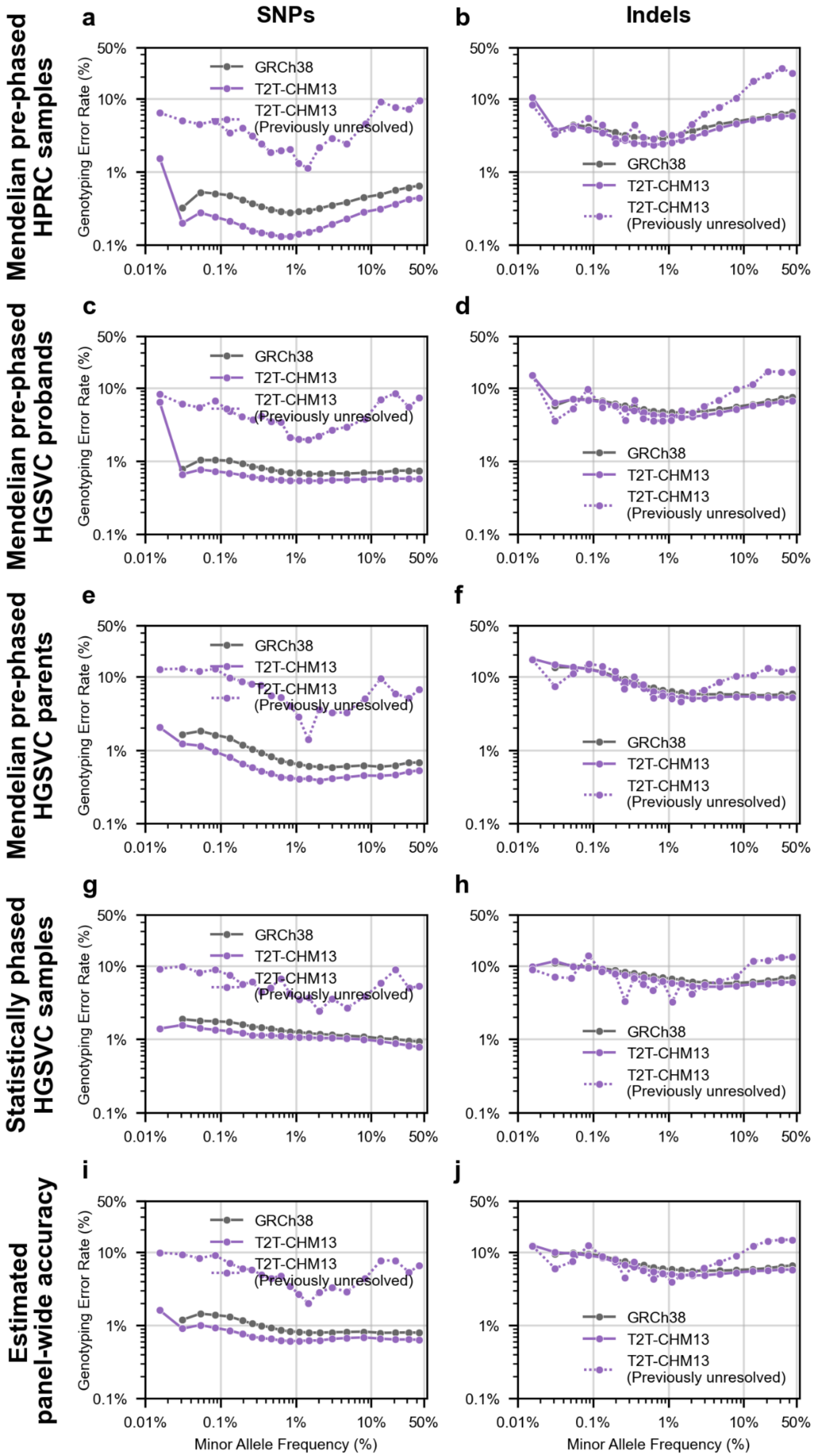


**Supplemental Figure 11. SNP (L column) and Indel (R column) genotyping error rates for 1KGP phased haplotype panel variation, binned by minor allele frequency.**

Genotyping error rates were defined as the number of inaccurate heterozygous or homozygous alternative genotypes (both alleles must be called correctly; there was no partial credit for correctly calling one allele) divided by the number of heterozygous or homozygous alternative genotypes. Correct and incorrect variant calls were determined via comparison to ground-truth assemblies for the following subsets of samples: a) Mendelian-corrected trio probands with ground truth assemblies present in the HPRC human pangenome; b) Mendelian-corrected trio probands with ground truth assemblies produced by the HGSVC; c) Mendelian-corrected trio parents with ground truth assemblies produced by the HGSVC; d) Uncorrected samples which are not part of a 1KGP trio with ground truth assemblies produced by the HGSVC. e) The genotype error rates of trio probands, trio parents, and non-trio samples were weighted by each category's prevalence in the 1000 Genomes Project to produce estimated panel-wide average genotype error rates. Genotype error rates and average minor allele frequency per bin are displayed on a log scale.

a

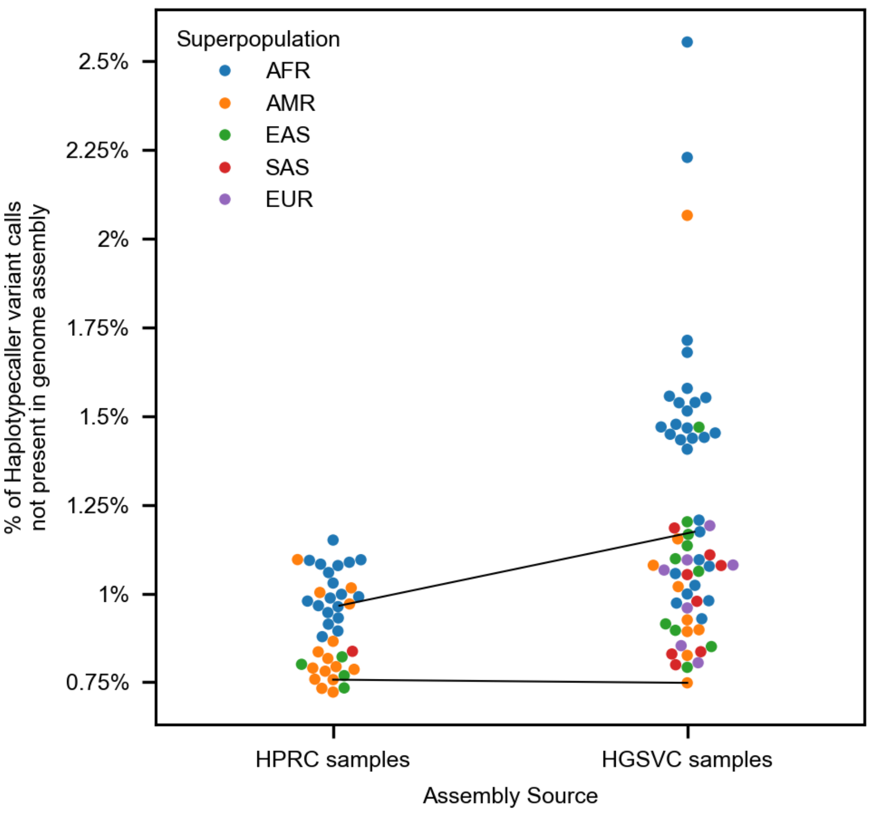


**Supplemental Figure 12: Concordance of GATK Haplotypecaller variant calls with genomic assembly-derived reference variants**

A) Genotype variants from the T2T-CHM13 reference haplotype panel were compared to variants derived from a joint HPRC-HGSVC pangenome released by the HGSVC. Each point is one sample-assembly pair. Two samples had assemblies generated by both the HPRC and the HGSVC: HG00733 and HG02818. The lower line connects the HG00733 datapoints, and the upper line connects the HG02818 data points. Datapoints are colored by their 1KGP assigned superpopulation group.

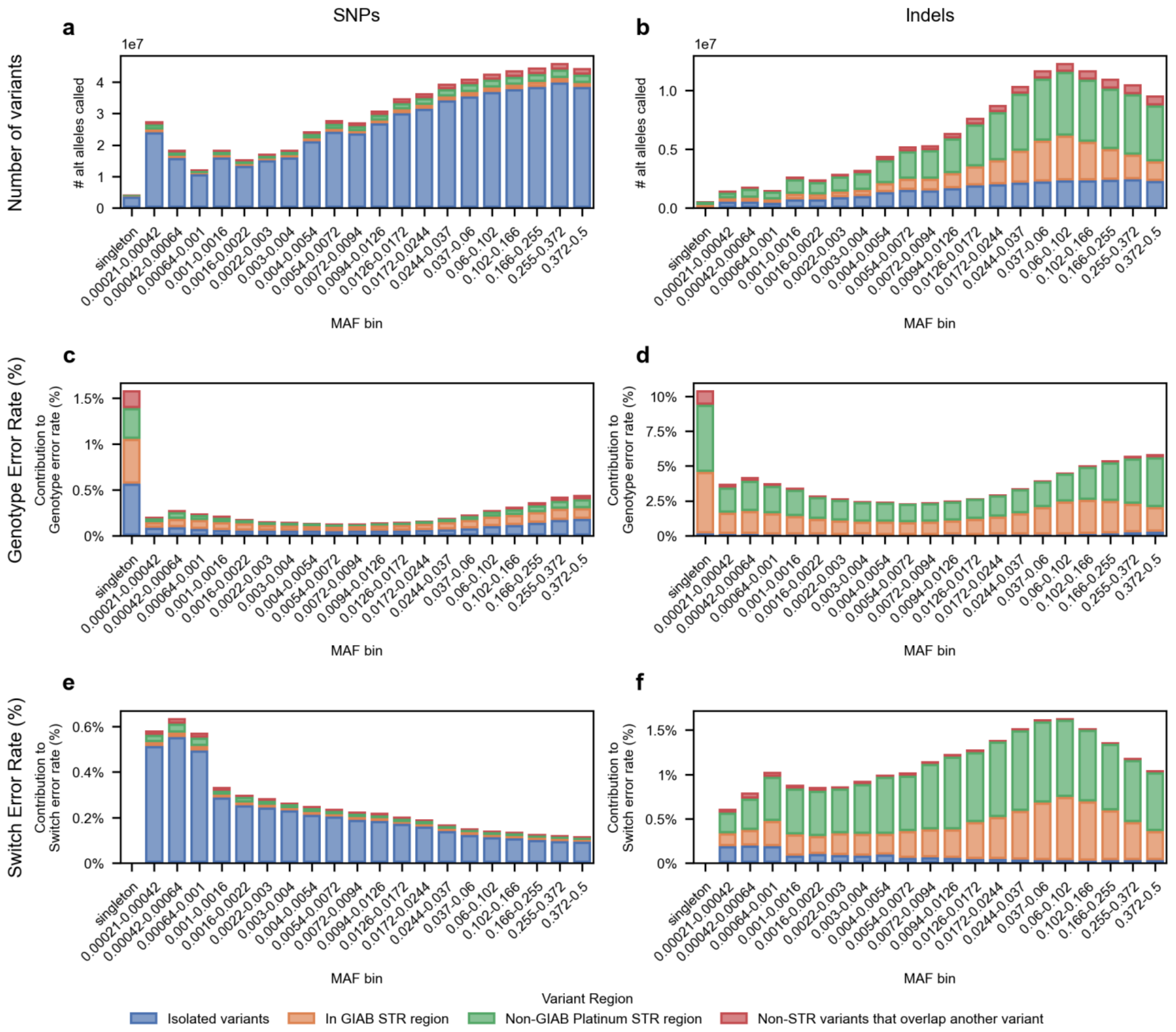


**Supplemental Figure 13: Properties of called SNPs and Indels in increasing complex genetic regions.**

Variants from 39 samples were extracted from this paper's T2T-CHM13 panel. These variants were compared to those samples' HPRCv1.1 assemblies to identify switch errors and genotyping errors, and then stratified by region complexity and minor allele frequency. "GIAB STR region" variants were present in short tandem repeat (STR) regions as defined by the GIAB project (Dwarshuis 2024). "Non-GIAB Platinum STR region" variants are additional variants that are in STR regions as defined by the Platinum Genomes Project (Porubsky 2025), which covers a much larger proportion of the genome. "Non-STR variants that overlap another variant" are defined as additional 1KGP variants that are not in either of the two annotated STR regions, but are overlapped by at least one other variant in the 1KGP callset. "Isolated Variants" are variants that are not in any of the other three variant sets. a) The total number of SNP non-reference T2T-CHM13 genotype calls in 39 HPRC samples, stratified by region complexity and minor allele frequency bin. b) The total number of indel non-reference T2T-CHM13 genotype calls in 39 HPRC samples, stratified by region complexity and minor allele frequency bin. c,d) The contribution of c) SNP and d) Indel variation in each region to the overall SNP or Indel genotype error rate in that bin. The number of SNP or Indel genotype errors in each region-MAF bin was normalized to the genome-wide number of alt variant calls in that bin, ensuring that the height of each bar adds up to the total bin genotyping error rate. e,f) The contribution of c) SNP and d) Indel variation in each region to the overall SNP/Indel switch error rate in that bin.

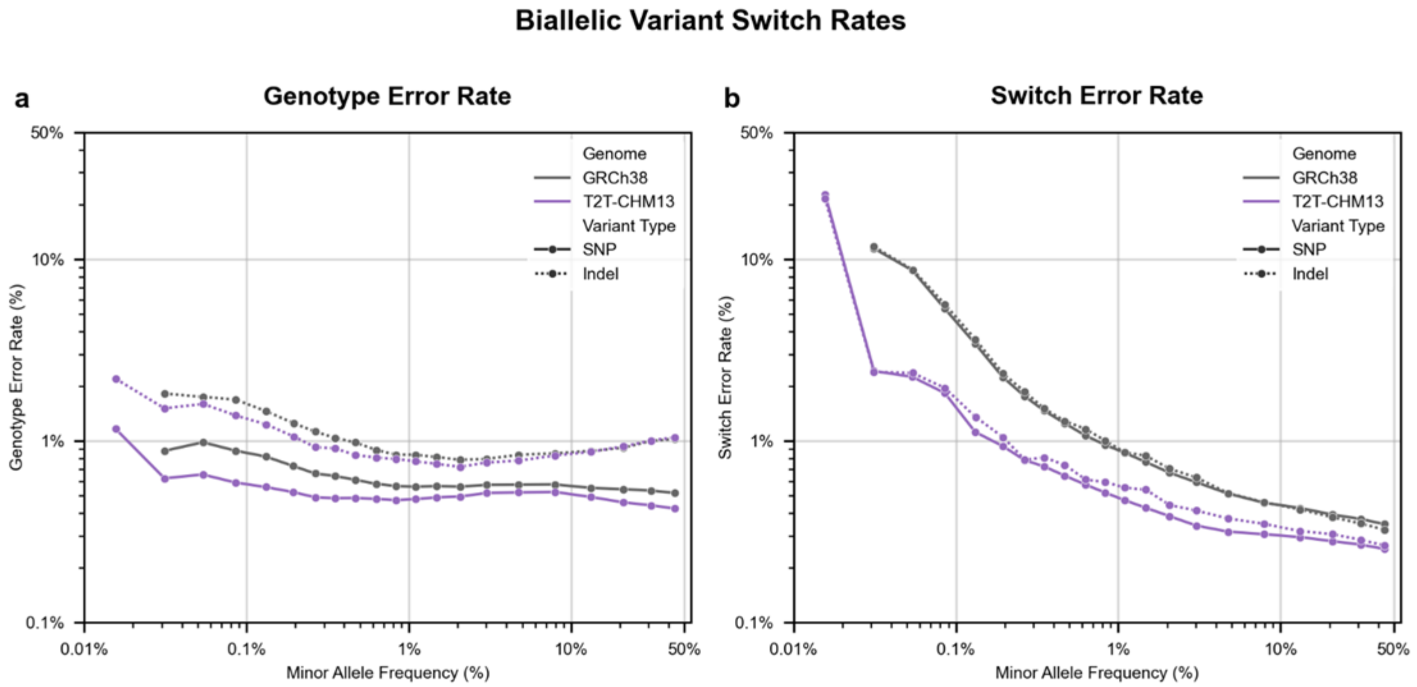


**Supplemental Figure 14. Genotyping and switch error rates for isolated variants in GRCh38 and T2T-CHM13 panels.**

Estimated panel-wide error rates for isolated, biallelic variants that do not overlap another variant in the panel. Error rates are stratified by variant type (SNPs: solid lines; indels: dashed lines), reference genome (GRCh38: gray; T2T-CHM13: purple), and minor allele frequency (MAF) bin. a) Genotype error rates were calculated as the proportion of panel heterozygous or homozygous alternative genotypes that were discordant with assembly-derived ground truth genotypes. Estimated panel-wide error rates were calculated for trio probands, trio parents, and non-trio samples. A weighted average per-bin panel genotype error was then calculated. b) Switch error rates were calculated as the proportion of heterozygous variants incorrectly phased with respect to the nearest 3' heterozygous variant. Estimated panel-wide error rates were calculated as in panel a. Analysis based on 100 samples present in both the 1KGP dataset and the HPRCv1.1/HGSVC joint pangenome.

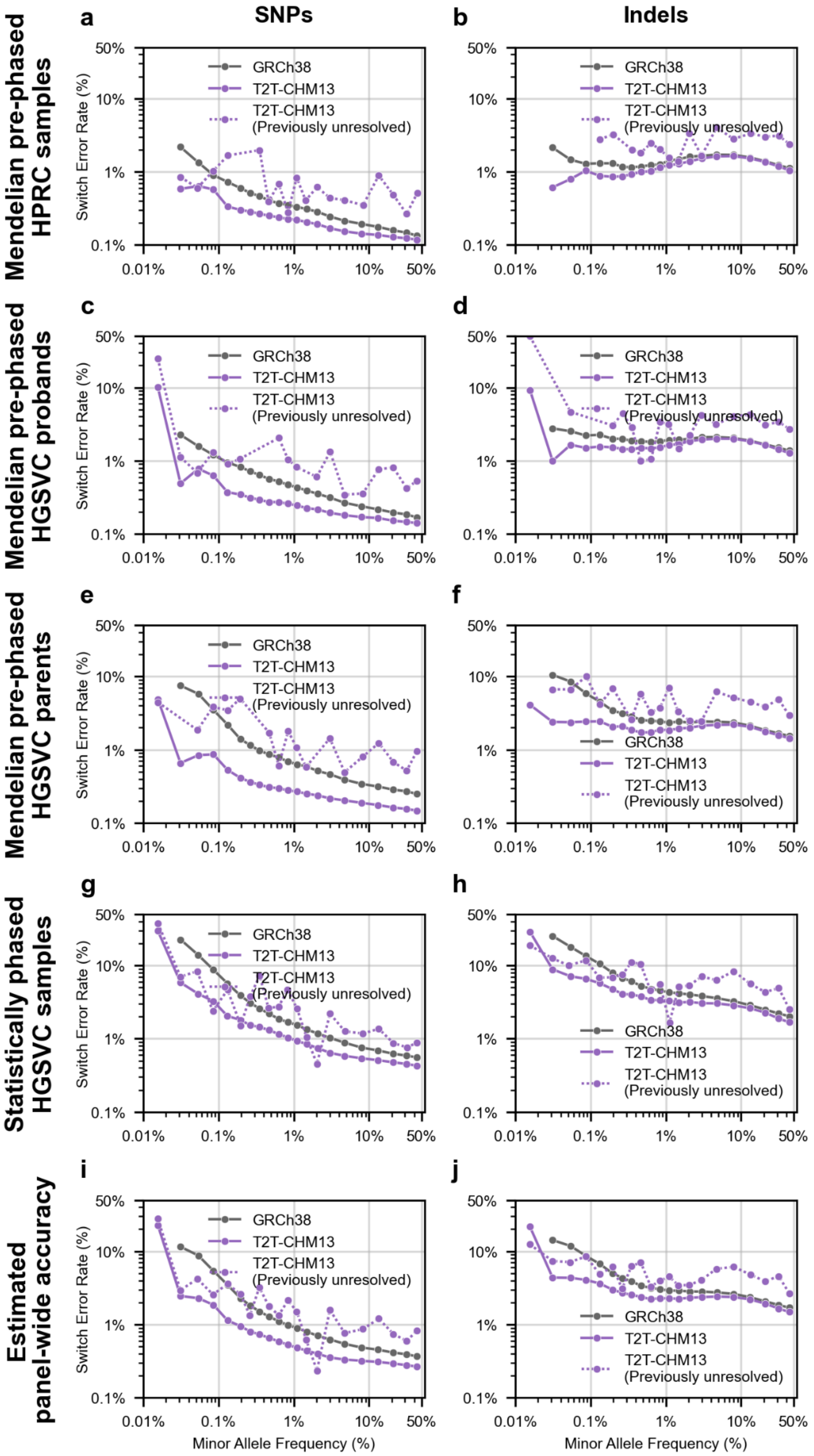


**Supplemental Figure 15. SNP (L column) and Indel (R column) switch error rates for 1KGP phased haplotype panel variation, binned by minor allele frequency.**

Switch error rates were determined by comparing estimated phasing to ground-truth assemblies for the following subsets of samples: a) Mendelian-corrected trio probands with ground truth assemblies present in the HPRC human pangenome; b) Mendelian-corrected trio probands with ground truth assemblies produced by the HGSVC; c) Mendelian-corrected trio parents with ground truth assemblies produced by the HGSVC; d) Uncorrected samples which are not part of a 1KGP trio with ground truth assemblies produced by the HGSVC. e) The switch error rates of trio probands, trio parents, and non-trio samples were weighted by each category's prevalence in the 1000 Genomes Project to produce estimated panel-wide average switch error rates. Switch error rates and average minor allele frequency per bin are displayed on a log scale.

a

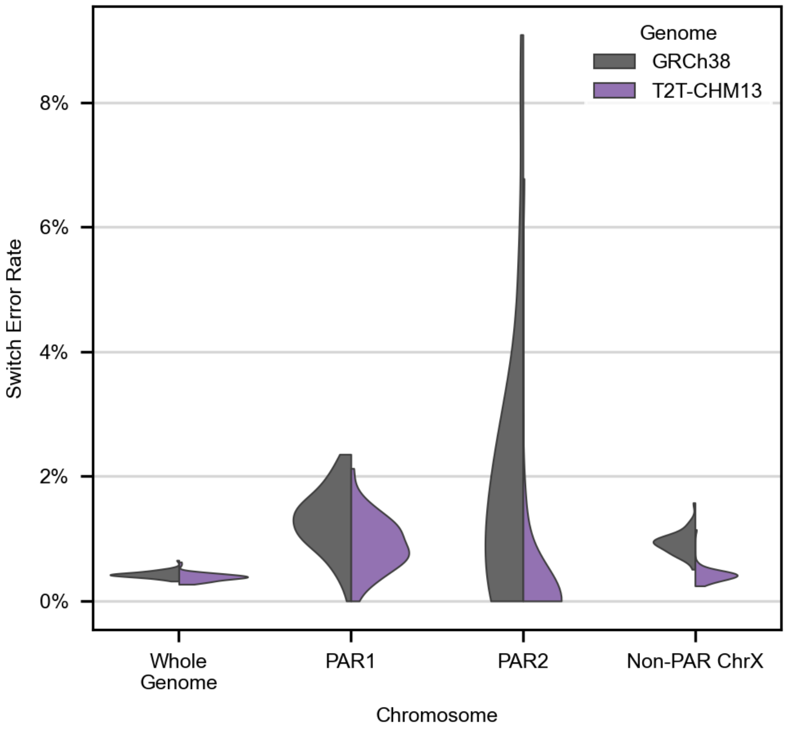


**Supplemental Figure 16. Whole genome, pseudoautosomal region 1 (PAR1), PAR2, and non-PAR chromosome X switch error rates.**

Switch error rates for trio probands with HPRC or HGSVC assemblies from either the GRCh38 or T2T-CHM13 1KGP reference haplotype panels, with each sample's assembly used as ground truth. PAR switch error rates are calculated separately from the non-PAR region of chromosome X. Non-PAR chromosome X switch error rates are only calculated from female samples.

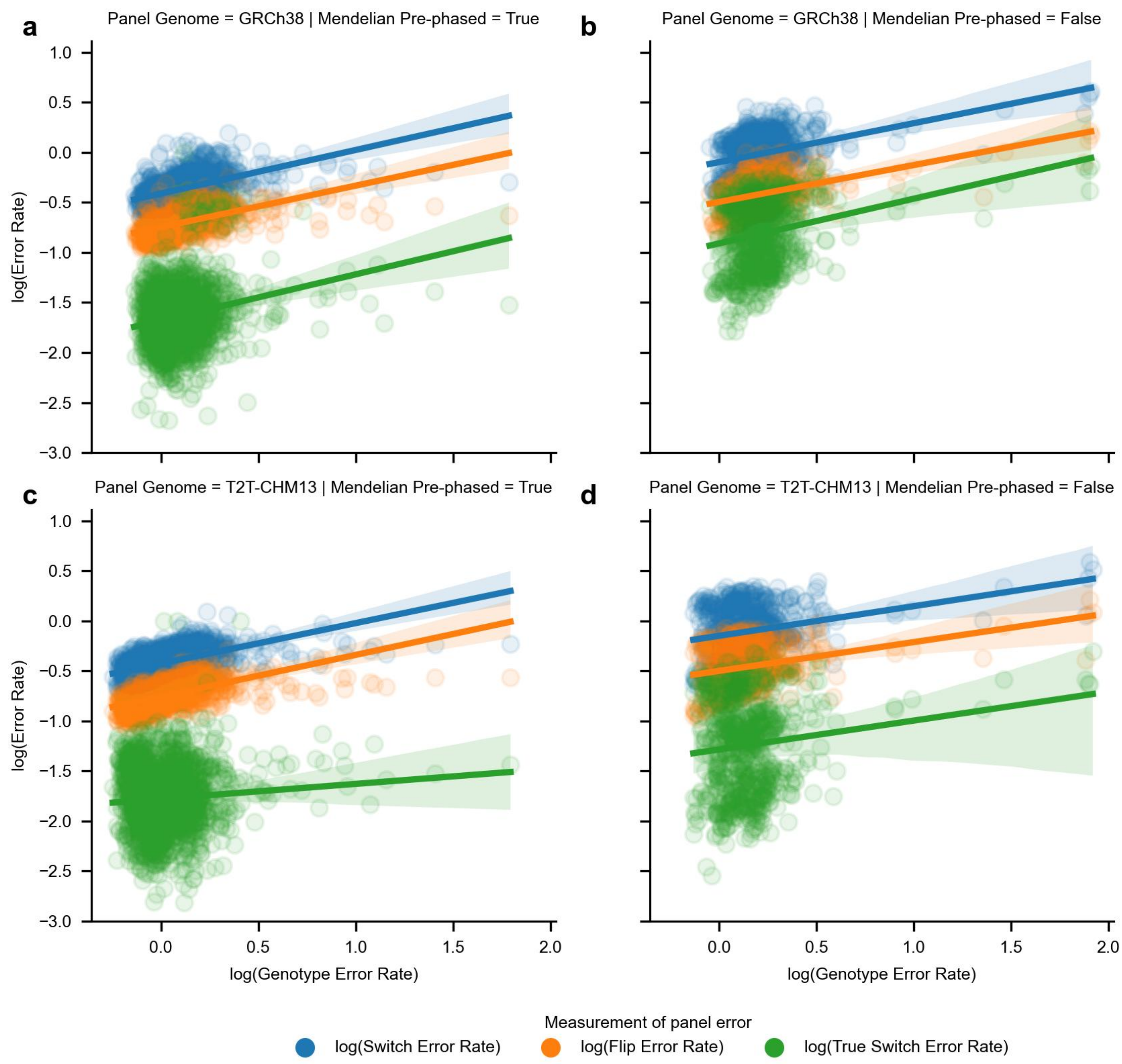


**Supplemental Figure 17. Association between flip error rate, switch error rate, true switch error rate, and genotyping error rate.**

The flip error rate, switch error rate, and true switch error rate was calculated for all chromosomes and all samples for which assembly-based ground truths existed. Genotype error rates were calculated for each chromosome in each sample. Log10 values of each error rate were taken to transform each rate to a gaussian distribution. Linear regression was sequentially performed with the formula log10(Error_Rate) ~ log10(genotype_error_rate), and r2 correlation values were calculated. a) Per chromosome per sample error rates in the GRCh38 haplotype panel for Mendelian error-corrected samples assemblies generated by the HPRC or the HGSVC. b) Per chromosome per sample error rates in the GRCh38 haplotype panel for non-trio samples with HGSVC assemblies that did not undergo mendelian error correction. c) Per chromosome per sample error rates in the T2T-CHM13 haplotype panel for Mendelian error-corrected samples assemblies generated by the HPRC or the HGSVC. d) Per chromosome per sample error rates in the T2T-CHM13 haplotype panel for non-trio samples with HGSVC assemblies that did not undergo mendelian error correction. All regression lines are plotted with a +/- 95% confidence interval.

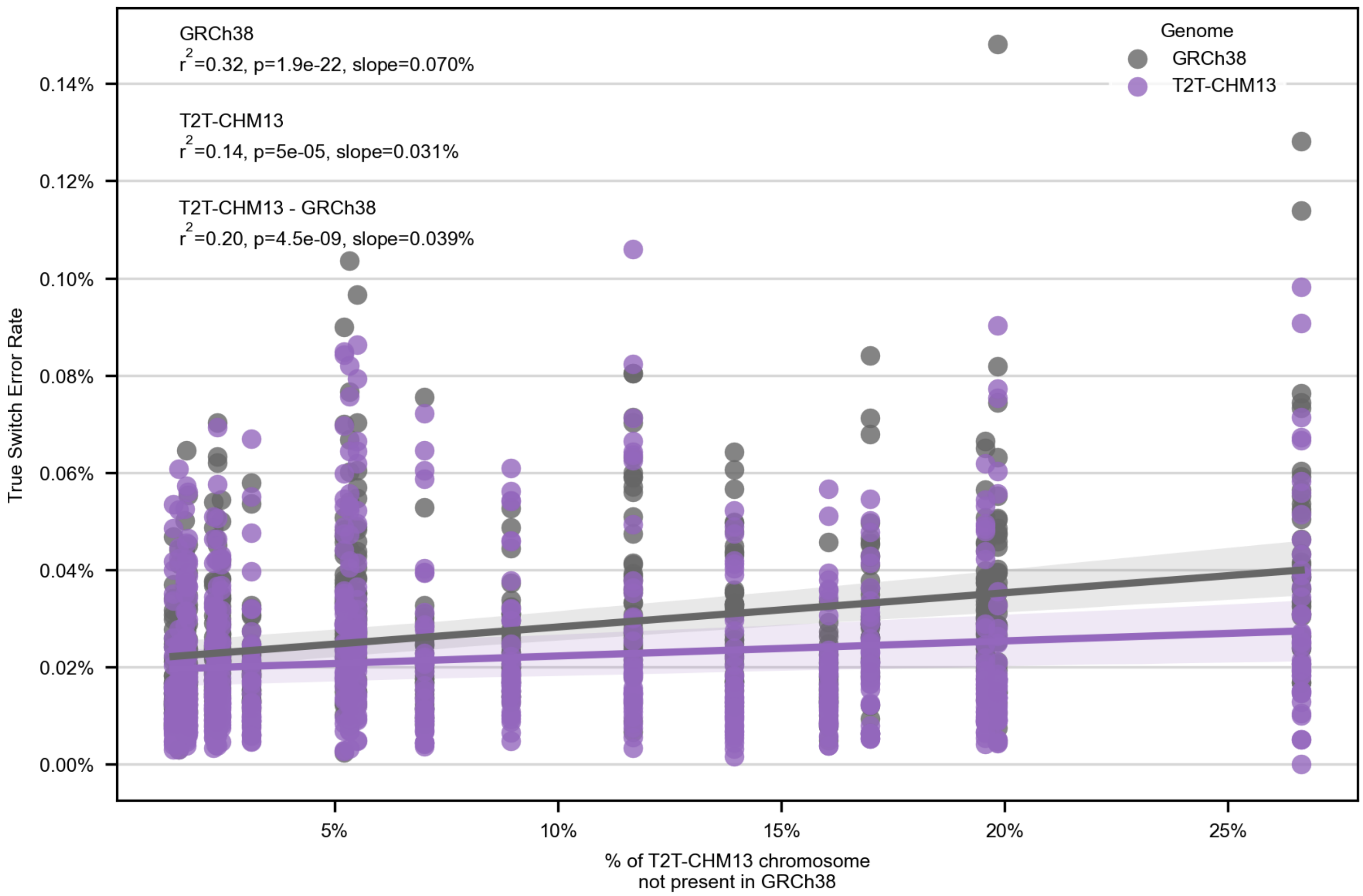


**Supplemental Figure 18. True switch error rate as a function of GRCh38 chromosomal completeness.**

Scatterplot of all chromosome-wide switch error rate in the GRCh38 and T2T-CHM13 1KGP panels for all chromosomes from HPRC-assembled samples by the percentage of each chromosome that is newly resolved in T2T-CHM13. A linear regression (true_switch_error_rate ~ %_chrom_novel), run separately for each 1KGP panel, is also plotted. Shaded areas indicate 95% ci of the regression slope. A linear regression was also performed of ΔSER% ~ %_chrom_novel. Summary statistics of all three regressions are presented in the upper left corner of the plot.

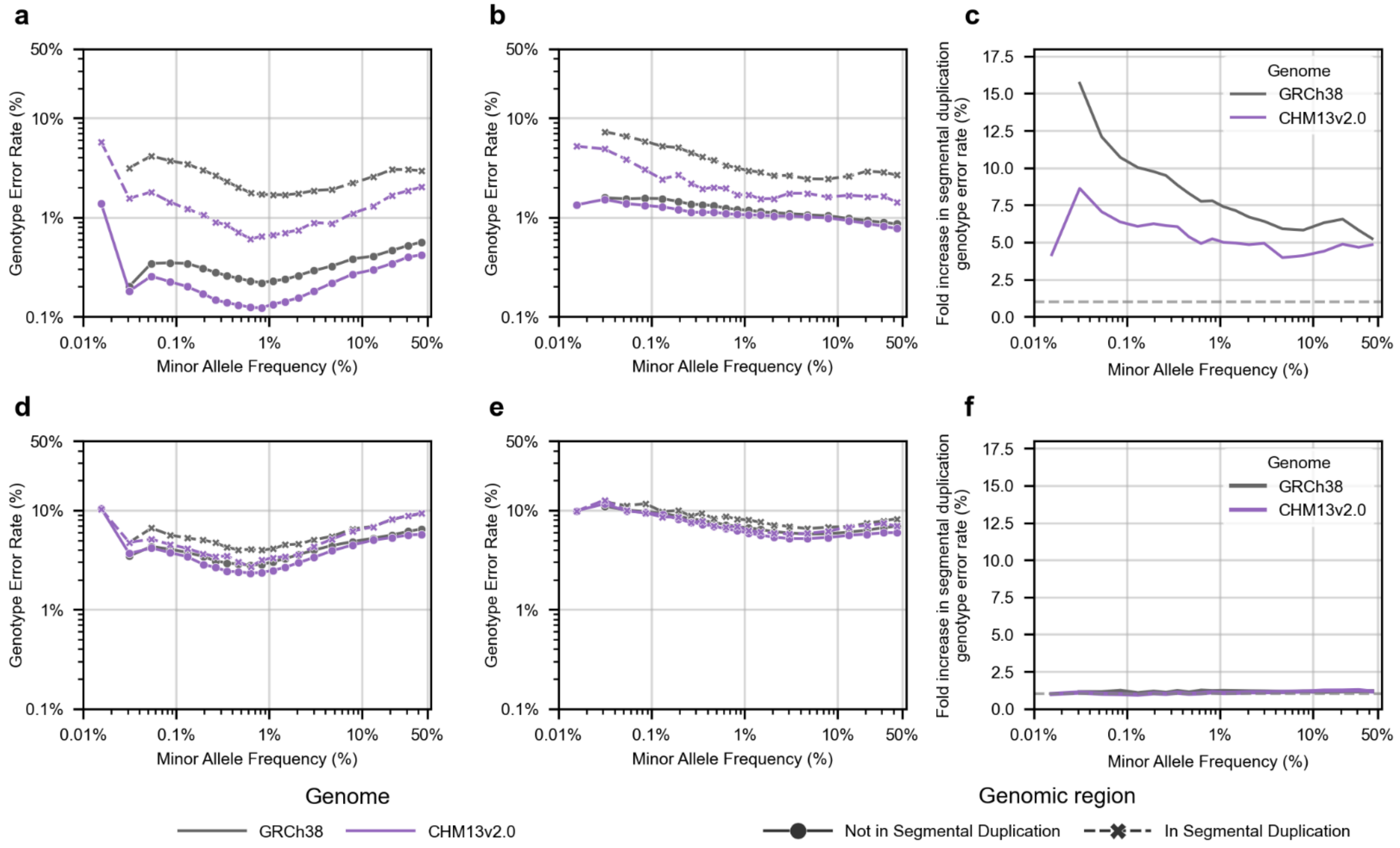


**Supplemental Figure 19: Genotype error rates of 1KGP variants located in segmental duplications (SDs) annotated in both GRCh38 and T2T-CHM13**

Variants from the Bykstrom-Bishop (2022) GRCh38 reference haplotype panel and this paper's T2T-CHM13 panel were stratified by whether they overlapped segmental duplication regions as defined in Vollger (2022). They were additionally binned into 21 distinct minor allele frequency bins. Genotyping error rates of each bin were calculated as # of genotyping errors/# alt allele genotype calls in that bin. a) Genotyping error rates of SNPs from 39 trio-corrected HPRC samples, with HPRC genomes used as a source of ground truth. b) Genotyping error rates of SNPs from 19 1KGP samples that were not trio-corrected, with assembled HGSVC genomes used as a source of ground truth. c) Fold increase in the genotype error rate of SD SNPs over non-SD SNPs by MAF bin, stratified by panel. d) Genotyping error rates of indels from the same 39 HPRC samples as in figure a. e) Genotyping error rates of indels from the same 19 HGSVC samples as in panel b. f) Fold increase in indel genotyping error rate (SD/non-SD).

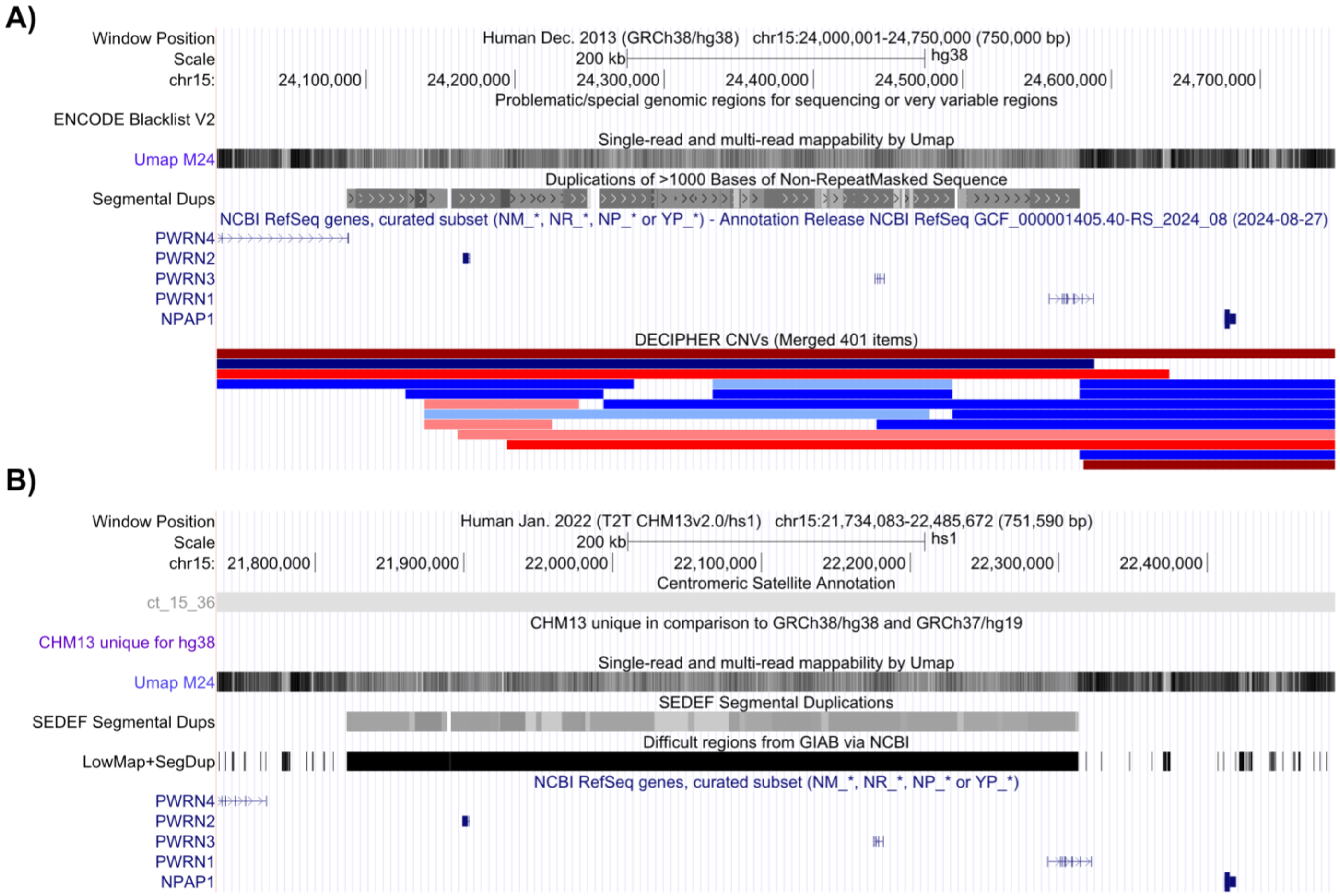


**Supplemental Figure 20. UCSC Genome Browser information on mappability of a region near the Prader-Willi breakpoint 2 (roughly chr15:24100000-24600000) that is enriched for genetic variants in the GRCh38 phased genomic panel.**

a) Mappability, blacklist status, location of segmental duplications, and genic regions in GRCh38. b) Mappability, location of segmental duplications, “difficult region” status from GIAB, and genic regions in T2T-CHM13.

a

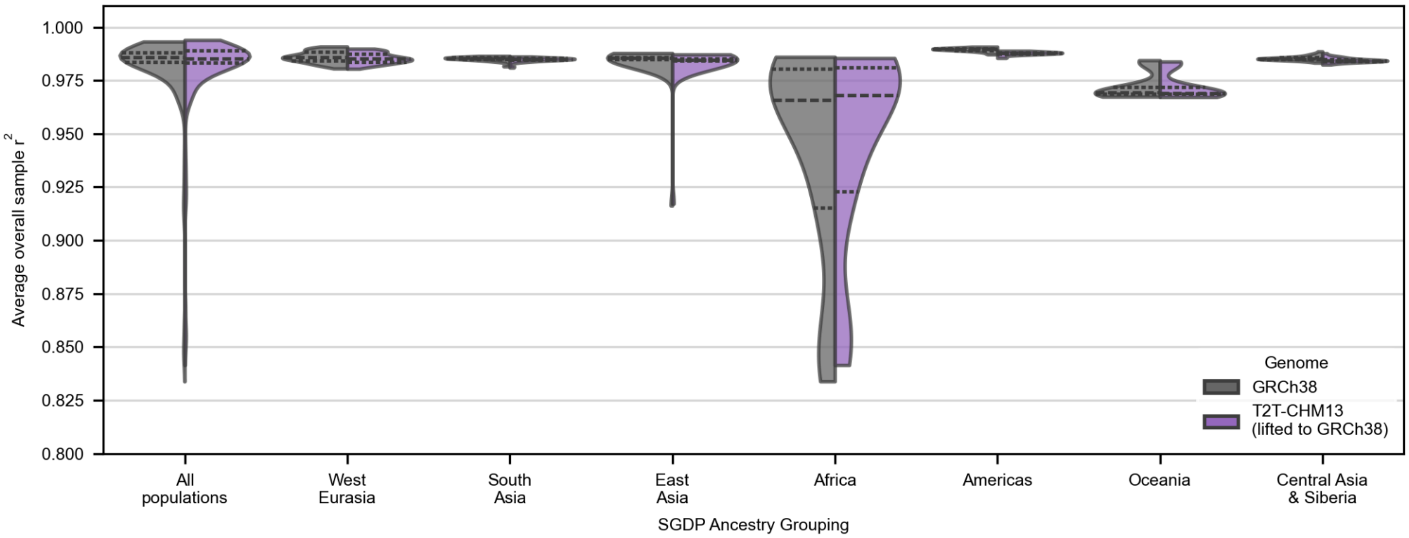


**Supplemental Figure 21. A violin plot displaying the distribution of overall per-sample r2 values when imputing GRCh38 SGDP variation, stratified by SGDP-defined sample continental groups.**

Distributions are split by reference panel used during imputation: either the Byrska-Bishop 2022 GRCh38 reference haplotype panel, or the T2T-CHM13 panel lifted over to GRCh38 coordinates.

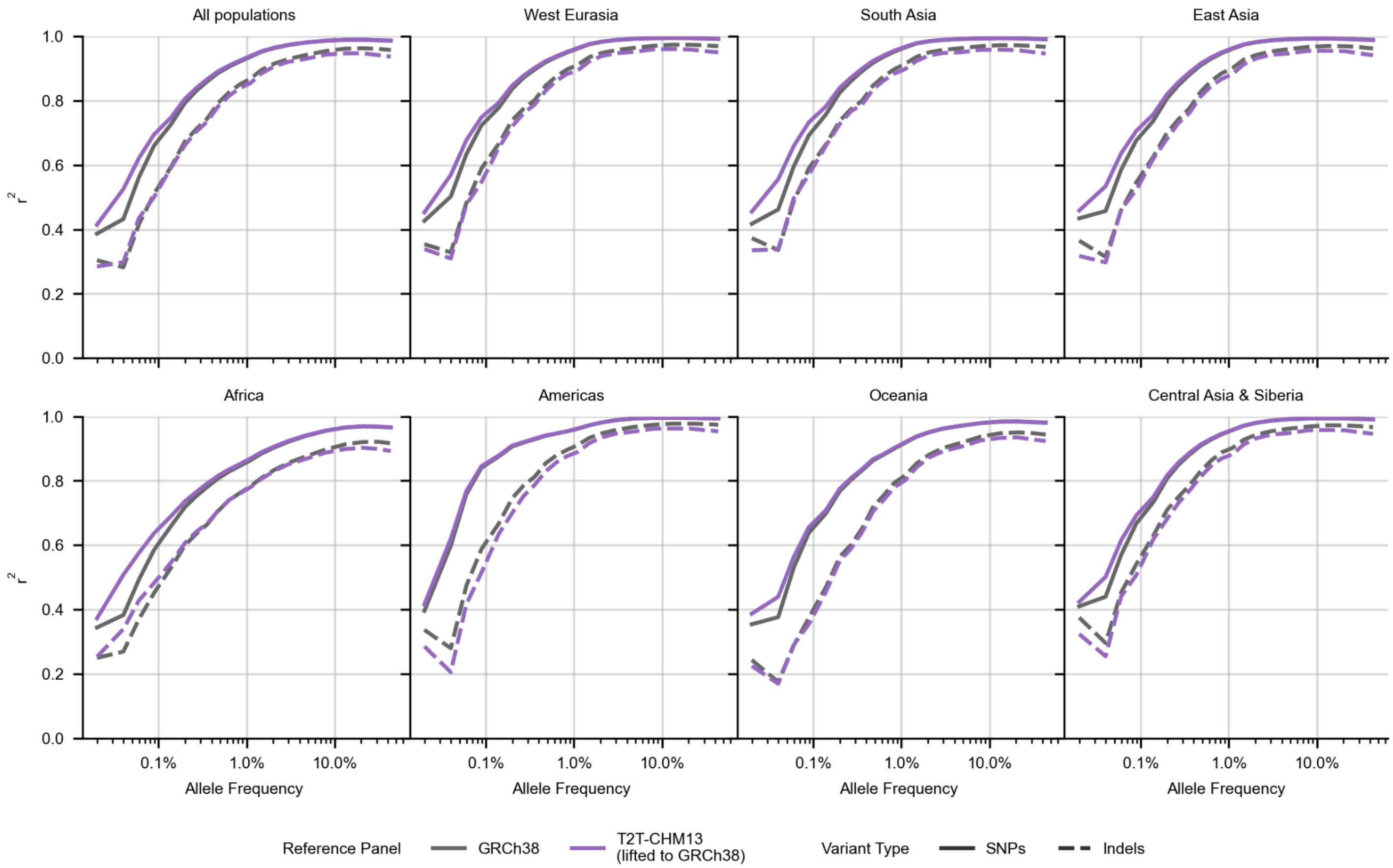


**Supplemental Figure 22: Imputation of 256 non-1KGP HGDP GRCh38 genetic variation using 1KGP haplotype panels as references, stratified by SGDP population of origin.**

As previously presented, genotyping array data was simulated by downsampling variation derived from short reads aligned to GRCh38 were downsampled to those variants present in the Infinium Omni2.5 genotyping array. Variants present in our callset and in the genotyping array were then phased and imputed using either the GRCh38 1KGP haplotype panel, or the T2T-CHM13 1KGP panel lifted to GRCh38 coordinates. $r^2$ values were calculated from the intersection of variants that were in both callsets. $r^2$ statistics were calculated from imputed variants in samples assigned by the SGDP to the indicated ancestry group. $r^2$ values were separately calculated in SNP variation and Indel variation. Average minor allele frequency per bin is displayed on a log scale.

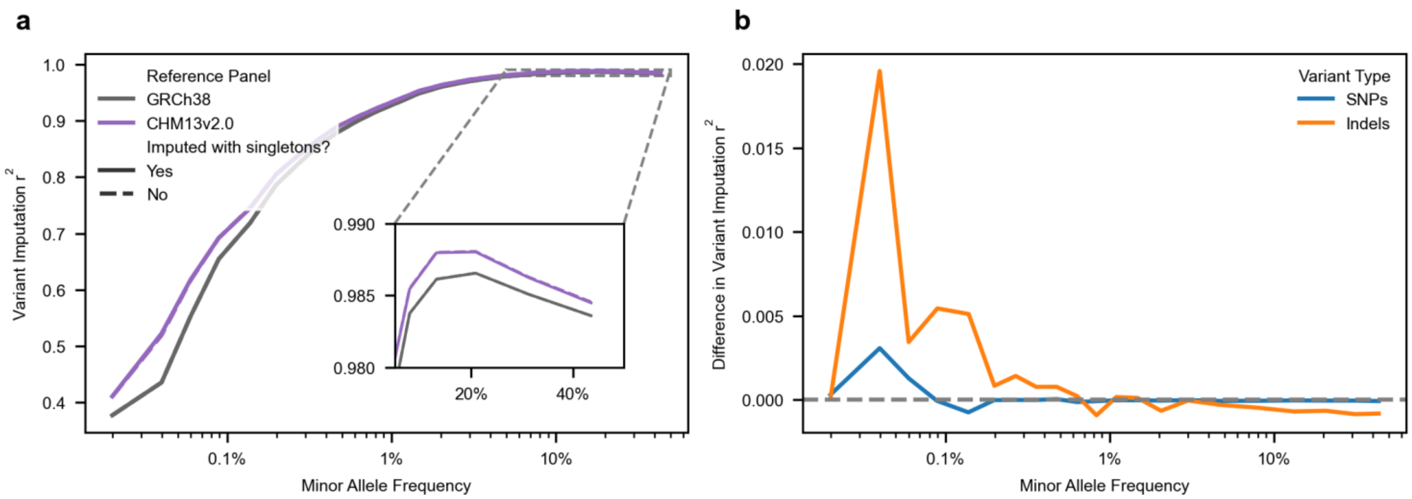


**Supplemental Figure 23: Impact of including singleton variants in reference panels on imputation accuracy.**

A) SGDP GRCh38 variants downsampled to OMNI2.5 sites were imputed using 1KGP reference haplotypes constructed in either GRCh38 or T2T-CHM13 coordinates (the latter lifted to GRCh38). Reference panels were processed identically and included either all variants or excluded singletons (minor allele count = 1); related samples were removed, yielding 2,504 unrelated individuals per panel. Imputation accuracy was measured as genotype dosage $r^2$ against whole-genome sequencing truth and summarized across 21 bins of reference panel minor allele frequency. B) Difference in either SNP or Indel MAF bin imputation accuracy ($\Delta r^2$) between GRCh38 variants imputed using lifted T2T-CHM13 reference panels that include versus exclude singleton variants. Positive values indicate improved accuracy when singletons are included in the reference haplotype panel during imputation.

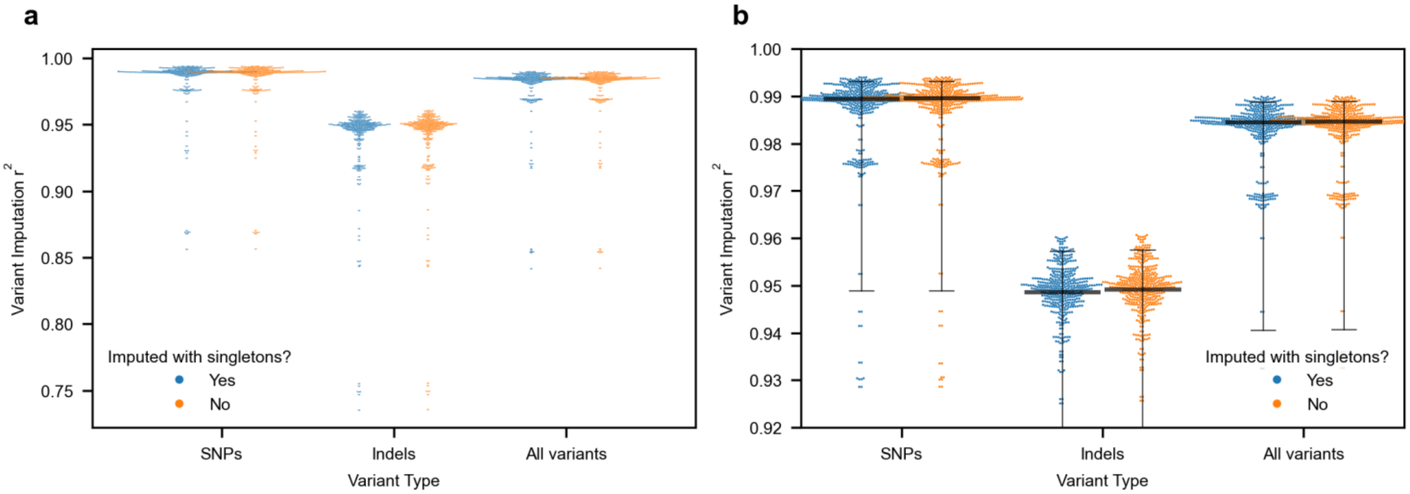


**Supplementary Figure 24: Per-sample impact of singleton inclusion on imputation accuracy when using T2T-CHM13 reference panels**

A) Per-sample imputation accuracy ($r^2$) for SNPs, indels, and all variants, comparing reference panels that include versus exclude singleton variants. Each point represents a single SGDP sample; distributions summarize sample-level variability in imputation performance. B) A duplicate of panel A, showing only $r^2$ values of 0.92 or higher to highlight small but consistent differences in per-sample imputation accuracy. The wide central marker indicates the median $r^2$ across samples, with whiskers denoting the 5th and 95th percentiles.

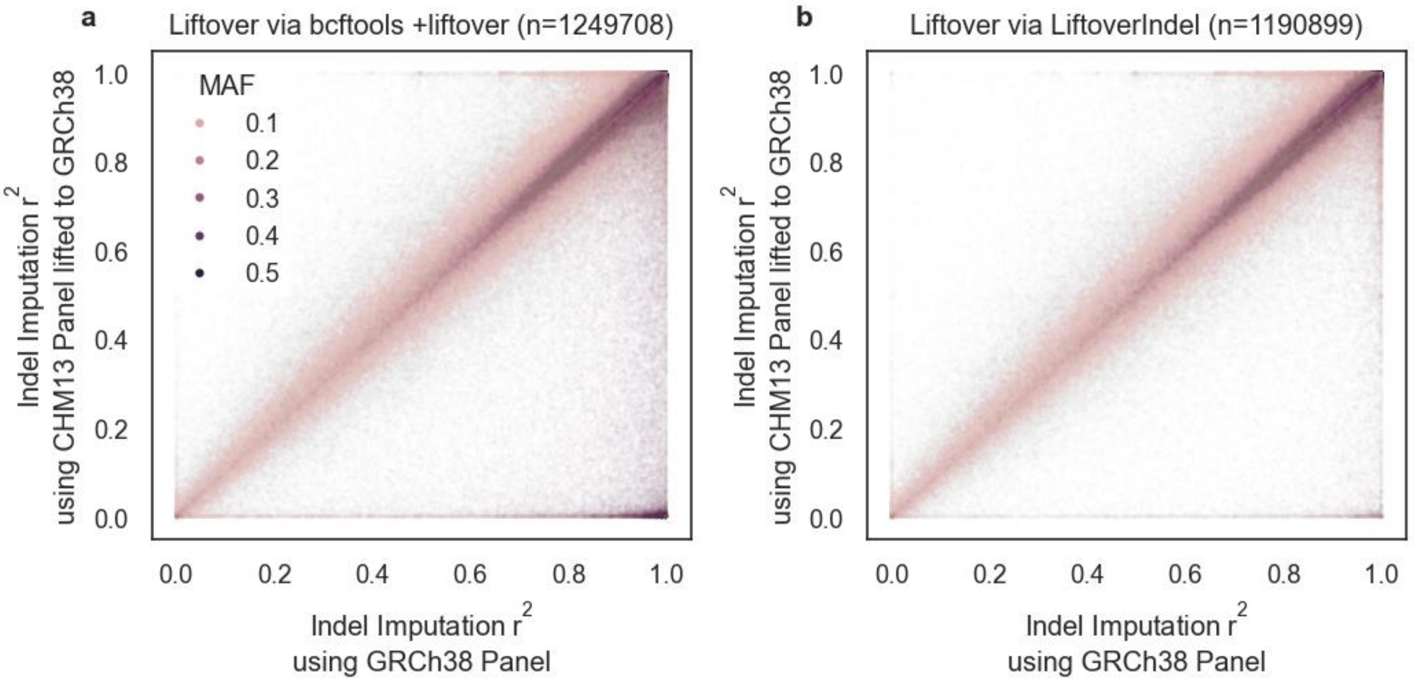


**Supplemental Figure 25. Improved liftover algorithm results in better imputation of common indel variants.**

a) Scatter plot of the $r^2$ value of imputed indel variants over 10% MAF when using a GRCh38 1KGP reference panel (x-axis) or the T2T-CHM13 panel lifted to GRCh38 coordinates using either a) GATK LiftoverVCF or b) the modified liftover algorithm implemented in this paper. Manual inspection of the high MAF variants that were poorly called by the lifted-over T2T-CHM13 panel showed that the majority of these variants are short tandem repeats that are lifted to a site that is several bases away from the same variant in the 1KGP GRCh38-native variant callset. Each dot represents one biallelic indel. The panel minor allele frequency is colored by 1KGP GRCh38 MAF.

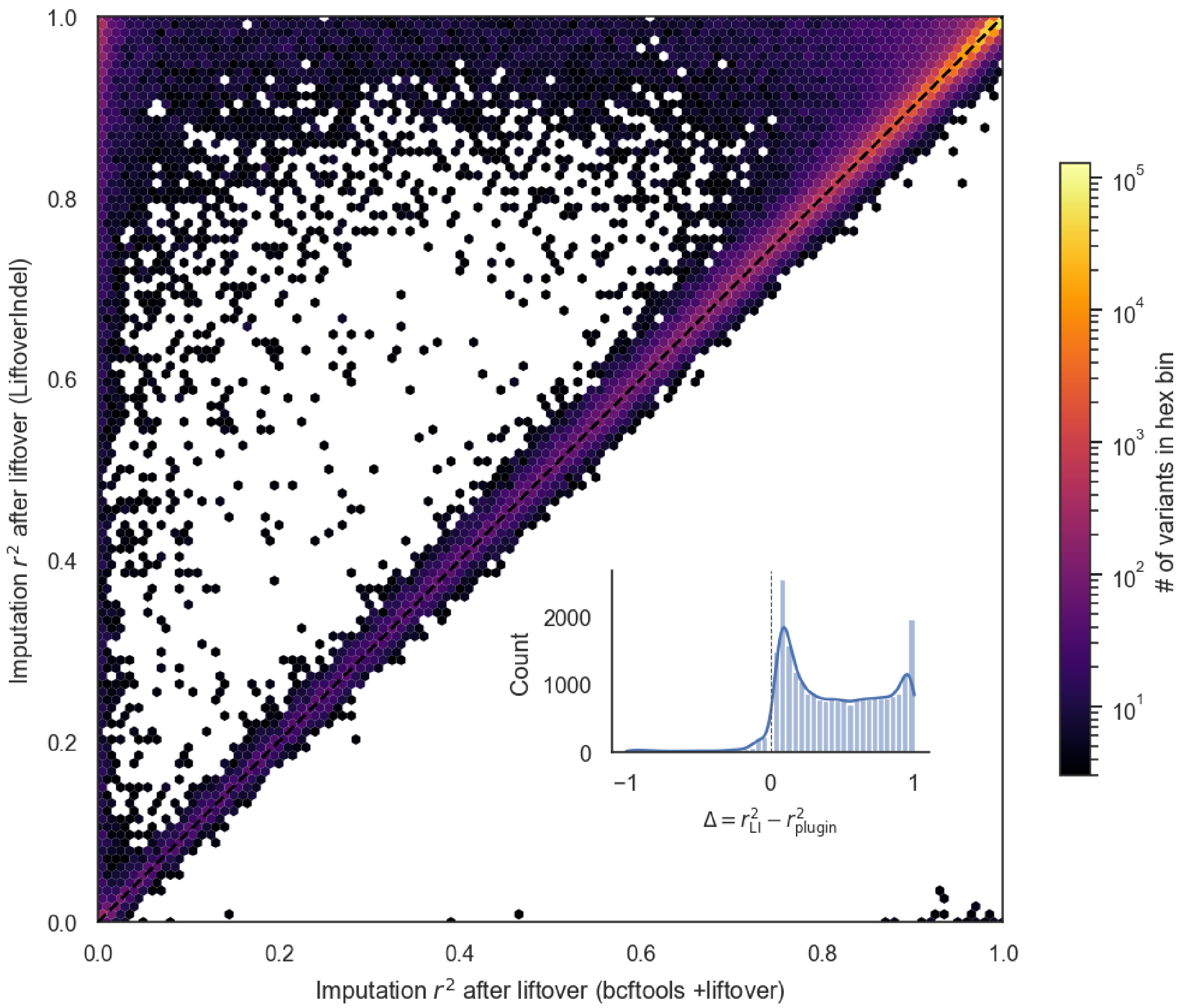


**Supplemental Figure 26: Distribution of $r^2$ values of imputed common indels (>10% MAF) when imputed using a reference panel lifted to GRCh38 coordinates with bcftools +liftover or LiftoverIndel.**

Each hexagonal bin represents the density of variants with a given pair of imputation $r^2$ values (log-scaled color bar). Only bins with 3 or more variants are shown. The dashed diagonal denotes equality between methods. Points above the diagonal indicate higher imputation accuracy with LiftoverIndel. Inset*:* For each variant, we calculated the difference in imputed $r^2$ value when using LiftoverIndel or Bcftools +liftover. Higher values indicate more accurate imputation when using a LiftoverIndel panel. The distribution of values for variants with an absolute difference over 0.05 are shown.

Supplemental Table 1

| Panel | Ground Truth | Trio Status | Switch Error Rate (%) | Flip Error Rate (%) | True Switch Error Rate (%) | Genotype Discordance Rate (%) |
|---|---|---|---|---|---|---|
| GRCh38 | HPRC | Proband | 0.408 | 0.186 | 0.029 | 1.114 |
| GRCh38 | HGSVC | Proband | 0.504 | 0.232 | 0.030 | 1.333 |
| GRCh38 | HGSVC | Parent | 0.705 | 0.303 | 0.080 | 1.457 |
| GRCh38 | HGSVC | Non-trio | 1.285 | 0.487 | 0.257 | 1.443 |
| T2T-CHM13 | HPRC | Proband | 0.355 | 0.166 | 0.019 | 0.918 |
| T2T-CHM13 | HGSVC | Proband | 0.428 | 0.202 | 0.019 | 1.153 |
| T2T-CHM13 | HGSVC | Parent | 0.495 | 0.235 | 0.020 | 1.277 |
| T2T-CHM13 | HGSVC | Non-Trio | 1.110 | 0.474 | 0.130 | 1.253 |

Error rates, stratified by ground truth source and whether samples are trio parents, trio probands, or not part of a trio (aka statistically phased).
